# Oral Lichen Planus and its relation with Oral Squamous Cell Carcinoma: new insights into the potential for malignant transformation

**DOI:** 10.1101/2021.10.29.466364

**Authors:** Cristóvão Antunes de Lanna, Beatriz Nascimento Monteiro da Silva, Andreia Cristina de Melo, Martín H. Bonamino, Lísia Daltro Borges Alves, Luis Felipe Ribeiro Pinto, Abel Silveira Cardoso, Héliton Spíndola Antunes, Mariana Boroni, Daniel Cohen Goldemberg

## Abstract

Oral Lichen Planus (OLP) is a chronic inflammatory disorder of unknown etiology. However, evidence suggests that it consists of an immunological process that leads to degeneration of the keratinocytes in the basal layer of the oral mucosa. Despite being recognized by WHO as a potentially malignant disorder with progression to oral squamous cell carcinoma (OSCC), the relationship between both pathologies is still controversial. Different studies have investigated factors associated with the potential for malignant transformation of OLP but it remains unclear. Through a bioinformatics approach, we investigated similarities in gene expression profiles of OLP and OSCC in early and advanced stages. Our results revealed gene expression patterns related to processes of keratinization, keratinocyte differentiation, cell proliferation and immune response in common between OLP and early and advanced OSCC, with the cornified envelope formation and antigen processing cross-presentation pathways in common between OLP and early OSCC. Together, these results reveal that key genes such as *PI3, SPRR1B* and *KRT17*, in addition to genes associated with different immune processes such as *CXCL-13, HIF1A* and *IL1B* may be involved in this oncogenic process. In addition, we performed an analysis of differentially and co-expressed genes and proposed putative therapeutic targets and associated drugs.

## Introduction

Oral Lichen Planus (OLP) is a chronic inflammatory disease clinically characterized by six distinct subtypes that can be seen individually or in combination: white reticular striations, papular, plaque-like, erythematous erosions, ulcerative, and bullous forms. Of all the presentations, the reticular form is the most common, exhibiting a delicate white banding, called Wickham’s striae (Kurago, 2016; Warnakulasuriya et al., 2007; Wickham, 1895). Histologically, OLP is characterized by vacuolar degeneration, a band-like dense inflammatory infiltrate of T lymphocytes at the epithelial-stromal junction, and hyperkeratosis or parakeratosis (Cheng et al., 2016).

The origins of this persistent cytotoxic T-cell-mediated damage are currently unknown. However, many authors suggest that the disease is associated with an autoimmune process (Farhi and Dupin, 2010; Ismail et al., 2007; Roopashree et al., 2010; Rutz et al., 2016). In 1910, Hallopeau and colleagues described for the first time a case of oral squamous cell carcinoma (OSCC), a malignant neoplasm that originates in the lining epithelium and is considered the most common malignancy in this region, in a patient with OLP. Since then, different studies have suggested a premalignant potential for OLP injuries over the years (Aghbari et al., 2017; Bardellini et al., 2013; Barnard et al., 1993; Hallopeau, 1910).

The World Health Organization (WHO) defined in 2017 that OLP is an *oral potentially malignant disorder* (OPMD), with a possible progression to OSCC (Müller, 2017; Peng et al., 2017; van der Waal, 2010). Giuliani and colleagues (2019), in a systematic review, demonstrated that 92 of 6559 patients diagnosed with OLP developed OSCC. The authors found a malignancy potential of 1.4% and an annual transformation rate of 0.2% when analyzing retrospective and prospective studies on OLP. Ruokonen et al. (2017) revealed that 17.9% of patients with oral cancer had OLP and 4% had oral lichenoid lesions, suggesting that this dysfunction may be an important etiological factor of OSCC and that smoking and alcohol use was less frequent in patients with OLP and lichenoid lesions, suggesting that OLP may progress to OSCC, even in the absence of well-known etiological agents.

In this study, we investigated gene expression signatures observed in OLP lesions when compared to healthy oral tissues that are also disturbed in the early and advanced stages of OSCC in order to shed light on the potential for malignant transformation of OLP. We demonstrate a repertoire of pathways that could be connected to the potential transition between OLP and OSCC. Our findings suggest that OLP and OSCC share the activation of keratinization- and inflammation-related pathways with a potential role in malignization. We have also explored potential novel therapeutic targets and propose drugs that can interact with them and revert expression alterations.

## Results

### Gene expression profiles among OLP and OSCCs

Gene expression data (16,656 features) comprising 13 normal oral mucosa, 7 OLP, 10 early-stage OSCC, and 20 advanced-stage OSCC samples were integrated and compared. Of note, although OLP samples tend to group closer to normal samples, at least one sample shows high similarity with OSCC samples (Fig. S1).

Differential expression analysis was performed for each group in comparison to normal tissue. A total of 107 DEGs were identified for OLP (83 overexpressed, 24 underexpressed), 331 for early-stage OSCC (182 overexpressed, 149 underexpressed), and 282 for advanced OSCC (96 overexpressed, 186 underexpressed) (Tables S1-S3; Fig. S2A-C). Gene set enrichment analysis (GSEA) was performed for each group. Enriched pathways in OLP were related to keratinization, extracellular matrix (ECM) and its interaction with surrounding tissue, and immunity-related pathways such as complement, interleukin 10 (IL-10) signaling, antimicrobial peptides, antigen presentation, among others (Table S5). In early and advanced OSCC, enriched pathways included pathways related to keratinization, DNA replication, RNA transcription, DNA repair, cell cycle, multiple interleukin and interferon signaling pathways, among others (Tables S6-S7). To better explore OLP’s potential for malignization and its relation with OSCC, the overlapping differentially expressed genes (DEGs) between OLP and OSCC were also investigated, resulting in a total of 35 overlapping DEGs (29 Up- and 6 Down-regulated). Fifteen genes were consistently differentially expressed in all comparisons (10 overexpressed, 5 underexpressed) (Fig. 1A-B; Table S4). Most of the overlapped genes (51.4%) occurred between OLP and early stage OSCC. When using a non-supervised clustering approach based on the expression of the 35 gene signature, all OLP samples were clustered with OSCC samples, mainly in the early stage (Fig. 1C). We also observed overlap among pathways enriched in the three conditions. The enrichment analysis revealed that OLP has six main pathways in common with early or advanced OSCC: antigen processing cross-presentation; formation of the cornified envelope; interleukin-10 signaling; collagen chain trimerization; non-integrin membrane-ECM interactions; and neutrophil degranulation, with the antigen presentation pathway enriched in all conditions. Interestingly, antigen presentation and formation of the cornified envelope were the only common pathways between early OSCC and OLP, with the latter being common only between these two groups (Fig. 1D). Some of the genes influencing the clusterization are those coding for keratins, including the downregulation of *KRT4* and up-regulation of *KRT16, KRT17, KRT10*, and *KRT75* in comparison to normal samples (Fig. 1E).

**Figure 1.**
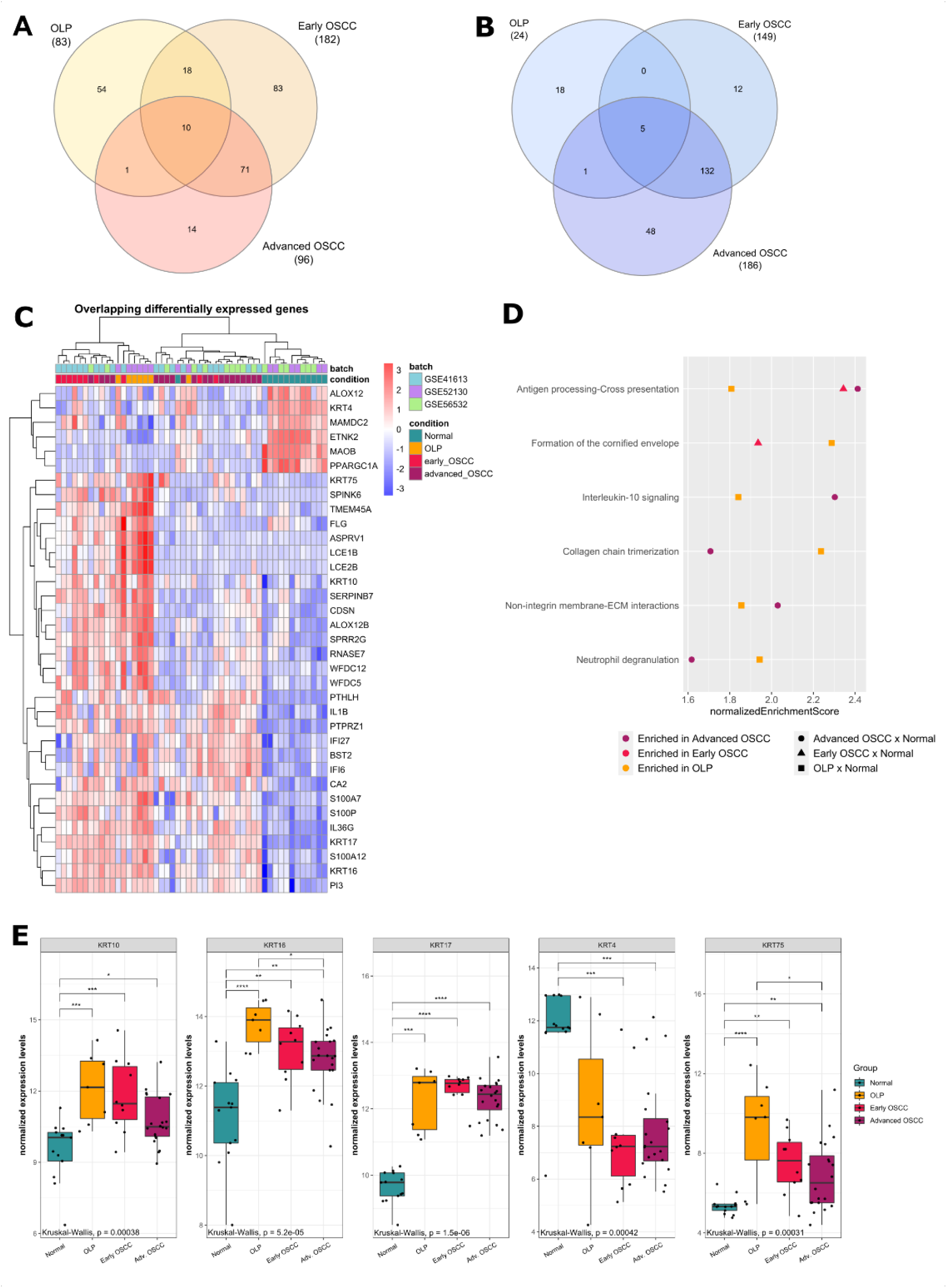
Differentially Expressed Genes between OLP, early, and advanced OSCC microarray datasets compared to normal oral tissue. (A) Venn diagram of overlapping DEGs of overexpressed genes. Ten of the differentially expressed genes were overexpressed in all three groups, while 18 overexpressed genes were shared by OLP and early OSCC. (B) Venn diagram of overlapping DEGs of underexpressed genes. Five of the differentially expressed genes were underexpressed in all three groups. (C) Clustered heatmap of DEGs shared between OLP, early, and advanced OSCC, as well as overexpressed genes shared by OLP and early OSCC. (D) Gene set enrichment analysis (GSEA) of Reactome pathways in Oral Lichen Planus (OLP), early, and advanced Oral Squamous Cell Carcinoma (OSCC). All pathways are upregulated in comparison with normal tissue. For each pathway, orange squares, red triangles, and purple circles correspond to the normalized Enrichment Score for said pathway in OLP, early OSCC, and advanced OSCC, respectively. Only pathways enriched simultaneously in OLP and at least one of the OSCC stages are represented. (E) Differentially expressed keratin genes between OLP and both OSCC groups compared to normal tissue. Box plots represent the normalized expression distribution in each group. Comparisons among groups were made using the Kruskal-Wallis test followed by Dunn’s post-hoc test, with p-values lower than 0.05 considered significant for both tests. *, p < 0.05; **, p < 0.01; ***, p < 0.001; ****, p < 0.0001.

Of note, genes coding for keratinocyte differentiation-related members of the S100 protein family such as *S100A7, S100P*, and *S100A12* as well as immunity-related genes such as *IL1B, IL36G, IFI6*, and *IFI27*, all of them overexpressed in OLP samples compared to normal tissue, were also identified. To validate our findings, independent datasets corresponding to each of the tested groups were used. Similar results were found to the expression of *KRT4* (lower expression) and *KRT75* (higher expression), whereas differences found in *KRT10, KRT16* and *KRT17 expression*, all of them higher than normal in our analysis, have not been validated (Fig. S3A and S3B). Data obtained from TCGA for early and advanced OSCC showed lower *KRT4* expression and higher *KRT17* expression when compared with normal samples, also consistent with our analysis (Fig S3C).

Overrepresentation analysis (ORA) was performed for all overlapping DEGs between OLP and at least one OSCC stage (29 up and 6 down-regulated, Fig 1A and 1B). Pathways enriched in overlapping up-regulated genes included keratinization and formation of the cornified envelope (Fig. S4A). These pathways were also among those enriched in OLP-exclusive overexpressed DEGs, which also included immunity-related pathways, such as interferon alpha and beta and IL-1 family signaling (Fig. S4B). In OLP-exclusive underexpressed DEGs, the keratinization and formation of the cornified envelope pathways were also enriched, along with cell junction- and compound metabolization-related pathways (Fig. S4C). When considering overlapping underexpressed DEGs, no pathway was significantly enriched.

### Immune Microenvironment Composition of OLP and OSCC

Given that OLP is a disease with a significant participation of the immune system and both immune evasion and tumor-promoting inflammation are hallmarks of cancer, we have sought to identify the most prominent proportions of infiltrating cells in both conditions. We have used a method that estimates the abundances of cell populations by deconvolution of gene expression data.

When clustering samples based on cell’s population composition, OLP samples mostly clustered with normal samples, although some samples showed high similarity with the immune cells composition found in OSCC samples, enriched in plasma cells and memory activated CD4+ T cells (Fig. 2A).

**Figure 2.**
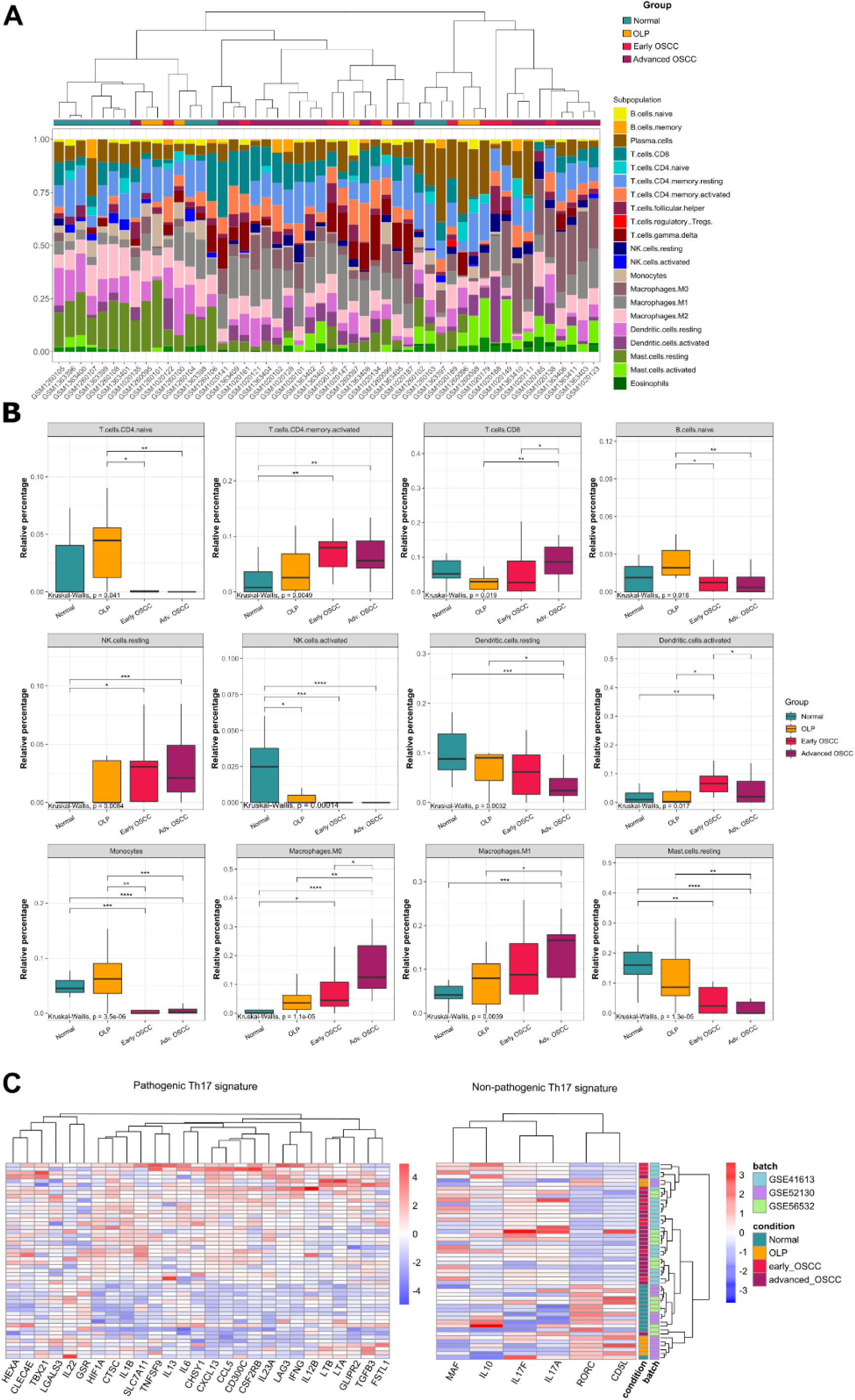
Proportion of cell population signatures identified using CIBERSORTx. (A) Stacked barplot showing cell population proportions in each sample (Oral normal mucosa, OLP, early OSCC, advanced OSCC) based on CIBERSORTx’s 22 cell gene expression signatures. Dendrogram represents Ward clustering of the samples. (B) Relative percentage of each immune cell from oral normal mucosa, OLP, early and advanced OSCC. Comparisons among groups were made using the Kruskal-Wallis test followed by Dunn’s post-hoc test, with p-values lower than 0.05 considered significant for both tests. *, p < 0.05; **, p < 0.01; ***, p < 0.001; ****, p < 0.0001. (C) Clustered heatmap of Th17-related genes. Genes were separated based on non-pathogenic and pathogenic Th17 phenotype-related expression profiles.

The proportions of activated NK cells in OLP samples were significantly lower than the proportions found in the normal oral mucosa. The same was observed for activated NK cells in early or advanced OSCC. The proportions of CD8+ T lymphocytes, M0 and M1 macrophages in advanced OSCC showed significantly higher values when compared to those observed in OLP. The opposite was observed in resting Mast cells, naïve B cells and monocytes. Reduced proportions for all three populations were observed in both early and advanced OSCC when compared to OLP and healthy oral mucosa. The proportions of activated memory CD4+ T cells, resting NK cells, and M0 macrophages were higher in OSCC than in healthy oral mucosa, regardless of staging (Fig. 2B). Considering the immune infiltrate populations in both validation datasets, significantly reduced values of activated NK cells were also observed in OLP, corroborating our results for this group. Similarly, monocyte proportions were consistent with our analysis for early and advanced OSCC in the microarray and TCGA datasets. The resting mast cell proportions were also consistent with those observed in the analysis, while macrophages M0 showed elevated proportions in advanced OSCC in the microarray dataset and early and advanced OSCC in the TCGA dataset, consistently to the proportions in the discovery dataset (Fig. S5).

Since genes that play an important role in the T helper (Th) 17 cell phenotype such as *IL1B, CCL5* and *CXCL13* were differentially expressed in OLP, early and advanced OSCC (Tables S1-S3), we decided to investigate the genes involved in its differentiation given the role of Th17 cells in maintaining mucosal immunity homeostasis. Samples were clustered using a 33-gene panel built based on pathogenic and non-pathogenic Th17 phenotypes characterized in the literature (Lee et al., 2012) (Fig. 2C). When clustering samples with regard to pathogenic and non-pathogenic Th17 signature, two groups can be highlighted: one mostly composed of OSCC samples, showing high expression of many genes from the pathogenic signature and another composed mainly of OLP and normal samples. In this group, genes from the non-pathogenic signature are highly expressed (Fig. 2C). Of note, some genes important in the pathogenic signaling were also found differentially modulated in OLP samples, such as *CTSC, HIF1A, IL1B, LTA, LTB*, and *TGFB3* in relation to normal oral mucosa. Additionally, *CTSC, HIF1A*, and *IL1B* also show higher expression levels in early and late-stage OSCC, with an increasing pattern (Fig. S6A). The validation data also revealed significant high levels of expression of the *CTSC, LTA*, and *TGFB3* genes in OLP (Fig. S6B). Regarding the OSCC groups, however, only *CTSC* and *LTB* exhibited similar expression patterns in comparison to the discovery dataset (Fig S6C-D). *LTA*, while significantly altered compared to normal mucosa in the validation datasets, showed different patterns depending on the validation dataset, with lower expression in the microarray dataset (Fig. S6C) and higher expression in the TCGA validation dataset (Fig. S6D).

### Co-expression analysis

In addition to differential expression analysis, we have also constructed co-expression modules using WGCNA to investigate the connection strength between genes with similar expression patterns and identify potentially co-regulated genes associated with the potential malignization process in OLP. By analysing a total of 15,000 genes (highest gene expression variability among samples), 12 co-expression modules were obtained, each identified by a color: magenta, with 311 genes; purple, with 262 genes; blue, with 5059 genes; greenyellow, with 174 genes; tan, with 142 genes; black, with 381 genes; pink, with 370 genes; red, with 525 genes; turquoise, with 5364 genes; yellow, with 628 genes; brown, with 1070 genes; and green, with 624 genes. A 13^th^ module, to which 90 genes with no co-expression patterns were assigned, was also identified and assigned to the color grey (Fig. 3A).

**Figure 3.**
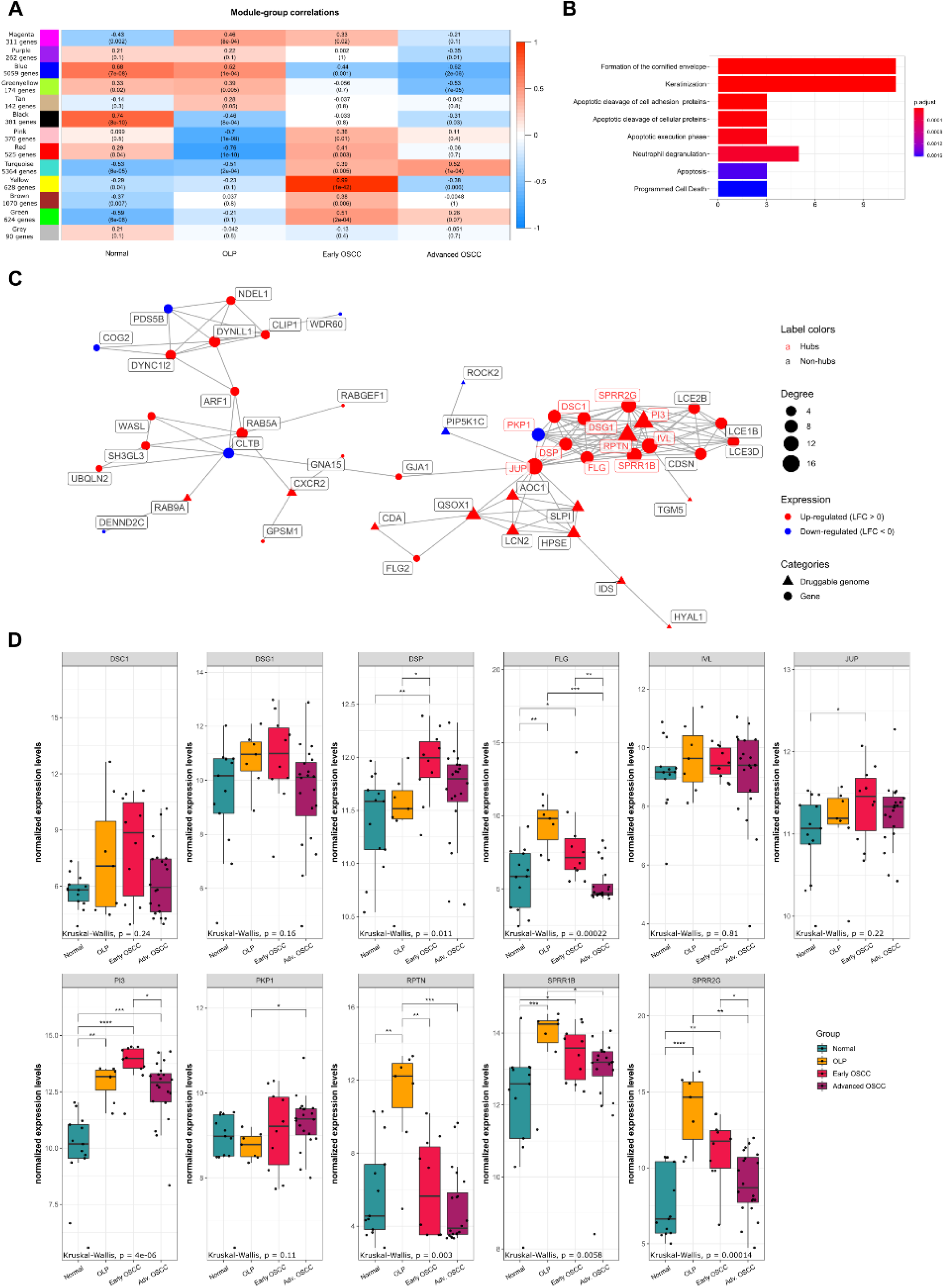
Analysis of co-expressed genes and their regulation. (A) Correlations between the co-expression modules and each of the sample groups (normal tissue, OLP, early OSCC, and advanced OSCC). Cell colors correspond to Pearson correlation values between each co-expression module (rows) with groups (columns), from blue (100% inverse correlation) to red (100% direct correlation). Numbers in each cell refer to Pearson’s correlation r values, with p-values represented below in parentheses. Correlations with p < 0.05 were considered statistically significant. (B) Pathway enrichment of genes in the magenta module using ORA. All represented pathways are significantly enriched (FDR > 0.05). Bar length and colors indicate gene counts and BH-adjusted p-values, respectively. (C) Representations of the magenta module as a PPI network (largest connected component). Node size, shape, and color represent degree, category, and LFC in OLP, respectively. Red node labels indicate hubs. (D) Hub genes from the magenta module compared between OLP, both OSCC groups, and normal tissue. Box plots represent the normalized expression distribution in each group. Comparisons among groups were made using the Kruskal-Wallis test followed by Dunn’s post-hoc test, with p-values lower than 0.05 considered significant for both tests. *, p < 0.05; **, p < 0.01; ***, p < 0.001; ****, p < 0.0001.

To better understand the relationship between OLP and OSCC, we have investigated which co-expression modules had a similar correlation to both conditions simultaneously. For each module, the Pearson correlation coefficient of the module eigengene to the sample groups was calculated. None of the identified modules had a simultaneous significant correlation with OLP and early and late stage OSCC. Interestingly, only the magenta module showed a significant and positive correlation with OLP (Pearson’s correlation r = 0.46; p < 0.0001) and early OSCC (r = 0.33; p = 0.02). Co-expressed genes belonging to this module were mainly associated with keratinization and formation of the cornified envelope (Fig. 3B).

### Network analysis

Co-expression modules, while grouping correlated genes, offer only a glimpse of their dynamic in the cells. To understand how genes in the magenta module interacted, we added another layer of information, searching for protein-protein interaction (PPI) data in the STRING database and building a network (Fig. 3C).

By characterizing genes from the network according to their characteristics and connectivity, we identified 11 hubs, 15 transcription factors, 6 clinically actionable genes and 112 members of gene families in the druggable genome, with some of those genes classified in more than one category (Table S8). Additionally, drug-gene interactions were identified using all genes in the module. A total of 89 drugs were identified, which interacted with 3 of the 11 hubs (Table S9).

The hubs’ expression levels were compared among conditions, with *PI3* being the only gene significantly up-regulated in OLP and all OSCC stages compared to the normal mucosa. Additionally, *FLG, SPRR1B*, and *SPRR2G* were significantly up-regulated in OLP and early OSCC, while *DSP* and *JUP* showed higher expression levels only in early OSCC and *RPTN* was up-regulated only in OLP. The four remaining hubs (*DSC1, DSG1, IVL*, and *PKP1*) didn’t exhibit significant differences in expression compared to the normal mucosa (Fig. 3D).

The hubs with known drug-gene interactions are *PI3* (up-regulated in OLP and OSCC), *IVL* (up-regulated in OLP and early OSCC), and *DSP* (up-regulated in OLP and OSCC) (Fig. 3 C-D, Table S9). This result may direct future selections of drug targets that could lead to efficient treatment for both diseases simultaneously.

### Expression drug-response analysis

Considering the possibility that the 35 DEGs in common between OLP, early and advanced OSCC are involved in the potential for malignant transformation, we investigated pharmacological agents that would best reverse the signature of these genes. A clustergram of the main drugs was obtained according to L1000 CDS2 output, which demonstrates the expected positive or negative regulation of each drug in relation to each DEG by comparing them to the LINCS L1000 small molecule-related expression profiles (Fig. 4). The top fifty matched signatures, corresponding to 42 drugs, were identified based on perturbation data for nineteen of the overlapping DEGs. Six (14.3%) of these drugs are already in use in the clinic or tested in clinical trials, as indicated in Table S10. Among the predicted drugs the most represented class was that of the PI3K/mTOR pathway inhibitors, which includes INK-128, GSK 1059615, GDC-0980, Torin-2, KU 0060648 Trihydrochloride, AZD-8055, and PI103 Hydrochloride. The signature of some genes has been reversed by a large number of drugs in the LINCS L1000 cells’ signatures such as *PI3* (32/42 drugs), *KRT17* (30/42), *KRT10* (29/42), *S100A7* (29/42), and *S100P* (23/42).

**Figure 4.**
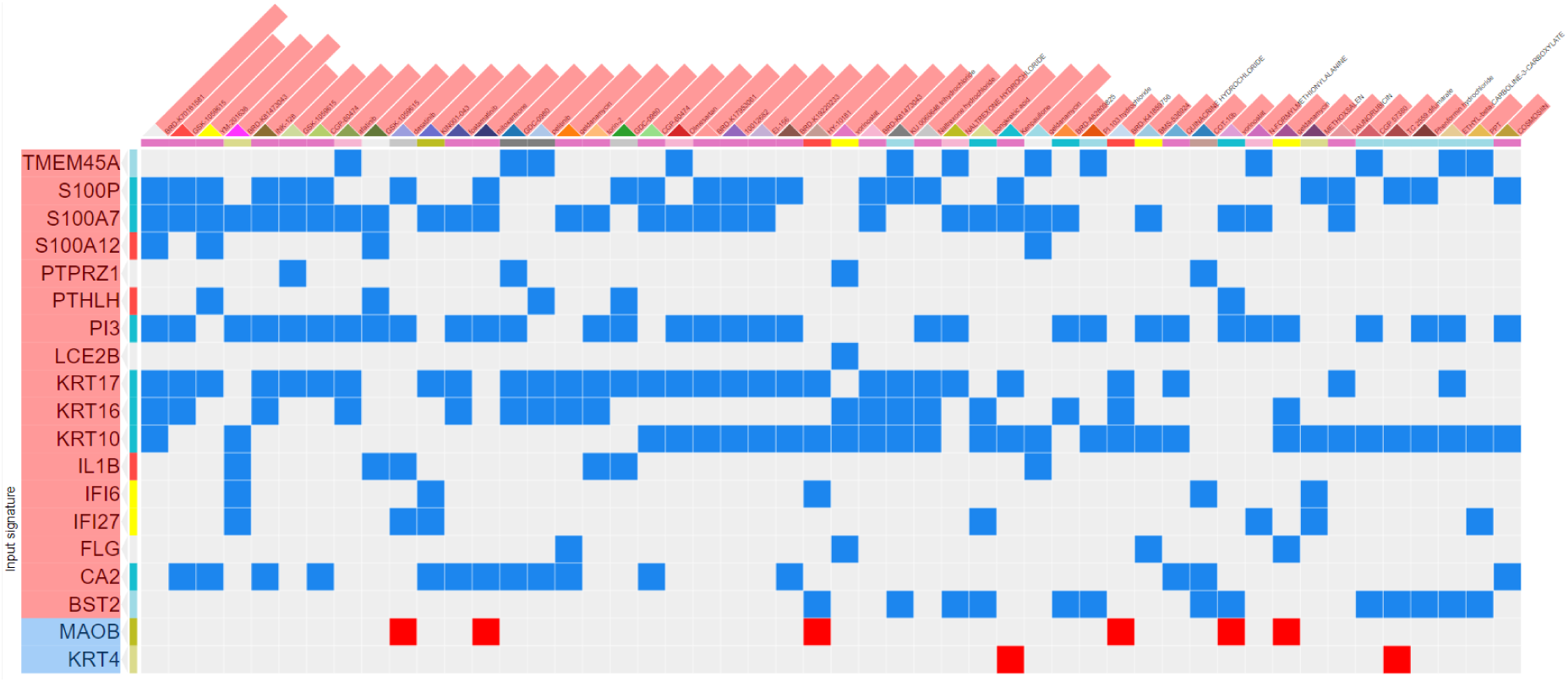
Potential drug target search from shared DEGs using the L1000 CDS^2^ tool on the LINCS Program platform. Genes are represented in rows and drugs in columns, with overexpressed genes colored red and underexpressed genes in blue. Colored squares represent drug-gene interactions, with blue cells representing inhibition and red cells representing gene activation.

### Drug repositioning evaluation

All identified drugs in this work were evaluated for their repositioning potential for use in OLP and OSCC. Of 89 drugs identified based on the network’s hub genes, 70 were tested in cancer and 67 were approved in their respective clinical trials. Of these, only 2 were already tested in OSCC and none was approved for use in this type of cancer, Cisplatin and Sunitinib (Table S9). Considering the drugs identified based on the DEGs using the L1000 CDS2 tool, 6 of the 42 drugs were tested and approved for use in cancer. None of them were tested in OSCC. Additionally, none of the drugs identified in both analyses have been tested in OLP (Table S10).

## Discussion

According to the World Health Organization, OLP is categorized as a potentially malignant disorder, with the possibility of progression to OSCC. Despite its well-known status, the relationship between OLP and OSCC remains controversial (Gonzalez-Moles et al., 2008; Peng et al., 2017; Shen et al., 2011). Also, few studies have compared OLP with OSCC using high-throughput data (Giacomelli et al., 2009; J. Liu et al., 2020) and, to the best of our knowledge, no studies have simultaneously analyzed OLP and OSCC progression using high-throughput molecular data analysis. Based on this lack of information, we have sought to uncover which mechanisms are involved in this transformation. OLP, early OSCC, and advanced OSCC mRNA microarray data were analyzed to investigate similarities between gene expression profiles from the three conditions and correlating them to OLP malignant transformation potential.

Differential expression analysis revealed a subset of genes that are consistently modulated in OLP and OSCC, suggesting that there are similar pathways activated in the two conditions. Most of these genes modulated in both OLP and OSCC are related to keratins and keratinocyte differentiation. Shimada et al. (2018) demonstrated that cornified envelope formation proteins would be up-regulated in OLP, contributing to a hyperkeratosis state commonly observed in this pathology. In accordance with this, OLP presented the enriched and up-regulated cornified envelope formation pathway that was also found in early OSCC, suggesting that changes in proteins belonging to this pathway may be involved in the malignization processes leading to progression from OLP to OSCC. Our co-expression and network analyses reinforce this hypothesis by revealing that genes belonging to the cornified envelope formation and keratinization pathways were consistently co-regulated in OLP and early OSCC. Among the genes belonging to these pathways, *PI3* (peptidase inhibitor 3) stands out, which was also significantly upregulated in the three conditions. Kengkarn et al. (2020), through microarray analysis, identified that *PI3*, along with *KRT17*, were upregulated in 100% of OSCC samples (39 cases), suggesting that these genes can be used as molecular biomarkers for patients with OSCC. This gene was also identified as belonging to the druggable genome, a set of genes that code for proteins belonging to families suitable for drug development, which opens possibilities for developing compounds that interact with its protein (Hopkins and Groom, 2002; Russ and Lampel, 2005). In addition, it is interesting to note that the *SPRR1B, SPRR2G* and *FLG* genes were significantly upregulated in OLP and early OSCC, but not in advanced OSCC, when compared to healthy oral mucosa samples. The *SPRR* (Small PRoline Rich) gene family encodes cornified envelope precursor proteins and are closely related to keratinocyte differentiation (Patel et al., 2003). The role of *SPRR2G* in OLP and OSCC has not yet been investigated, however, it has been seen that squamous cell carcinoma of the vulva overexpresses this gene (Micci et al., 2013). As for *SPRR1B*, it was shown that its expression is up-regulated in OSCC-derived stem cells, in addition to having a role in the growth and proliferation of these cells by regulating *RASSF4*, a tumor suppressor gene related with the MAPK pathway, suggesting that the overexpression of *SPRR1B* may be related to the carcinogenesis of OSCC, as well as the maintenance of stem cells of this carcinoma (Michifuri et al., 2013). Filaggrin, coded by the *FLG* gene, is a protein located in the stratum corneum of the skin, contributing to its integrity and strength (McGrath and Uitto, 2008), and patients with OLP have an altered distribution and overexpression of this protein in the oral mucosa (Larsen et al., 2017). Though filaggrin may be involved in lesions in the oral mucosa (Itoiz et al., 1985), further investigations are needed to show the role of this protein and the malignancy potential of OLP.

*KRT17* is overexpressed in OSCC and it may be associated with tumor progression by stimulating multiple signaling pathways (Kitamura et al., 2012; Ohkura et al., 2005). Shen et al. (2006) suggested that Keratin 17 may serve as immunodominant T cell epitopes by stimulating peripheral blood lymphocytes in psoriasis. Liu et al. (2020) demonstrated that high *KRT17* expression levels and tumor differentiation stage were significantly associated with overall survival in 64 patients with esophageal squamous cell carcinoma (ESCC), suggesting that *KRT17* may be a tumor-promoting factor and that an increase in the expression of this keratin may contribute to the malignant progression of the carcinoma. Furthermore, the authors demonstrated that *KRT17* plays a role in proliferation, migration, growth, and metastasis of ESCC cells *in vitro* and *in vivo*. Similar results were observed by Wang et al. (2019) in non-small cell lung cancer (NSCLC). The authors noted that high *KRT17* expression levels correlated with poor prognosis in NSCLC, especially in lung adenocarcinoma. Although not significantly differentially expressed in OLP samples from our validating dataset, which may be related to the low number of samples available, the upregulation of *KRT17* in OLP may also play an important role associated with the malignization process. Interestingly, *KRT17* is one of the main therapeutic targets found in our analyses. Based on these results, it is suggested that *KRT17*, as well as the related pathways, are important for studying the pathology and malignancy potential of OLP to OSCC, since this gene seems to be involved in the transition between both diseases. Besides, our data reveal the elevated and significant expression of *KRT10* in contrast with low levels of *KRT4* in both conditions when compared to the normal oral mucosa. Sakamoto et al. (2011) demonstrated that *KRT4* down-regulation in oral squamous cell carcinoma is associated with changes in the morphology of the epithelium and could be associated with the overexpression of other keratins such as *KRT17*. Therefore, the authors suggest that *KRT4* may serve as a diagnostic biomarker for OSCC. The same was suggested by Schaaij-Visser et al. (2009) when revealing that the low expression of *KRT4* in samples from patients with head and neck squamous cell carcinomas (HNSCC), including OSCC samples, may serve as a screening biomarker for local recurrence risk and allow selection for adjuvant treatment or tertiary prevention studies. Similar results were observed by Liao et al. (2012) in OLP lesions, which generally affect the non-masticatory mucosa (such as the bilateral buccal mucosa), with a shift in keratin expression observed by an increased expression of *KRT10* and reduced expression of *KRT4*. These data were in agreement with our analyses on the reduction of *KRT4* in OLP. However, further investigation is needed regarding *KRT10*, because although the validation analyses suggest an increase in its expression, the difference in expression is not significant, in disagreement with what was observed in our analysis.

Our analysis showed that antigen presentation is the only pathway enriched and upregulated in the three evaluated conditions. Antigen cross-presentation seems to be involved in the early processes of OLP pathogenesis.

However, the nature of the antigen responsible for triggering the immune response in OLP has not yet been unveiled. When analyzing the genes present in the antigens cross-presentation pathway in OLP and early and advanced OSCC, we observed genes associated with different tumor responses such as genes belonging to HLA (*HLA-E*; *HLA-F* and *HLA-G)*, among which *HLA-G* which has been identified as an immune evasion-related gene in different tumors through the inhibition of effector cells such as NK, T cells, monocytes, and dendritic cells (Krijgsman et al., 2020). This gene was associated with OSCC prognosis and indicated as a new therapeutic target (Shen et al., 2018). In addition, most of the genes associated with antigen cross-presentation belong to the proteasome subunits, which have been associated with worse prognosis, tumor progression, and metastasis in different tumors (Ding et al., 2020; Kakumu et al., 2017; Munkácsy et al., 2010; Tan et al., 2018). Interestingly, among these *PSMB10*, one of the subunits of the immunoproteasome, was found differentially expressed in the three conditions. Immunoproteasome has been the subject of several studies due to its role in the differentiation of T cells, cytokine regulation, and tumor progression (Chen et al., 2020; Kiuchi et al., 2021; Zerfas et al., 2020). In addition, it is known that the peptides generated by the immunoproteasome for MHC class I are capable of generating a more efficient and accentuated cytotoxic lymphocyte response than those generated by the constitutive proteasome, contributing to CD8+ T cell infiltration (Groettrup et al., 2001; Kloetzel, 2001). We hypothesise that antigen presentation mediated by the immunoproteasome may contribute to the immune infiltrate profile present in OLP and OSCC. Further investigations, however, are needed on the interaction between antigen presentation and cytokines participating in the pathogenesis of OLP and OSCC, mainly on their role in the potential for malignant OLP transformation. Although previous studies have shown that the immune infiltrate profile in OLP is predominantly composed of CD8+ and CD4+ T cells (Iijima et al., 2003; Wang et al., 2016), our analyses through gene signature deconvolution have demonstrated a significantly reduced proportion of CD8+ T lymphocytes and NK cells in those samples. Still on the predominant microenvironment components in OLP, we demonstrate an up-regulated signature of the chemokine CXCL-13, which has a dual role in tumorigenesis (Kazanietz et al., 2019).

The participation of Th17 cells in OLP and OSCC pathogenesis has also been recently investigated. Th17 is related to the maintenance of chronic inflammation in many conditions, notably autoimmune diseases (Awasthi and Kuchroo, 2009). Pathway enrichment and gene expression analyses performed in this study demonstrated that pathogenic Th17-related pathways were positively correlated to OSCC, which is corroborated by findings by Gaur et al. (2012), that also demonstrated an increase in Th17 cell prevalence in OSCC patients’ peripheral blood when compared to a healthy control group. In contrast, it is observed that in the heatmap of Th17 phenotype-related genes, the general expression profile of most OLP samples is closely related to the sample profile of the normal mucosa. However, two samples clustered with early and advanced OSCC samples. This result suggests that different samples of OLP may have different expression profiles, which may be related to lower or higher risk of malignant transformation. Also, individual investigations of gene expressions related to the pathogenic Th17 molecular signature in OLP showed significantly high levels when compared to samples of normal oral mucosa for several genes such as *TGFB3, IL1B, HIF1A, LTA*, and *LTB*. The role of HIF-1α in the potential for malignant transformation of OLP has been the subject of investigation. Wang et al. (2017) showed that *HIF1A* was upregulated in OLP and OSCC samples, contributing to changes in the expression of genes involved in adaptation to hypoxia and tumor progression. Additionally, Yang et al. (2020) demonstrated that the activation of *HIF1A*, by the accumulation of succinate, plays a fundamental role during the malignant transformation of OLP by stimulating the apoptosis of keratinocytes. Besides, as *HIF1A* is also related to increased transcription of *IL1B* by cells of the immune system (Corcoran and O’Neill, 2016; Ge et al., 2019), it is suggested that the increased expression of both may be correlated. Together, these results suggest that OLP may have elements of a tumor-like microenvironment as proposed by Peng et al. (2017).

Interestingly, in our drug signature analysis, IL-1β was suggested as a target for treatment with the drugs such as GSK-1059615, Torin-2, and GDC-0980, which are PI3K/mTOR inhibitors (Leontieva and Blagosklonny, 2016). Only GSK-1059615 has already been investigated in OSCC, being able to reduce the proliferation of OSCC cell lines (Yang et al., 2020). In OLP lesions, Ma et al. (2019) showed the overexpression of phosphorylated IGF1R and TRB3, which are related to the PI3k/AKT/mTOR signaling pathway, suggesting that this pathway mediates the relationship between T cells and keratinocytes, and influences the imbalanced cytokine networks on the immune microenvironment. According to the data presented in this study, four other mTOR pathway inhibitors were identified as candidate drugs (PI-103; INK-128; KU-0060648 and AZD-8055). Among them, PI-103 and INK-128 treatment demonstrated inhibition of cell growth and proliferation of OSCC (Aggarwal et al., 2019; Liang et al., 2019). Also, AZD-8055, an inhibitor of both mTORC1 and mTORC2, was able to induce autophagy in HNSCC cells (Li et al., 2013). Although corticosteroids are recommended for the treatment of OLP, in this work we demonstrated different pharmacological agents that could assist in the treatment and possibly interfere with the malignancy potential of such lesions.

Although our analyses have uncovered some of the genes and pathways potentially involved in the transformation from OLP to OSCC, the limited availability of public OLP data made it difficult to acquire gene expression data. However, even though the low number of available OLP data led us to perform the analysis using batch-corrected datasets from different platforms, we were able to discover important biological similarities between the conditions which may point us to better understand the malignization process.

In conclusion, *in silico* analysis revealed that OLP is a pathology that has a proximity to the gene expression profile of OSCC, mainly with early OSCC. This result is compatible with the fact that OLP is a differential diagnosis of epithelial precursor lesions, namely leukoplakia and erythroleukoplakia, which can give rise to initial OSCC if not removed surgically. We clearly reveal signatures in common with the two conditions that can be important targets for drug treatment, as well as in the development of diagnostic and prognostic strategies for the disease. It is considered that OLP and OSCC have multifactorial etiology, and the intersections between the keratinization and differentiation of lymphocytes are interesting potential targets for further investigation.

## Materials and methods

### Datasets

Gene expression data from mRNA microarray experiments were obtained from NCBI’s Gene Expression Omnibus (GEO) (Barrett et al., 2013) using the GEOquery R package (Davis and Meltzer, 2007). The datasets used corresponded to accession numbers GSE52130, GSE56532, and GSE41613. From the dataset GSE52130 only oral samples were used, consisting of 7 OLP samples and 7 normal oral tissue samples, with expression values measured using the Illumina HumanHT-12 V4.0 expression BeadChip array platform. GSE56532 consisted of gene expression from 10 advanced OSCC samples and 6 normal oral mucosa samples, measured using the Affymetrix Human Gene 1.0 ST Array platform. GSE41613 is a dataset containing gene expression data of 97 HPV-negative samples from OSCC at varied stages, measured using the Affymetrix Human Genome U133 Plus 2.0 Array platform. Since this dataset contained a much larger sample size compared to the others, we have randomly selected a subset of 20 samples, ten of them at stages I and II (samples GSM1020161, GSM1020136, GSM1020149, GSM1020147, GSM1020122, GSM1020134, GSM1020188, GSM1020189, GSM1020138, and GSM1020179), and ten of them at stages III and IV (samples GSM1020128, GSM1020141, GSM1020185, GSM1020101, GSM1020121, GSM1020135, GSM1020123, GSM1020111, GSM1020102, and GSM1020187), to keep group sizes similar and avoid disproportionately large groups to skew subsequent analyses.

Results were validated using independent microarray expression datasets from GEO as well as RNA-seq expression data from The Cancer Genome Atlas (TCGA). Validation microarray expression datasets from GEO correspond to accession numbers GSE38616 and GSE3524. GSE38616 consisted of 7 OLP and 7 normal oral mucosa, with expression measured on the Affymetrix Human Gene 1.0 ST Array platform. GSE3524 was composed of 16 OSCC samples in stages II and IV and 4 normal tissue samples, with staging information missing for 2 of the tumor samples, which were removed from the analysis. mRNA expression for this dataset was measured on the Affymetrix Human Genome U133A Array platform. Outliers were identified in each dataset using PCA and removed. GSE38616 had three outliers, GSM946266 (OLP), GSM946263 (OLP), and GSM946254 (Normal); while GSE3524 had one outlier, GSM80467 (advanced OSCC). For TCGA samples, expression data were downloaded in the form of raw counts from the Head and Neck Squamous cell Cancer (TCGA-HNSC) project using the TCGAbiolinks R package (Colaprico et al., 2016). Primary tumor and normal samples belonging to the “Other and unspecified parts of tongue”, “Base of tongue”, “Lip”, “Palate”, “Gum”, “Floor of mouth”, “Other and unspecified parts of mouth”, and “Oropharynx” sites were considered OSCC samples and used in this step.

### Data integration and cross-platform normalization

Data integration and cross-platform normalization were performed in the discovery datasets according to the methods described in (Binato et al., 2018) and (Walsh et al., 2015) Files containing probe-level intensity data (CEL files for Affymetrix arrays and txt files for Illumina BeadChip arrays) were downloaded using GEOquery. Raw data files were preprocessed using the appropriate package for each platform (*oligo* for Affymetrix and *beadarray* for Illumina). Probe level data was extracted, background corrected, and normalized, with the robust multi-array average (RMA) method used for Affymetrix datasets and neqc for the Illumina BeadChip dataset. Probes were mapped to genes using each platform’s annotation, with the resulting matrix containing 16,656 features. When multiple probes corresponded to the same gene, normalized expression was aggregated to gene mean values. Batch effects were corrected using the ComBat method (Johnson et al., 2007) implemented in the sva R package (Leek et al., 2012), considering dataset of origin and sample type as variables. Samples were grouped using principal component analysis (PCA) to validate that the batch effect correction was successful.

### Differential expression analysis

Differential expression analysis was conducted using the limma R package (Ritchie et al., 2015). Gene expression from OLP, early OSCC (stages I and II), and advanced OSCC (stages III and IV) were compared to normal samples. Differentially expressed genes (DEGs) were identified based on the following cutoffs: Benjamini-Hochberg (BH)-adjusted p < 0.05 and log_2_ fold change (LFC) > 2. DEGs from each comparison were overlapped using the InteractiVenn online tool (Heberle et al., 2015). Heat maps were constructed using the pheatmap R package (Kolde, 2019), using normalized expression values z-scored across samples. Rows representing genes were clustered using Pearson correlation. Columns representing samples were clustered using the *hclust* function, with non-supervised hierarchical clustering performed based on sample distance, measured as 1 - Pearson correlation coefficient. For individual genes, boxplots were plotted using the ggplot2 package (Wickham, 2016). Comparisons among groups were made using the Kruskal-Wallis test followed by Dunn’s post-hoc test, with p-values lower than 0.05 considered significant for both tests. For RNA-Seq data, expression counts were normalized using variance stabilizing transformation through the DESeq2 R package (Love et al., 2014).

Differences in some of the genes’ expressions among analyzed groups were also evaluated using the Kruskal-Wallis test followed by pairwise comparisons using Dunn’s test. In both cases, p < 0.05 indicated a significant difference.

### Pathway enrichment analysis

Gene Set Enrichment Analysis (GSEA) was performed on LFC-ranked genes in each condition using the WebgestaltR R package (Liao et al., 2019; Subramanian et al., 2005). The analysis was performed with 1,000 permutations. The false discovery rate (FDR) cutoff was 0.05. Pathways with a minimum of 5 genes were selected. Result lists for each group had redundant pathways reduced using Webgestalt’s implementation of the Affinity Propagation algorithm. A subsequent filter kept all gene sets that were enriched in OLP and at least one of the OSCC stage groups. These results were presented in dotplots using the ggplot2 package.

Overrepresentation analysis was performed for overlapping and OLP-exclusive DEGs, as well as co-expression modules’ genes using the ReactomePA R package (Yu and He, 2016). Pathways with a minimum of 5 genes and FDR < 0.05 were selected, and the Benjamini-Hochberg (BH) method for multiple testing p-value correction was used (Benjamini and Hochberg, 1995). Pathways from the Reactome database were used for both GSEA and ORA (Jassal et al., 2020).

### Immune Infiltration Cells Analysis

Tumor immune infiltration cells composition was estimated using CIBERSORTx (Newman et al., 2019). This tool uses a deconvolution algorithm to estimate immune cell types using gene expression data from samples composed of multiple cells (bulk). Batch-corrected, normalized expression data was used to estimate TIICs using CIBERSORTx’s gene signatures for 22 cell types. These cell populations include naïve B cells, memory B cells, plasma cells, 7 T cell types (CD8+ T cells, naïve CD4+ T cells, resting CD4+ memory T cells, activated CD4+ memory T cells, follicular helper T cells, Tregs, γδ T cells), macrophages (M0 macrophages, M1 macrophages, M2 macrophages), resting mast cells, activated mast cells, resting NK cells, activated NK cells, resting dendritic cells (resting DC), activated dendritic cells (activated DC), monocytes, eosinophils, and neutrophils.

Stacked bar plots were generated from relative cell type populations using ggplot2. Samples were clustered using the hclust function. Distances based on 1 - Pearson correlation coefficient were used for Ward clustering of samples. Population scores were individually compared between groups using Kruskal-Wallis test, followed by pairwise comparisons using Dunn’s test. In both cases, significant differences were identified by p < 0.05.

Additionally, the signatures of genes related to pathogenic and non-pathogenic Th17 cells were investigated using a 33-gene signature panel based on a previous characterization of Th17 phenotypes in the literature (Lee et al., 2012). Gene expression and sample clustering visualizations for pathogenic and non-pathogenic Th17 cell signatures were made using the pheatmap R package (Kolde, 2019). Boxplots for the signature’s genes were plotted using the ggplot2 package (Wickham, 2016). Comparisons among groups were made using the Kruskal-Wallis test followed by Dunn’s post-hoc test, with p-values lower than 0.05 considered significant for both tests.

### Co-expression analysis

Gene co-expression modules were constructed using the Weighted Gene Co-expression Network Analysis (WGCNA) R package (Langfelder and Horvath, 2008; Zhang and Horvath, 2005). The 15,000 genes with the highest median absolute deviation (MAD) were selected from the integrated dataset and used as input. A gene pair similarity matrix was generated based on Pearson correlation and converted to a weighted adjacency matrix by elevating it to a β value of 6. This matrix was used to build a topological overlap (TOM) and a dissimilarity matrix (1 - TOM). The dissimilarity matrix was used to build unsigned co-expression modules with a minimum size of 100 genes. WGCNA’s module-trait relationship function was used to calculate correlations between module eigengenes and each of the groups (OLP, early OSCC, advanced OSCC, and normal samples). Correlations were considered significant when | r | ≥ 0.3, and p < 0.05.

### Interaction networks construction and drug-gene interactions identification

Genes in the magenta module were used to build a protein-protein interaction (PPI) network using interaction data from the STRING database, v. 11 (Szklarczyk et al., 2019). Interactions with a confidence score < 0.9 and disconnected vertices were discarded. Hub genes were determined by selecting vertices with a degree over the 9th decile of the network’s degree distribution and comparisons in the individual hubs’ expression levels were performed as described in the differential expression section.

Gene categories and FDA-approved, antineoplastic drug-gene interactions for hubs were identified using the DGIdb online tool (Freshour et al., 2021). Clinically actionable genes, transcription factors, and genes coding for protein families belonging to the druggable genome were identified (Hopkins and Groom, 2002; Russ and Lampel, 2005). Information such as log_2_ fold change, identified gene categories, degree, and whether a gene is a hub were also added to the network. The graph’s largest connected component was used for visualization. Network manipulation was made using the igraph and tidygraph R packages (Csardi and Nepusz, 2006; Pedersen, 2020). Network plots were constructed using the ggraph R package (Pedersen, 2021).

### Search for expression drug-response for differentially expressed genes

Overlapping DEGs were used to search for drugs able to revert their expression signatures using the L1000 Characteristic Direction Signature Search Engine (L1000CDS^2^) tool on the Library of Integrated Network-Based Cellular Signatures (LINCS) Program platform (Duan et al., 2016; Stathias et al., 2020). This tool compares input differentially expressed genes to LINCS-L1000 gene perturbation data. Drug-gene combinations were ranked by search score, calculated based on the overlap between input DEGs and signature DEGs, that is, gene sets that follow the same perturbation patterns (up-regulating underexpressed genes or down-regulating overexpressed genes) when interacting with a small molecule. The top 50 drug signatures are presented as output. Additionally, putative drug combinations among the small molecule signatures were estimated using this tool. Drug combinations were ranked based on their signature overlaps and the top 50 combinations were provided.

### Drug repositioning opportunities evaluation

Drugs identified both by interactions with hub genes and by investigating expression reversion signatures were evaluated for repositioning opportunities using the repoDB database, which compiles information from clinical trials (Brown and Patel, 2017). Drugs were searched for in the database and evaluated for whether they were already tested in OSCC, OLP, or other neoplasms. In the case when a drug was used in clinical trials, we have evaluated if it was approved or not for clinical use.

## Acknowledgements

The authors would like to thank the Plataforma Multiusuário de Bioinformática of Instituto Nacional de Câncer (INCA) for providing the infrastructure for performing the analyses.

## Supporting information

**Supplementary File 1** – Supplementary Figures S1-S6

**Supplementary File 2** – Supplementary Tables S1-S10

