## Supplemental file 1 for "Oral Lichen Planus and its relation with Oral Squamous Cell Carcinoma: new insights into the potential for malignant transformation"

### Supplementary figures

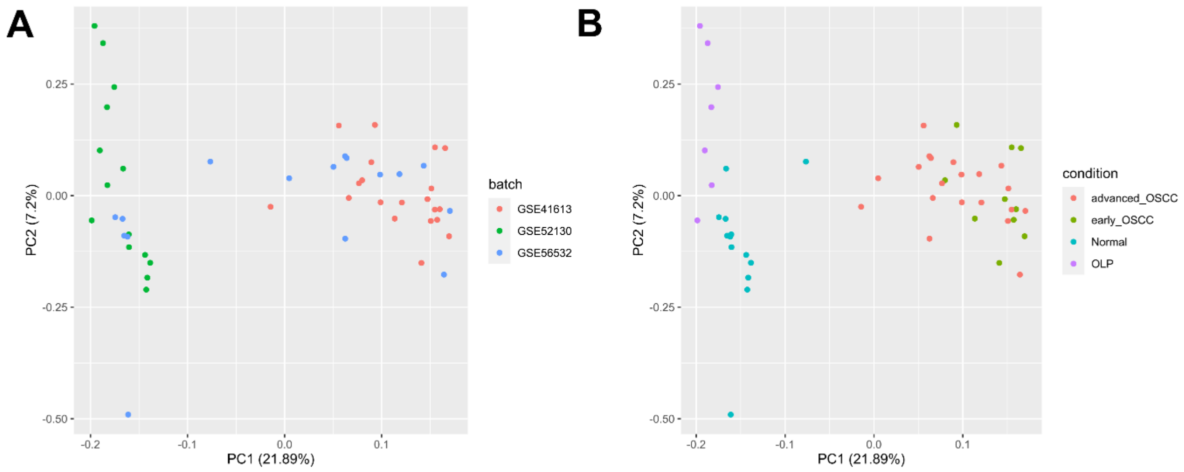


**Figure S1.** Principal component analysis (PCA) of unified dataset after batch effect correction using ComBat. (A) Samples identified by batch (dataset of origin). (B) Samples identified by group (Normal, OLP, early OSCC, advanced OSCC). The resulting graphs show that samples were no longer clustered by batch after correction, while still clustering by group, showing that the correction successfully removed batch effects while maintaining biological differences.

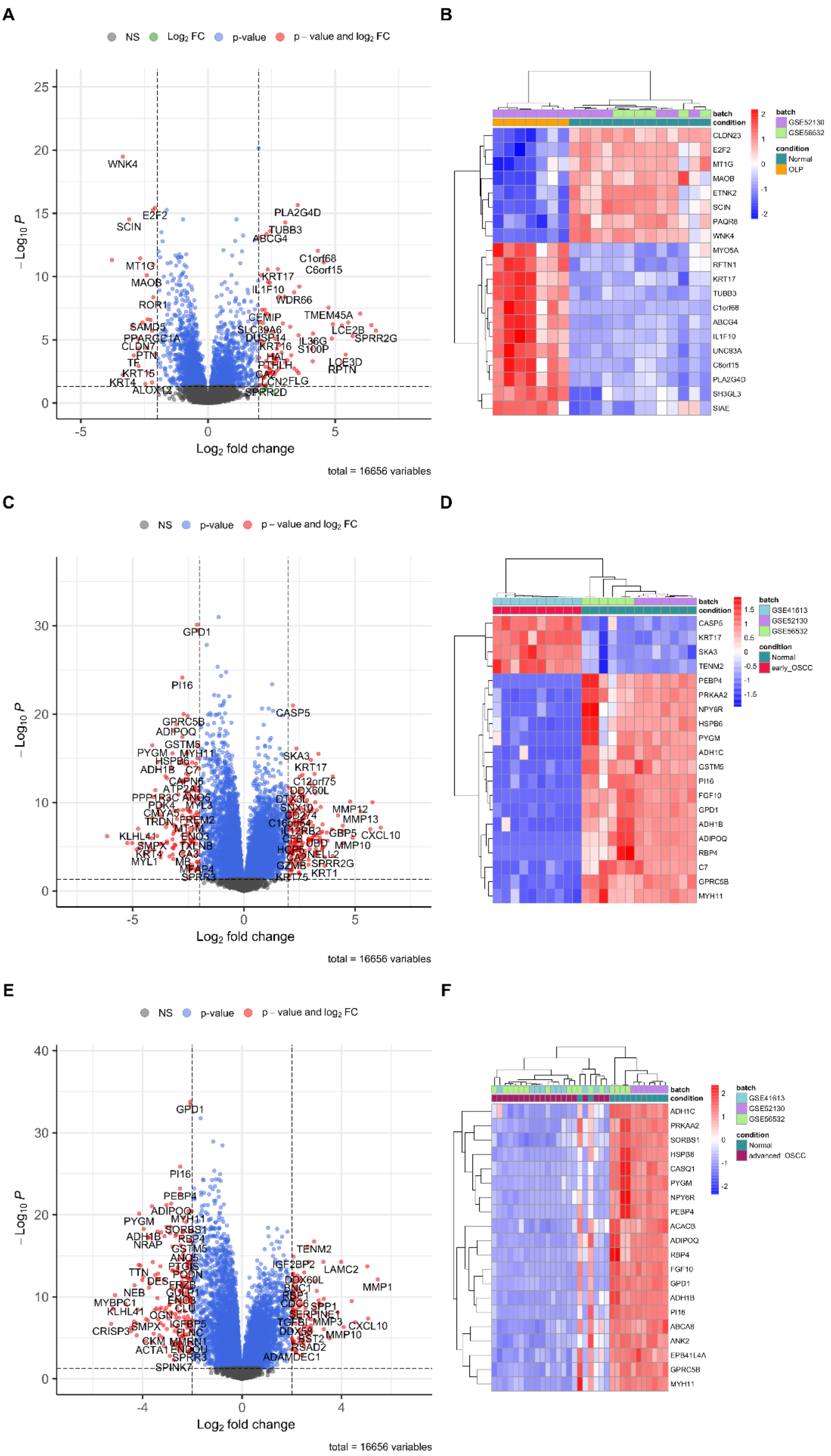


**Figure S2.** Differentially expressed genes in OLP (A, B), early OSCC (C, D), and advanced OSCC (E, F). Volcano plots (A, C, E) represent all DEGs in each group. Dots in red represent genes filtered by both adjusted $p < 0.05$, and $|\log_2 \text{fold change}| > 2$. Clustered heatmaps (B, D, F) represent the top 20 DEGs in each group.

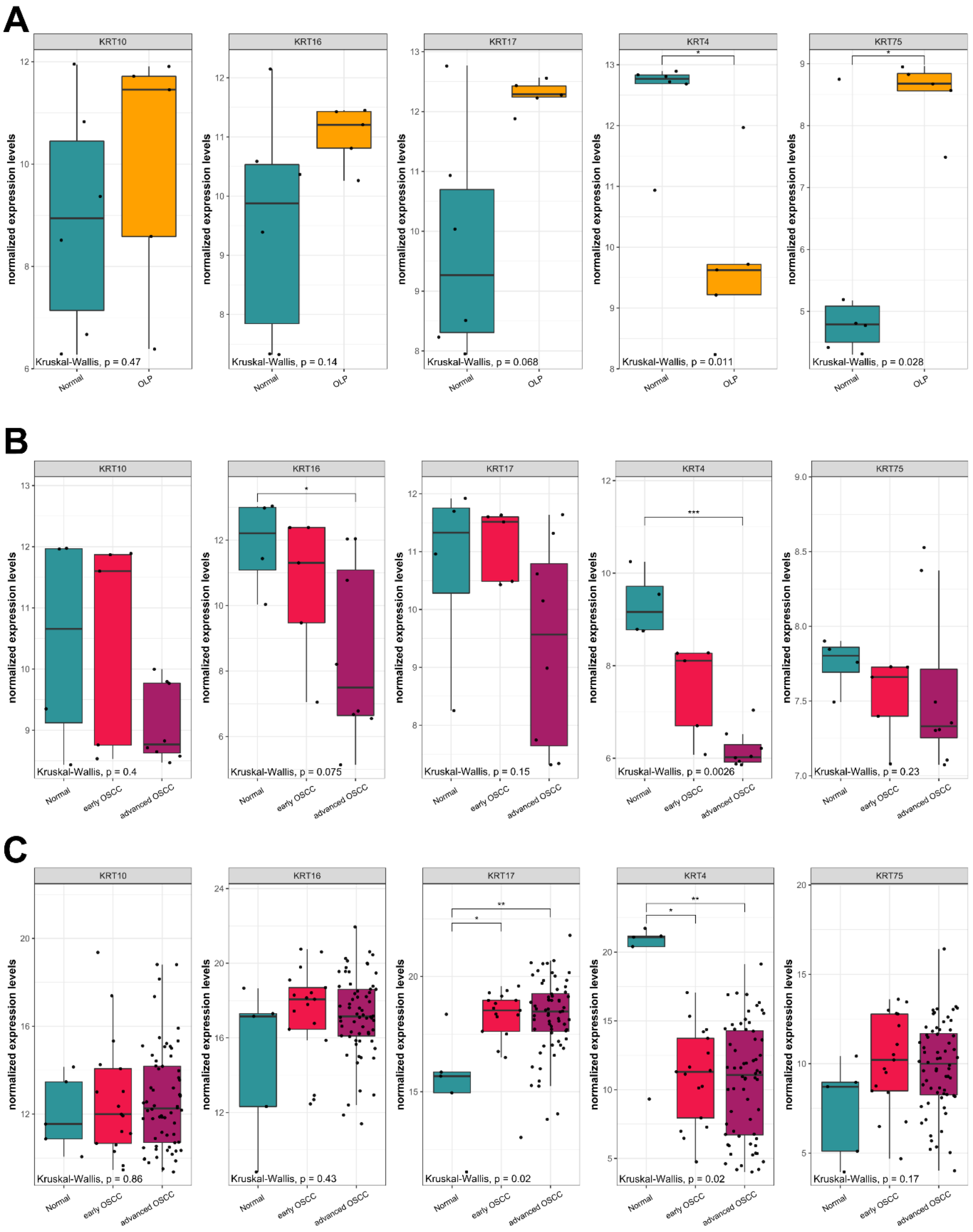


**Figure S3.** Differentially expressed keratin genes between OLP (microarray dataset GSE38616, A) and both OSCC groups (microarray dataset GSE3524, B; and TCGA, C) compared to normal tissue in validation datasets. Box plots represent each group's normalized expression value distributions, with dots representing each sample in the group. Comparisons among groups were made using the Kruskal-Wallis test followed by Dunn's post-hoc test, with p-values lower than 0.05 considered significant for both tests. *, $p < 0.05$; **, $p < 0.01$; ***, $p < 0.001$; ****, $p < 0.0001$.

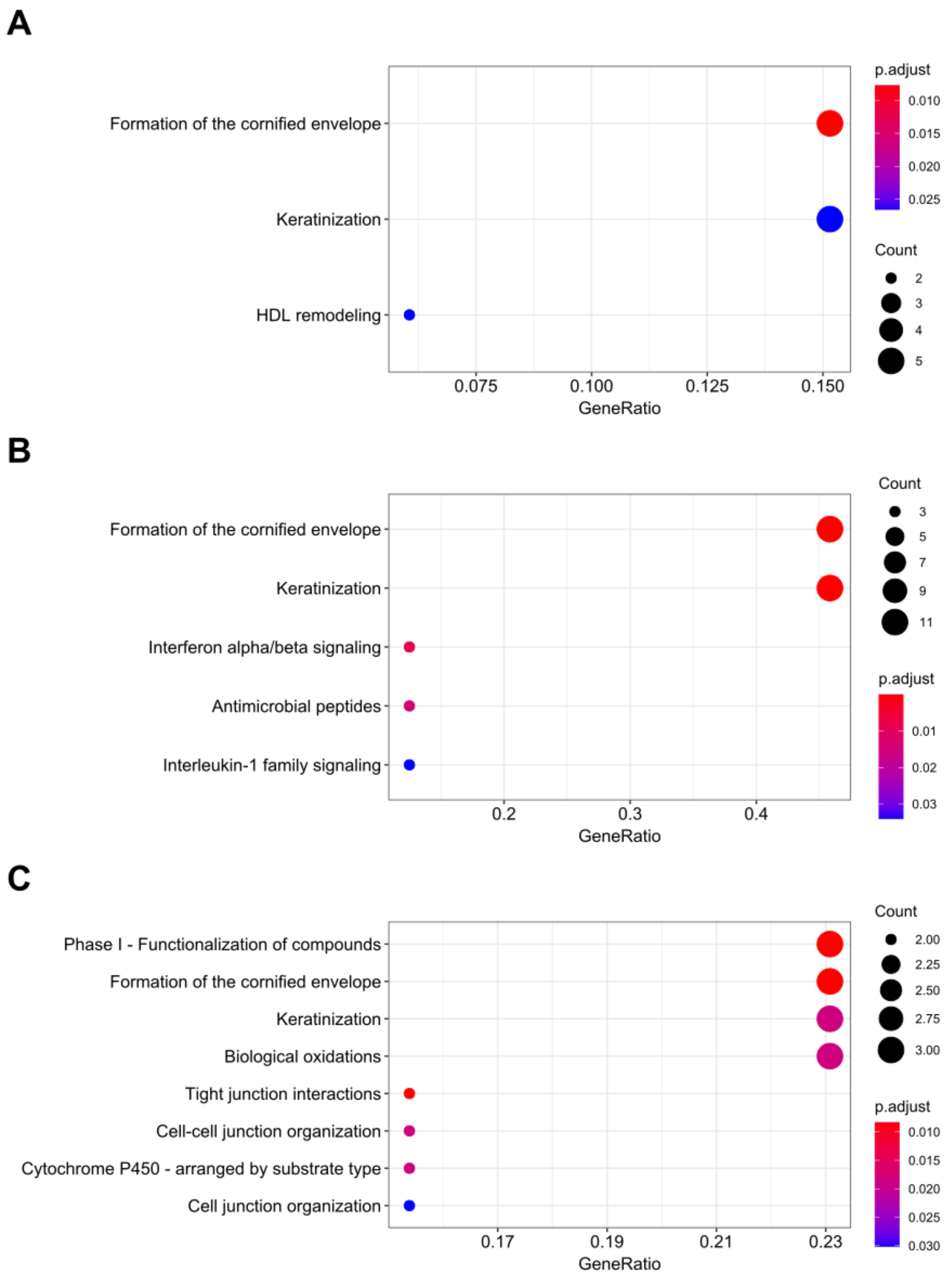


**Figure S4.** Pathway enrichment using overlapping DEGs. Pathways found in overrepresentation analysis (ORA) of (A) overlapping overexpressed between OLP and at least one OSCC group (both early and advanced); (B) OLP-exclusive overexpressed DEGs; (C) OLP-exclusive underexpressed DEGs. All pathways indicated are significantly enriched (FDR < 0.05). Color intensity indicates BH-adjusted p-value and dot sizes represent gene counts.

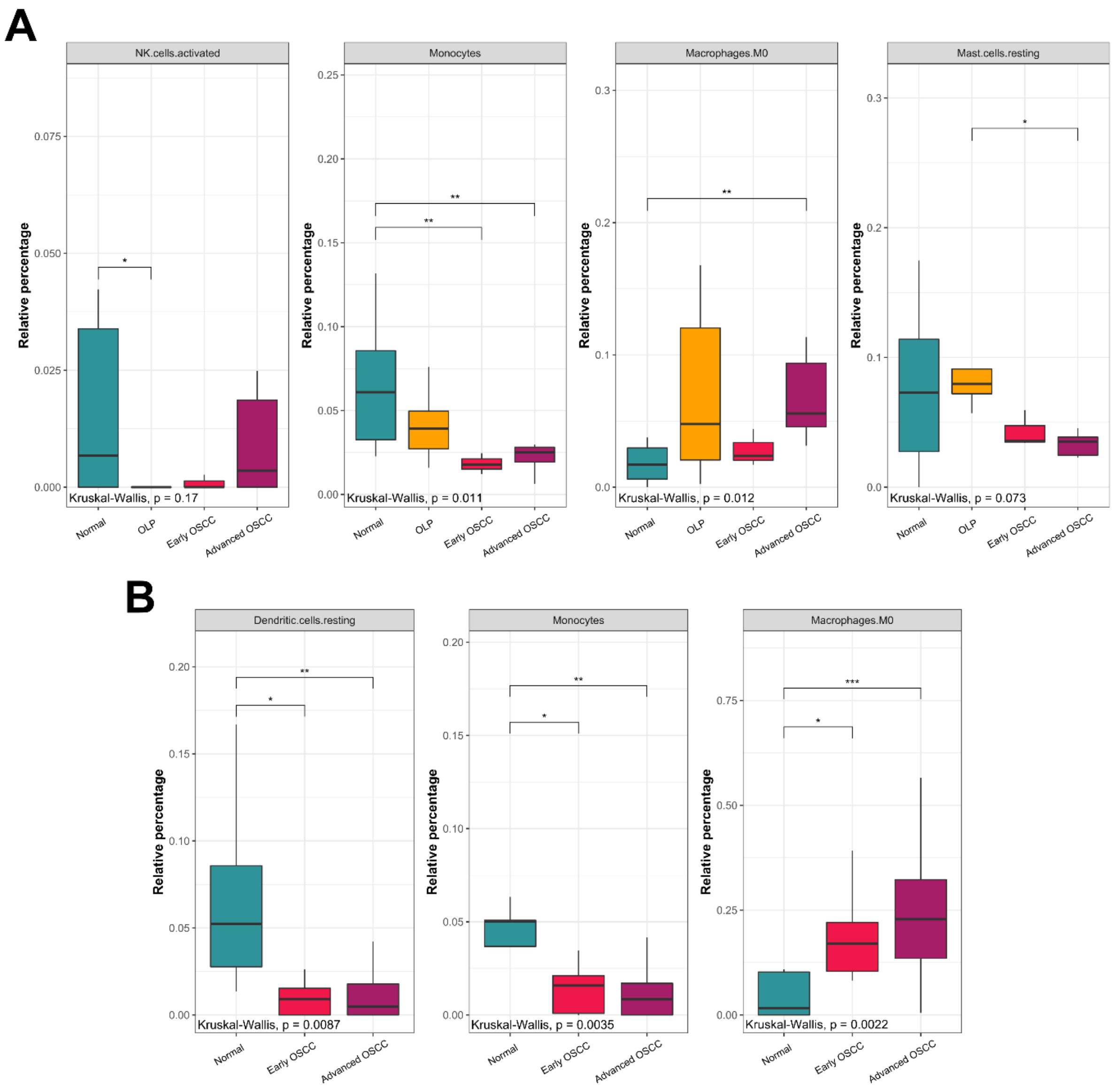


**Figure S5.** Proportions of cell populations with significant differences in validation datasets. (A) Proportions of activated NK cells, monocytes, macrophages M0, and resting mast cells, in OLP and both OSCC groups compared to normal tissue (microarray datasets GSE38616 and GSE3524). (B) Proportions of monocytes, macrophages M0, and resting dendritic cells in OSCC samples compared to normal tissue (TCGA dataset). Box plots represent each group's relative proportions of immune infiltrate cells as estimated by CIBERSORTx. Comparisons among groups were made using the Kruskal-Wallis test followed by Dunn's post-hoc test, with p-values lower than 0.05 considered significant for both tests. *, $p < 0.05$; **, $p < 0.01$; ***, $p < 0.001$; ****, $p < 0.0001$.

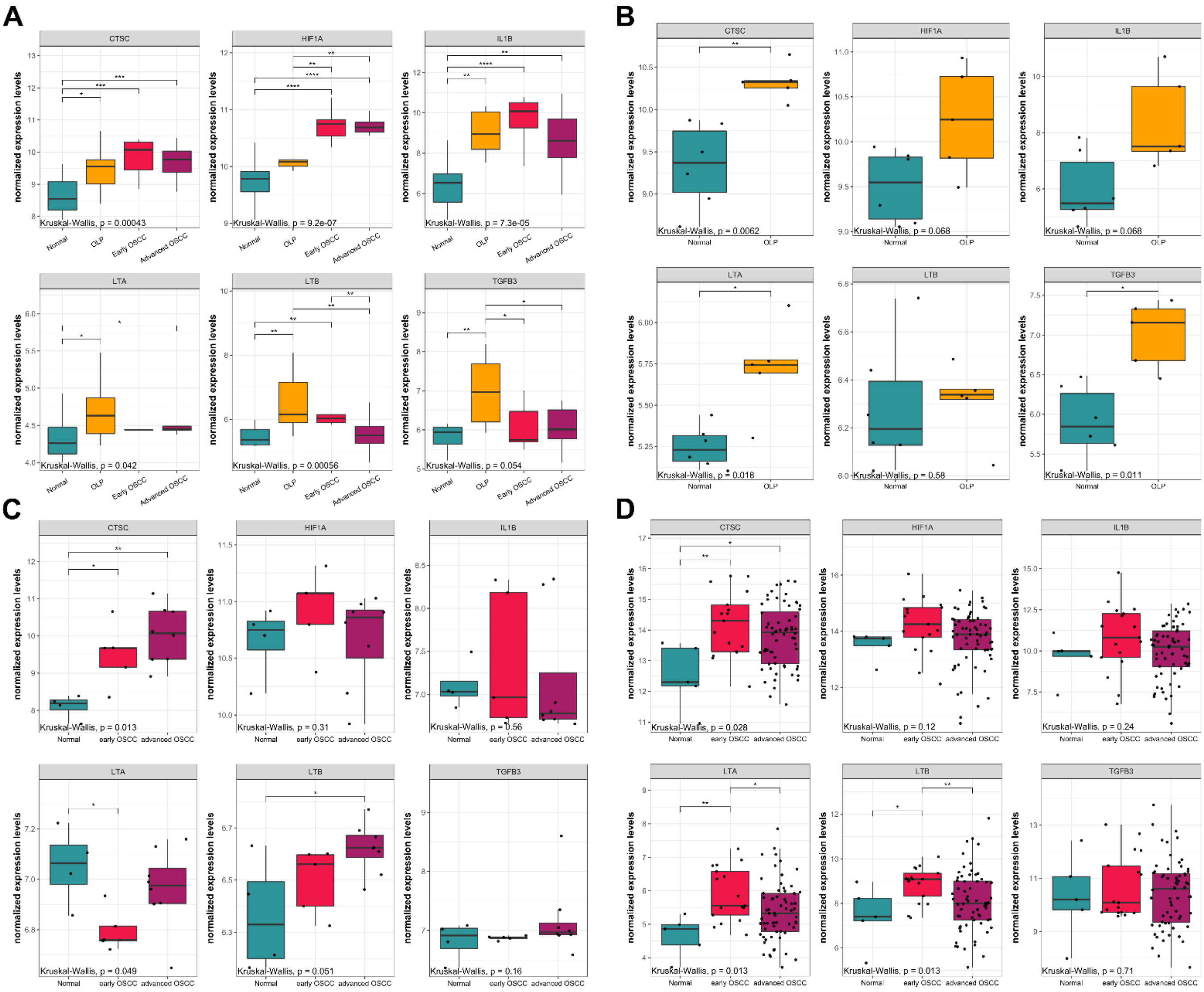


**Figure S6.** Th17-related genes compared between OLP, both OSCC groups, and normal tissue in discovery (A) and validation datasets (B-D). Box plots represent the normalized expression distribution in each group. Comparisons among groups were made using the Kruskal-Wallis test followed by Dunn's post-hoc test, with p-values lower than 0.05 considered significant for both tests. *, $p < 0.05$; **, $p < 0.01$; ***, $p < 0.001$; ****, $p < 0.0001$.
