## Supplemental file 2 for "Oral Lichen Planus and its relation with Oral Squamous Cell Carcinoma: new insights into the potential for malignant transformation"

**Table S1** - Differentially expressed genes in OLP in comparison to normal samples (healthy oral mucosa).

| Gene | log2 Fold Change | Average Expression | t | p-value | adjusted p-value | B |
|---|---|---|---|---|---|---|
| ***WNK4*** | -3.34766886090828 | 4.77342676305012 | -19.1597389109429 | 3.95689140173574e-24 | 3.29529915936552e-20 | 43.9844743166194 |
| ***PLA2G4D*** | 3.52865155968346 | 5.07398798629847 | 15.3323800441925 | 3.98506544559253e-20 | 2.21250833539298e-16 | 35.3192835368748 |
| ***E2F2*** | -2.0878790900879 | 5.74920407939165 | -15.0301526074396 | 8.81777590202027e-20 | 3.67172188560124e-16 | 34.5622460771881 |
| ***CLDN23*** | -2.16364790703234 | 5.65988497197448 | -14.7774222430864 | 1.72709348405692e-19 | 5.40577130973713e-16 | 33.9203577058731 |
| ***SCIN*** | -3.10523040887248 | 4.75489878711886 | -14.0421722636289 | 1.27450155401071e-18 | 2.94752466588493e-15 | 32.0062825342409 |
| ***TUBB3*** | 3.04359371702758 | 9.11002668648875 | 13.7207154457612 | 3.11714189802471e-18 | 5.19191154534996e-15 | 31.1471968139345 |
| ***ABCG4*** | 2.44930873405894 | 4.95341821647545 | 13.1394061145377 | 1.6227259648345e-17 | 2.45710215184394e-14 | 29.5585941439014 |
| ***UNC93A*** | 2.30126343737749 | 4.59368616999367 | 12.884490487967 | 3.39036770561856e-17 | 4.34384342344483e-14 | 28.8475427993215 |
| ***RFTN1*** | 2.04672267608184 | 6.73545443915594 | 12.5966561323904 | 7.86826040378141e-17 | 8.73691635235888e-14 | 28.0340012813305 |
| ***C1orf68*** | 4.32235962299866 | 4.97623513021162 | 11.7172950722719 | 1.10030231679163e-15 | 9.16331769424068e-13 | 25.4779428001913 |
| ***MT1G*** | -2.66530835824768 | 8.90083784168041 | -11.1232257913724 | 6.9181893181463e-15 | 3.60091754009515e-12 | 23.6908276789602 |
| ***ETNK2*** | -3.78249325260787 | 5.48562231017895 | -11.0208428875231 | 9.5410784540521e-15 | 4.81564250699066e-12 | 23.3779496814506 |
| ***C6orf15*** | 4.55907365795108 | 5.33286831106136 | 10.8652884126564 | 1.5589078130752e-14 | 7.4186195813087e-12 | 22.8998556339527 |
| ***PAQR8*** | -2.15181834143778 | 6.4358351451092 | -10.7075065753564 | 2.57314921621257e-14 | 1.09893264987786e-11 | 22.4115757311802 |
| ***KRT17*** | 2.7517498274232 | 11.6662867931092 | 10.4177799437914 | 6.51139956140808e-14 | 2.52218304871658e-11 | 21.5063001757402 |
| ***SIAE*** | 2.3460897269958 | 6.58845743610742 | 10.3836283845288 | 7.26949117024904e-14 | 2.75183283935609e-11 | 21.3988578664352 |
| ***MAOB*** | -2.41386555816903 | 5.24177456169648 | -10.0269362006632 | 2.31632918808145e-13 | 7.71615579133693e-11 | 20.2676035544241 |
| ***MYO5A*** | 2.08746777131415 | 6.82300249925747 | 10.014385392753 | 2.41341961533539e-13 | 7.88194453196594e-11 | 20.2274999542176 |
| ***IL1F10*** | 2.36051151260966 | 5.24992892354374 | 9.58671789474748 | 9.88924027858139e-13 | 2.74525310133419e-10 | 18.8491750275122 |
| ***SH3GL3*** | 2.42745628512091 | 5.29786553934313 | 9.54449437740439 | 1.13800444816714e-12 | 3.05719388526965e-10 | 18.7118744146256 |
| ***ALOXE3*** | 3.58800085959281 | 5.34990569057954 | 9.30380752482222 | 2.54354279794015e-12 | 6.13989113659292e-10 | 17.9251767776449 |
| ***WDR66*** | 3.38535962445437 | 6.33556452212327 | 8.94347017594976 | 8.58130323796485e-12 | 1.76457020656225e-09 | 16.7350218351613 |
| ***ADAP2*** | 3.01237885612313 | 7.57665244762842 | 8.67252264667378 | 2.16032930646897e-11 | 3.99804943650524e-09 | 15.8309378587442 |

| | | | | | | |
|---|---|---|---|---|---|---|
| ***ERP27*** | 2.79552498790453 | 5.36474401992848 | 8.66610368937332 | 2.20829664711895e-11 | 4.04191087411135e-09 | 15.8094291056565 |
| ***ROR1*** | -2.15349609595332 | 5.56648806362222 | -8.63880890726766 | 2.42455449136632e-11 | 4.38949778349972e-09 | 15.7179240132884 |
| ***TMEM45A*** | 4.7415351255839 | 8.85973680112553 | 8.02636612052767 | 2.00761879928119e-10 | 2.80999148914517e-08 | 13.6468984869663 |
| ***CEMIP*** | 2.23777422315241 | 6.03321040251497 | 7.89241399686577 | 3.20062285764225e-10 | 4.27037377180674e-08 | 13.1898707609187 |
| ***RIMS3*** | 2.12633919726468 | 6.0076119505131 | 7.89203711261137 | 3.20483142096447e-10 | 4.27037377180674e-08 | 13.1885830902361 |
| ***ASPRV1*** | 5.98813565475813 | 6.95635316941558 | 7.67290690526988 | 6.89152005814905e-10 | 8.44008515356842e-08 | 12.4383328875909 |
| ***FETUB*** | 2.33992944989181 | 4.94692345312157 | 7.65365818164294 | 7.37201497803031e-10 | 8.8336893146815e-08 | 12.372289553078 |
| ***SCNN1D*** | 2.26403674609707 | 5.10340089907077 | 7.64568906007373 | 7.58066858082203e-10 | 9.01882970586941e-08 | 12.34494092514 |
| ***SAMD5*** | -2.37433428631755 | 5.10179130343802 | -7.3296296879337 | 2.2995527685569e-09 | 2.40888999453357e-07 | 11.2576873513927 |
| ***KRT31*** | -2.25755991941901 | 5.71335790180011 | -7.29077499552444 | 2.63653151735373e-09 | 2.71074499710147e-07 | 11.1237247333907 |
| ***SLC39A6*** | 2.0294155491576 | 8.17863474694719 | 7.15329104695156 | 4.27939292880374e-09 | 4.12009067180088e-07 | 10.6493009146887 |
| ***LCE2B*** | 5.51783009239539 | 5.8169656708233 | 7.13495921398766 | 4.5651122129242e-09 | 4.34494337248374e-07 | 10.5860000114572 |
| ***COX7A1*** | -2.57924546867795 | 7.30035107689708 | -7.13308710541231 | 4.59534636041326e-09 | 4.34886869199109e-07 | 10.5795350167163 |
| ***FBN2*** | 2.16033172237205 | 5.09092524497629 | 7.09227231421447 | 5.30677049510097e-09 | 4.88340162245314e-07 | 10.4385668573534 |
| ***SUGCT*** | 2.94198791298541 | 5.72757041260871 | 7.08206961891076 | 5.50123426171185e-09 | 5.03453614632267e-07 | 10.4033221619545 |
| ***LCE1B*** | 4.91107306622118 | 5.22712494633727 | 7.0318166346913 | 6.56844392273965e-09 | 5.78857153318263e-07 | 10.2296935803891 |
| ***LOR*** | 6.4268744280096 | 5.84076121239222 | 6.97130690366756 | 8.13238101743158e-09 | 6.87578366631169e-07 | 10.0205654582558 |
| ***S100A7*** | 5.26300448131021 | 11.4293770773837 | 6.94738434430225 | 8.84908370769321e-09 | 7.40654966006724e-07 | 9.93787119658064 |
| ***CYP3A5*** | -2.96685824216288 | 5.44494195309289 | -6.89662972062915 | 1.0586113014574e-08 | 8.60108772540214e-07 | 9.76240267275975 |
| ***ARG1*** | 3.23055636211552 | 4.39003171343087 | 6.87161284073165 | 1.15640488565906e-08 | 9.26013450746984e-07 | 9.67590552719912 |
| ***DUSP14*** | 2.27940803340391 | 8.74570576701114 | 6.66161375442172 | 2.4285175457055e-08 | 1.73602524640647e-06 | 8.9497687961089 |
| ***SPRR2G*** | 6.60889821730699 | 9.71932402463414 | 6.62857599639823 | 2.7292495753152e-08 | 1.92620258162924e-06 | 8.8355500475145 |
| ***PPARGC1A*** | -2.2032537405059 | 5.06154304890019 | -6.62586222587981 | 2.75554783509468e-08 | 1.93655716208173e-06 | 8.8261685082785 |
| ***PTPRZ1*** | 2.55198906043607 | 7.86476800010851 | 6.52394068863925 | 3.95012254431515e-08 | 2.62124466526347e-06 | 8.47390644195403 |
| ***IL36G*** | 4.12660146081351 | 8.34753958512536 | 6.45303797246362 | 5.07443830447663e-08 | 3.23830821453497e-06 | 8.22897692541925 |

| | | | | | | |
|---|---|---|---|---|---|---|
| ***RNASE7*** | 3.55895081240293 | 7.14510856188076 | 6.33947341330971 | 7.57771090979407e-08 | 4.40694760603546e-06 | 7.83697679787598 |
| ***CDSN*** | 5.69105456670442 | 7.57880884256124 | 6.28210393243717 | 9.27818039439139e-08 | 5.13413198169379e-06 | 7.63912763944402 |
| ***ALOX12B*** | 4.86877695583258 | 7.17169223051124 | 6.13947558073236 | 1.53409777713622e-07 | 7.91081503900338e-06 | 7.14790689838994 |
| ***KRT16*** | 2.65520880672712 | 12.5278865458623 | 6.12088527459373 | 1.63792018436361e-07 | 8.34287418677683e-06 | 7.08396076391315 |
| ***CLDN7*** | -2.73227819491988 | 7.99834568069061 | -6.11930809618789 | 1.64704406663794e-07 | 8.36377011400044e-06 | 7.07853658949916 |
| ***APOE*** | 2.03008845431695 | 6.64633139801648 | 6.05770455403225 | 2.0459541972085e-07 | 1.01420872347336e-05 | 6.86679191194997 |
| ***CPE*** | 2.11406032752445 | 8.12548986265419 | 5.95758856335753 | 2.9093092378146e-07 | 1.34978982353872e-05 | 6.5232167701004 |
| ***S100P*** | 4.14365056001778 | 8.29427990492246 | 5.94691019195829 | 3.02052815785523e-07 | 1.39749769436768e-05 | 6.48661449259162 |
| ***C5orf46*** | 2.8126589920716 | 4.95024517393044 | 5.71955579553599 | 6.70158320875422e-07 | 2.61408828864193e-05 | 5.70963498146697 |
| ***CNTNAP2*** | 2.16763547865162 | 5.5057496053298 | 5.70652731624259 | 7.01387595369633e-07 | 2.71681669499456e-05 | 5.66525978258942 |
| ***PTN*** | -2.419005416015 | 6.37625007143804 | -5.54693795033979 | 1.22380900733066e-06 | 4.30946359959819e-05 | 5.12322696367399 |
| ***MAMDC2*** | -2.73629699233476 | 7.04465097578457 | -5.50241057421016 | 1.4288002754257e-06 | 4.84686301170886e-05 | 4.9725387139053 |
| ***HAL*** | 2.69913701908842 | 4.74270778438395 | 5.47660062399496 | 1.56284077448358e-06 | 5.21656832460891e-05 | 4.88531111498166 |
| ***KRT75*** | 4.08139450634746 | 6.93676443222859 | 5.47305326559155 | 1.58221193549351e-06 | 5.2601441112934e-05 | 4.8733293374209 |
| ***BAMBI*** | 2.89372040869592 | 5.85349954669589 | 5.24759376138931 | 3.45099096977474e-06 | 0.000102095391816284 | 4.11551922188724 |
| ***CYP4F12*** | -2.07306191379487 | 5.98102936425955 | -5.1792215316675 | 4.36591390184081e-06 | 0.000125161208173942 | 3.88728773475081 |
| ***LCE3D*** | 5.41465531939769 | 9.82567697455553 | 5.12443827462815 | 5.26858075711707e-06 | 0.000146990755595548 | 3.70500211819776 |
| ***ANXA9*** | 2.55708282047286 | 5.24736316364207 | 5.09415959843072 | 5.84411430190733e-06 | 0.000158275720020437 | 3.60448563212431 |
| ***SERPINB7*** | 3.27024735159734 | 5.99287693163596 | 5.07722073403047 | 6.19269770476381e-06 | 0.000165828895451039 | 3.54832769993977 |
| ***TF*** | -2.9269006203892 | 5.24559765023909 | -5.06975779413109 | 6.35271115047057e-06 | 0.000169568520708714 | 3.52360272946319 |
| ***CARD18*** | 2.43317122573716 | 4.84126381214546 | 5.04025024463549 | 7.02630000885408e-06 | 0.000184299295980273 | 3.42594734723664 |
| ***PTHLH*** | 2.64585243397726 | 8.34883023485729 | 4.9159557405234 | 1.07247585177569e-05 | 0.000258138118311791 | 3.0164950237256 |
| ***THY1*** | 2.01773542553415 | 7.90833152275557 | 4.91046084628397 | 1.09264647410628e-05 | 0.00026148160449302 | 2.99846745067847 |
| ***LIPG*** | 2.68166627842445 | 6.07533248724005 | 4.86413215436295 | 1.27823435307621e-05 | 0.000297350159006107 | 2.84673202667959 |
| ***IL1B*** | 2.71098257350699 | 8.28354416840212 | 4.73556814107714 | 1.97123165921885e-05 | 0.00041985721887403 | 2.42819215807515 |

| | | | | | | |
|---|---|---|---|---|---|---|
| ***IGFL2*** | 3.12591807632732 | 6.30870205171064 | 4.73351905106278 | 1.98483513318586e-05 | 0.000421676198703364 | 2.42155273819932 |
| ***WFDC5*** | 2.62214508697355 | 7.64133697070581 | 4.70427221238291 | 2.18934175691817e-05 | 0.000454118011248183 | 2.32689898313429 |
| ***DMKN*** | 2.28851684386335 | 9.59449898074596 | 4.6787823937273 | 2.38433829584811e-05 | 0.000483133073669661 | 2.24457609460169 |
| ***SPINK6*** | 4.11958144367227 | 5.91913010050111 | 4.66794674933936 | 2.47230056564596e-05 | 0.000497927910778708 | 2.20963003640944 |
| ***RPTN*** | 5.28113435021419 | 6.3059469488771 | 4.60680965406304 | 3.0315425321546e-05 | 0.000589187542772077 | 2.01301789837334 |
| ***PI3*** | 3.03820470463041 | 12.1410057361531 | 4.58426092808832 | 3.26765344845891e-05 | 0.000624868379305759 | 1.94074856212403 |
| ***KRT80*** | 2.52875688018613 | 7.56781619528042 | 4.54722015352443 | 3.69515531741609e-05 | 0.000690757653949298 | 1.82232650007613 |
| ***PRSS3*** | 2.60849349296511 | 7.48702512014752 | 4.48830376634911 | 4.49030680841653e-05 | 0.000813825355832271 | 1.63474027026748 |
| ***KRT15*** | -2.72864949018567 | 9.8790950405092 | -4.35028463645427 | 7.06491755306487e-05 | 0.00116971438135038 | 1.19919800552149 |
| ***CA2*** | 2.26610452608329 | 9.07196257992232 | 4.28735979130647 | 8.67223452618043e-05 | 0.00137174490283059 | 1.0025393957454 |
| ***ISG15*** | 2.40985969289059 | 9.13938151957253 | 4.17394482224474 | 0.00012513705723097 | 0.00185599539202051 | 0.651294629912709 |
| ***PNLIPRP3*** | 3.38976401730173 | 6.13343179925055 | 4.17029958102188 | 0.000126612915130315 | 0.00186956091703061 | 0.640076187955203 |
| ***LY6G6C*** | 2.77873417956909 | 7.84950701097736 | 4.15886002141482 | 0.000131355135822639 | 0.00192932199493992 | 0.60489959928568 |
| ***IFI27*** | 2.45483718011729 | 11.3446349931428 | 4.07063443780458 | 0.000174195005974524 | 0.00240979403613926 | 0.335128623370891 |
| ***WFDC12*** | 3.49986013318634 | 7.13881145718728 | 3.99654670242732 | 0.000220381922368241 | 0.00291092886515894 | 0.110734735978275 |
| ***BPIFC*** | 2.69223652783026 | 5.28472217315362 | 3.98315855191778 | 0.000229907243709816 | 0.00300923807512567 | 0.0704008666523244 |
| ***KRT10*** | 2.32380743009971 | 10.8174969265075 | 3.86209654322073 | 0.000336171105750222 | 0.00410965570047283 | -0.29121182416899 |
| ***KRT19*** | -2.34783271276469 | 6.24212947711988 | -3.85424911644556 | 0.000344494596308274 | 0.00418214431203398 | -0.314454469421047 |
| ***RDH12*** | 2.38218933764308 | 6.74862904467469 | 3.83725118218672 | 0.000363210698170935 | 0.00436481774079011 | -0.364714727350015 |
| ***FLG*** | 3.56147409861176 | 6.52026751010572 | 3.83565192058154 | 0.000365021230983551 | 0.00437710124064941 | -0.369437518072825 |
| ***BST2*** | 2.53233277624175 | 8.2008167962488 | 3.82614294508712 | 0.000375967383152152 | 0.00448254311652273 | -0.397497212174533 |
| ***KLK12*** | 2.43640062534941 | 7.91634388695961 | 3.82397348753443 | 0.000378508694522559 | 0.00449624971598505 | -0.40389385962127 |
| ***COMP*** | 2.34368657951952 | 6.37198742403377 | 3.81203050869332 | 0.000392797864868354 | 0.00462690327952426 | -0.439073505815057 |
| ***LCN2*** | 2.62244436463695 | 9.82334853840311 | 3.7556032995172 | 0.000467636492283654 | 0.00534221770608816 | -0.604494761256323 |
| ***KRT4*** | -3.35158685854725 | 8.90143884146788 | -3.71711728392644 | 0.000526356849392362 | 0.00588784397815929 | -0.716556399944929 |

| | | | | | | |
|---|---|---|---|---|---|---|
| ***PDZK1IP1*** | 2.07850551793216 | 8.70097258382392 | 3.68453315345034 | 0.000581553030866873 | 0.00638941113596216 | -0.81093938907765 |
| ***IFI6*** | 2.23032813627484 | 9.4762903438676 | 3.55801476881433 | 0.00085328470115944 | 0.00870852327359782 | -1.17298536564127 |
| ***S100A12*** | 2.18753293882056 | 9.22022466473405 | 3.47818602806517 | 0.00108329714244302 | 0.0104660076592407 | -1.39766802549536 |
| ***ALOX12*** | -2.19818359948987 | 6.6797485082754 | -3.11807050581898 | 0.00307391106715732 | 0.0233999372644298 | -2.37200890573904 |
| ***CES1*** | -2.45898833298829 | 6.61636028868134 | -2.98957367745711 | 0.00439595345584226 | 0.0308160777611569 | -2.70287593588458 |
| ***SPRR2D*** | 2.28519413432842 | 11.2603362721293 | 2.92472857042752 | 0.00524949960025674 | 0.0353704147823124 | -2.86624515336603 |
| ***MFAP5*** | 2.10953404294977 | 7.49037919988592 | 2.88170507306734 | 0.0058984794308285 | 0.0386183464622168 | -2.9732654727671 |
| ***CLDN17*** | 2.03295705515044 | 5.43917374510285 | 2.87625078891601 | 0.00598588409093039 | 0.0390371516908914 | -2.98675368112168 |
| ***CALML5*** | 2.27446628045827 | 8.07894174057854 | 2.7830717659261 | 0.00767760274437872 | 0.0476623746963742 | -3.21438040800305 |

Table S2 - Differentially expressed genes in early OSCC (stages I and II) in comparison to normal samples (healthy oral mucosa).

| Gene | log2 Fold Change | Average Expression | t | p-value | adjusted p-value | B |
|---|---|---|---|---|---|---|
| ***GPD1*** | -2.10475351453041 | 4.34764942208053 | -33.2213383441743 | 8.7773898739804e-35 | 7.30981028705087e-31 | 68.1639121620326 |
| ***FGF10*** | -2.07932590997188 | 3.98665990216155 | -31.8666712555666 | 5.94858505259213e-34 | 3.30265442119915e-30 | 66.3953433429412 |
| ***PI16*** | -2.77061181962147 | 5.4044534042165 | -23.7269573596629 | 3.62682105910897e-28 | 7.55104144506487e-25 | 53.7763374009944 |
| ***CASP5*** | 2.20625599770798 | 4.22671294969124 | 19.7780234000433 | 1.01901427682891e-24 | 1.02945928271534e-21 | 46.0619189449882 |
| ***GPRC5B*** | -2.70875151432749 | 5.27332415686965 | -18.5850115117831 | 1.44002519702991e-23 | 9.99377486738757e-21 | 43.466298470772 |
| ***PEBP4*** | -2.51939370864447 | 4.39126680783299 | -18.2895675525016 | 2.83097247371852e-23 | 1.57175591740852e-20 | 42.8022815740736 |
| ***ADIPOQ*** | -3.04407184300581 | 4.32248603708525 | -17.3080136117157 | 2.84111920787581e-22 | 1.35204804361084e-19 | 40.532583672363 |
| ***PRKAA2*** | -3.15052590145919 | 4.26300361290586 | -16.5646797623269 | 1.73778183770718e-21 | 7.05963275337823e-19 | 38.745939630278 |
| ***GSTM5*** | -2.7591335090674 | 4.74658663363261 | -15.817396672898 | 1.13843902090382e-20 | 3.71800790826941e-18 | 36.8879977487914 |
| ***PYGM*** | -4.11350740316356 | 4.70821972413499 | -14.8993769893942 | 1.24746612001253e-19 | 3.29806280871884e-17 | 34.5168751373412 |
| ***MYH11*** | -2.10074889632499 | 5.12377460935976 | -14.8316086304353 | 1.49433005894403e-19 | 3.77114567602603e-17 | 34.3378448146939 |
| ***SKA3*** | 2.36088074518165 | 5.79405326611842 | 14.5293219685234 | 3.36580640745257e-19 | 7.78623215590694e-17 | 33.5324358583302 |
| ***HSPB6*** | -3.21325998200715 | 4.87322560445144 | -14.0245292426281 | 1.33818519916081e-18 | 2.75170526879289e-16 | 32.162272032379 |

| | | | | | | |
|---|---|---|---|---|---|---|
| ***ADH1C*** | -2.53989769020433 | 4.30073332316218 | -14.0161961557857 | 1.36937892920528e-18 | 2.78150920059063e-16 | 32.1393858615475 |
| ***TENM2*** | 3.35149066919356 | 8.09174558368225 | 13.9780277508485 | 1.5219766606325e-18 | 3.05422207945721e-16 | 32.0344474167213 |
| ***RBP4*** | -2.09312556943033 | 4.10461929499294 | -13.5237409853892 | 5.42611713197069e-18 | 9.22218438266366e-16 | 30.7712576485129 |
| ***KRT17*** | 3.01187254123437 | 11.6662867931092 | 13.3142916429699 | 9.83477781074022e-18 | 1.48916417468808e-15 | 30.1799686069742 |
| ***NPY6R*** | -2.78669829605068 | 4.18982602637732 | -13.2295093134248 | 1.25310701072573e-17 | 1.84705755492458e-15 | 29.9390135929682 |
| ***ADH1B*** | -3.87191952456983 | 4.17835191661223 | -13.0778911054602 | 1.93704621849032e-17 | 2.68862015126456e-15 | 29.5057862321062 |
| ***C7*** | -2.32633053564481 | 4.00273245475094 | -13.039993380093 | 2.16080706576758e-17 | 2.92604898271747e-15 | 29.3970324721142 |
| ***CASQ1*** | -2.91127167204452 | 4.29836716287083 | -13.0014849006762 | 2.41511512333323e-17 | 3.21809259953906e-15 | 29.2863346014651 |
| ***CAB39L*** | -2.15728276815838 | 5.01798758255627 | -12.9086507553669 | 3.16056412879616e-17 | 4.08080280071541e-15 | 29.0186757245001 |
| ***MAOB*** | -2.65347501995397 | 5.24177456169648 | -12.8702037401581 | 3.53416685133563e-17 | 4.38392316419649e-15 | 28.9074961990569 |
| ***PADI2*** | -2.51481142650961 | 5.24952075527924 | -12.8023944791455 | 4.30591180902118e-17 | 5.2349829993472e-15 | 28.7109373317844 |
| ***ANK2*** | -2.0048281290859 | 4.50064250052564 | -12.402970591875 | 1.3947684192387e-16 | 1.49879114779612e-14 | 27.5408930562198 |
| ***NDNF*** | -3.265415649306 | 4.61978848335705 | -12.3586233970216 | 1.59119027650403e-16 | 1.66684687078309e-14 | 27.4096919295525 |
| ***CAPN6*** | -2.59251329493181 | 4.55210977407805 | -11.9096061290871 | 6.12730866644042e-16 | 5.45756433947763e-14 | 26.0666691631708 |
| ***C12orf75*** | 3.16820270798112 | 8.55302002530123 | 11.8856467172475 | 6.5891867528783e-16 | 5.74604683538958e-14 | 25.9942582281264 |
| ***CD80*** | 2.62521644824915 | 4.90239784139467 | 11.8061243478904 | 8.3910934349477e-16 | 7.24155711152792e-14 | 25.753380136371 |
| ***LDB3*** | -2.4984901895105 | 4.59365084194251 | -11.7984295807557 | 8.59005895038265e-16 | 7.33723189115761e-14 | 25.7300279482966 |
| ***NRAP*** | -3.52886296631658 | 4.59777507517331 | -11.6774412859424 | 1.24296763491756e-15 | 1.04559944076701e-13 | 25.3618243909701 |
| ***KYNU*** | 3.98730463179918 | 6.84414749578844 | 11.6604834757198 | 1.30923067572873e-15 | 1.07953198687811e-13 | 25.3100625244117 |
| ***PARP14*** | 2.51361507880285 | 6.80236053170888 | 11.6400264103453 | 1.39395424710919e-15 | 1.14372915959856e-13 | 25.2475691422021 |
| ***SH3BGRL2*** | -3.45763455432591 | 6.58993461191685 | -11.4743576949778 | 2.32087068842395e-15 | 1.80637486852286e-13 | 24.7394455973009 |
| ***SMYD1*** | -2.54749703454354 | 4.18494189107435 | -11.3550413025362 | 3.35784498687474e-15 | 2.48570071561714e-13 | 24.3712550879474 |
| ***GATM*** | -2.71517877800917 | 5.5625312855606 | -11.2793885699491 | 4.24797233055349e-15 | 3.08970424182091e-13 | 24.1368363716649 |
| ***ATP2A1*** | -2.76600173511415 | 4.4482577826067 | -11.1826022626824 | 5.74509105052253e-15 | 4.02059817384468e-13 | 23.8358412621022 |
| ***ABCA8*** | -2.90312217992166 | 4.725423402863 | -11.1731823136946 | 5.91678726094893e-15 | 4.10625035909856e-13 | 23.8064808992571 |
| ***TCEA3*** | -2.70056016117423 | 6.63687290162367 | -11.1328160664017 | 6.71344552478358e-15 | 4.6016110559998e-13 | 23.6805352118511 |
| ***DDX60L*** | 2.95348021351136 | 6.16269363947514 | 11.035550443496 | 9.10976048495951e-15 | 5.99731899752907e-13 | 23.3761878530345 |

| | | | | | | |
|---|---|---|---|---|---|---|
| ***GDPD2*** | 3.090459472698 | 4.858539801098 | 10.7632627733846 | 2.15475450989274e-14 | 1.29565310890879e-12 | 22.5176747364215 |
| ***PLIN4*** | -2.17700690612377 | 4.90464790586042 | -10.713510614352 | 2.52439818512146e-14 | 1.4963123192663e-12 | 22.3597768625163 |
| ***CASP4*** | 2.01833393781131 | 7.73064985417261 | 10.5737984351046 | 3.94434519196833e-14 | 2.24221889137967e-12 | 21.9146839371387 |
| ***TMOD1*** | -2.22513692557578 | 5.57232474043829 | -10.5697893804568 | 3.99532677744466e-14 | 2.26347492534416e-12 | 21.901875306951 |
| ***MYO1B*** | 2.55852240631578 | 8.47014060797121 | 10.4422103449179 | 6.0186359320383e-14 | 3.23842344672831e-12 | 21.4932083005761 |
| ***ANO5*** | -2.13527490241741 | 3.82590451937045 | -10.3921466729637 | 7.07242149332914e-14 | 3.76352244066742e-12 | 21.3322820790771 |
| ***PPP1R3C*** | -3.97625398495488 | 6.24931355678531 | -10.3689977520111 | 7.62103064893174e-14 | 4.00428664001915e-12 | 21.2577652785544 |
| ***PTGIS*** | -2.31225763318794 | 4.84570441005922 | -10.3515681117014 | 8.06236113973972e-14 | 4.22285179696556e-12 | 21.201614735199 |
| ***NFIX*** | -2.03797068651312 | 6.45112858030033 | -10.293422119317 | 9.73065625338565e-14 | 4.9412747120851e-12 | 21.0140202513657 |
| ***DTX3L*** | 2.15775444877099 | 8.25843511596767 | 10.2474275835176 | 1.12948710956721e-13 | 5.68360643412431e-12 | 20.8653319753296 |
| ***NFE2L3*** | 2.4546036637497 | 5.81812785558826 | 10.2221844078956 | 1.22591482737527e-13 | 6.13178299242117e-12 | 20.7836160785528 |
| ***AMPD1*** | -2.55549759216042 | 4.38398875798019 | -10.2071920809074 | 1.28708549719146e-13 | 6.41847186862904e-12 | 20.7350464467631 |
| ***INHBA*** | 3.2179059214169 | 7.49495296213354 | 10.0828188905307 | 1.92975148787423e-13 | 9.18341165200946e-12 | 20.3310580891399 |
| ***FABP3*** | -2.47477660742108 | 4.63460743731224 | -10.0512104477317 | 2.13960828816736e-13 | 9.96800620613348e-12 | 20.2280864739487 |
| ***CDH3*** | 2.609666421429 | 10.3515659739691 | 9.97804393107006 | 2.7184076650488e-13 | 1.24048761833021e-11 | 19.9892653599771 |
| ***DEPTOR*** | -2.93981147493827 | 5.65434555352311 | -9.97287812951773 | 2.76481820802944e-13 | 1.25479051969859e-11 | 19.9723793547508 |
| ***CTLA4*** | 2.21904692665849 | 5.04209205444926 | 9.89477091316366 | 3.57272258470506e-13 | 1.59965772502278e-11 | 19.7166715475025 |
| ***XDH*** | 2.68788229730149 | 6.02651755582369 | 9.86310098179463 | 3.96489494579569e-13 | 1.76104773912461e-11 | 19.6127826008811 |
| ***PPARG*** | -2.1458475348145 | 4.48386359234611 | -9.8309764684858 | 4.40723777478792e-13 | 1.93685890176431e-11 | 19.5072806889963 |
| ***APOBEC3B*** | 2.5855733846829 | 7.15927077450307 | 9.8132696040674 | 4.67205905443008e-13 | 2.0371155919002e-11 | 19.449076334132 |
| ***TTN*** | -4.06691539284744 | 5.2155187712561 | -9.76114399100326 | 5.5488584892341e-13 | 2.36372856769011e-11 | 19.2775192629528 |
| ***PLN*** | -2.65155825230494 | 4.37505645530077 | -9.74008744093059 | 5.94863044995403e-13 | 2.48945700438277e-11 | 19.2081269488488 |
| ***DLK1*** | -2.35865543153395 | 4.85876248416485 | -9.72483997499872 | 6.25618005932659e-13 | 2.58568077092168e-11 | 19.1578462200032 |
| ***MELK*** | 2.65170103310011 | 7.80291326289654 | 9.71003513121699 | 6.5701811144028e-13 | 2.67562192277489e-11 | 19.1089991397974 |
| ***PDK4*** | -3.68938149825171 | 5.90583464302015 | -9.68720407569708 | 7.08594074251563e-13 | 2.85771014545618e-11 | 19.0336203717796 |
| ***MYL3*** | -2.01632100798231 | 4.46130239624743 | -9.68407237631977 | 7.15980965577156e-13 | 2.88052631948143e-11 | 19.0232760630586 |
| ***BARX2*** | -2.09610407844274 | 5.91988461468087 | -9.64806978581773 | 8.06706597648803e-13 | 3.19916787867582e-11 | 18.9042743822338 |

| | | | | | | |
|---|---|---|---|---|---|---|
| ***ANGPTL1*** | -3.37643859336851 | 4.53545990954199 | -9.6142022576476 | 9.02644857528818e-13 | 3.50453443986014e-11 | 18.7921932843228 |
| ***CASP1*** | 2.06204603195378 | 7.87401785463025 | 9.59138139466826 | 9.73717966905218e-13 | 3.76293421270842e-11 | 18.7165956665614 |
| ***MYOM1*** | -2.51364056984839 | 4.44913809666507 | -9.5893373469864 | 9.80353600049058e-13 | 3.77980776907803e-11 | 18.709821533172 |
| ***PODN*** | -2.11285863304816 | 5.53254694538584 | -9.49323499194311 | 1.34987828868628e-12 | 4.97424176468113e-11 | 18.3907940177709 |
| ***SNX10*** | 2.36634934616769 | 6.93790613422438 | 9.47074678468298 | 1.45501248839539e-12 | 5.30299518746468e-11 | 18.3159899360967 |
| ***SLC28A3*** | 2.83640109018912 | 5.99601594336461 | 9.46266048362171 | 1.49480693180709e-12 | 5.43613630047576e-11 | 18.2890780319324 |
| ***AURKA*** | 2.08898627256098 | 6.589448343898 | 9.43318501252891 | 1.64940468406841e-12 | 5.9593241687296e-11 | 18.190919188136 |
| ***MMP12*** | 4.77190127458121 | 8.30543557759231 | 9.33849322264741 | 2.26426389905124e-12 | 7.87339864354852e-11 | 17.8749250822223 |
| ***MYPN*** | -2.30765474758847 | 4.33132487379578 | -9.28773004188934 | 2.68455315859786e-12 | 9.06976012365231e-11 | 17.7051187841325 |
| ***MMP1*** | 5.77527209776613 | 9.7704648852227 | 9.28418149137324 | 2.7167251345053e-12 | 9.15987324702839e-11 | 17.6932381577435 |
| ***CXCL1*** | 3.19771907709533 | 7.78239209693998 | 9.26984988474567 | 2.85067004408983e-12 | 9.55347288820124e-11 | 17.6452417435384 |
| ***TOP2A*** | 2.10311939604577 | 8.10403639720199 | 9.21081730376698 | 3.47651154606312e-12 | 1.13761839511252e-10 | 17.447308913302 |
| ***DDIAS*** | 2.11184602108199 | 6.2234428708306 | 9.13381117791403 | 4.50642971173402e-12 | 1.42697895967e-10 | 17.1885534833717 |
| ***MKI67*** | 2.19878289815253 | 6.74029336002639 | 9.11929534978496 | 4.73266631096853e-12 | 1.49294109991462e-10 | 17.1397074663311 |
| ***IFIH1*** | 2.8581534017242 | 6.78144741197759 | 9.01050048987508 | 6.83687991895e-12 | 2.06242081848517e-10 | 16.7729144070122 |
| ***CMYA5*** | -3.70507589021817 | 4.90452941250617 | -8.99606582299041 | 7.17950201983595e-12 | 2.15462676833131e-10 | 16.7241577713399 |
| ***LAMC2*** | 3.45692697080654 | 8.54873235040382 | 8.8663920393991 | 1.11504460098401e-11 | 3.20210049551546e-10 | 16.2852136693478 |
| ***CD274*** | 2.55941107208764 | 5.47190764626197 | 8.80278001459693 | 1.38470692755165e-11 | 3.90248368617603e-10 | 16.0692814932383 |
| ***APOL1*** | 3.06218514671182 | 7.33424397000342 | 8.71839722311021 | 1.84675299460325e-11 | 4.99342822696618e-10 | 15.782243139278 |
| ***ACTN2*** | -3.84095881961272 | 5.09151927976117 | -8.71062140143044 | 1.89647668563947e-11 | 5.11128085372346e-10 | 15.7557588853451 |
| ***ETNK2*** | -2.54982687412804 | 5.48562231017895 | -8.67486227017865 | 2.14310827235087e-11 | 5.7021743425361e-10 | 15.6338916527673 |
| ***SLC44A5*** | 2.43585811638819 | 5.50846139234266 | 8.65312672125235 | 2.30857613173625e-11 | 6.07450932862544e-10 | 15.5597590338469 |
| ***RBP1*** | 2.30151257924153 | 7.64571247957669 | 8.65238242052776 | 2.31446500304338e-11 | 6.08039891020355e-10 | 15.5572197065758 |
| ***IFI30*** | 2.70217065148016 | 9.7267663579285 | 8.6175019722014 | 2.60811593138528e-11 | 6.76647647245378e-10 | 15.4381615259514 |
| ***LRRC39*** | -2.73452309229753 | 4.17196297015904 | -8.59723124297967 | 2.79574680759449e-11 | 7.1902139236278e-10 | 15.3689203155523 |
| ***ZBTB16*** | -2.94361969771179 | 5.42610792625838 | -8.58962713134321 | 2.86959197830988e-11 | 7.34192380809975e-10 | 15.3429364803994 |
| ***PLA2G7*** | 3.2689941786459 | 7.01600668773773 | 8.58206406382327 | 2.94498913284566e-11 | 7.50026590163261e-10 | 15.3170877551044 |

| | | | | | | |
|---|---|---|---|---|---|---|
| ***TPM2*** | -2.43650038245994 | 6.94917881671078 | -8.54399219524372 | 3.35597607107783e-11 | 8.35532697158031e-10 | 15.1868900091194 |
| ***DES*** | -3.12332612112411 | 5.57057805060264 | -8.54273156774745 | 3.37053206135213e-11 | 8.37904209162404e-10 | 15.182576734567 |
| ***EIF2AK2*** | 2.2255991632582 | 8.17414000318922 | 8.53515246259606 | 3.45939639331062e-11 | 8.55438712019075e-10 | 15.1566416521718 |
| ***PRUNE2*** | -2.48062048424359 | 4.83737999859061 | -8.5223050179351 | 3.61545994617208e-11 | 8.88187328369354e-10 | 15.1126672235237 |
| ***LAMP3*** | 3.04114729858185 | 8.23239744589222 | 8.51281738939583 | 3.73524879499257e-11 | 9.13572744925055e-10 | 15.0801835610827 |
| ***OAS3*** | 2.88811041786301 | 7.54502085561267 | 8.51158054120748 | 3.75115693818501e-11 | 9.15216996618655e-10 | 15.0759482726203 |
| ***MOB3B*** | 2.04610252317816 | 6.86932591834275 | 8.49924359757571 | 3.91361870669165e-11 | 9.50222057997904e-10 | 15.0336961070579 |
| ***MMP13*** | 5.23904497540761 | 6.36886909977083 | 8.46962333014416 | 4.33323629833331e-11 | 1.0429824246393e-09 | 14.9321974810254 |
| ***PPARGC1A*** | -2.39706002387809 | 5.06154304890019 | -8.41728513756804 | 5.18867735253024e-11 | 1.23637496400205e-09 | 14.7526692484334 |
| ***CDK1*** | 2.02388202298559 | 6.53179469537793 | 8.41593404382371 | 5.21288241743031e-11 | 1.24023392324648e-09 | 14.7480317257822 |
| ***FREM2*** | -2.10887323263175 | 3.85162073623919 | -8.40448289268716 | 5.42265420744648e-11 | 1.28295068862541e-09 | 14.7087204287321 |
| ***KLHDC7B*** | 2.05190303937671 | 5.16921731649803 | 8.30112826821192 | 7.746116219622e-11 | 1.7673878322469e-09 | 14.3534212478718 |
| ***PARP12*** | 2.30204459594393 | 8.5599951114481 | 8.28144497923448 | 8.29132325358674e-11 | 1.87127750828917e-09 | 14.2856591396776 |
| ***TRDN*** | -3.80591151447394 | 4.27122912065231 | -8.24608276404679 | 9.36978544119882e-11 | 2.08084195078143e-09 | 14.1638440666964 |
| ***PTPN22*** | 2.17450670393741 | 4.90783481912885 | 8.22602817976388 | 1.00430847541315e-10 | 2.21559761145449e-09 | 14.0947173681694 |
| ***ARHGEF26*** | -2.05952275039686 | 4.51141011070394 | -8.21133160769856 | 1.05672483827877e-10 | 2.31589590873306e-09 | 14.0440397987176 |
| ***CDC6*** | 2.31516617535094 | 5.60455612334508 | 8.20545381215213 | 1.07845179932307e-10 | 2.3573088148983e-09 | 14.0237670475865 |
| ***TNC*** | 2.76288770630918 | 8.59328585705563 | 8.18427656054816 | 1.16053042270951e-10 | 2.50821853751438e-09 | 13.9507042444309 |
| ***FRZB*** | -2.18769217806192 | 5.08181173093489 | -8.1736366526328 | 1.20411383676598e-10 | 2.59117830299408e-09 | 13.9139832291679 |
| ***PTPRZ1*** | 2.7380617110053 | 7.86476800010851 | 8.17315382639354 | 1.20613022881716e-10 | 2.59216839886177e-09 | 13.9123166735169 |
| ***S100A7A*** | 4.22514470840923 | 5.80529484563635 | 8.14045915694166 | 1.35088612902671e-10 | 2.86994379656492e-09 | 13.7994254536946 |
| ***C16orf54*** | 2.06068775104996 | 5.85933092914003 | 8.1268559654912 | 1.41615373214113e-10 | 2.97272665422759e-09 | 13.7524320045684 |
| ***LMOD3*** | -2.76609935057455 | 5.05447855637064 | -8.12666058415108 | 1.41711393098986e-10 | 2.97272665422759e-09 | 13.7517569442377 |
| ***CD86*** | 2.05008802055053 | 6.11509753334005 | 8.06590084720333 | 1.7498864679361e-10 | 3.62063465962033e-09 | 13.541693954588 |
| ***MYH7*** | -3.69921931313724 | 5.07423292511225 | -8.05038457238082 | 1.84681357741516e-10 | 3.77429778471496e-09 | 13.4880082396116 |
| ***GPC3*** | -2.36104423673793 | 4.93965880028018 | -8.04584208940992 | 1.87619909319099e-10 | 3.82028998730918e-09 | 13.4722882639355 |
| ***ITGA3*** | 2.08636671459611 | 7.92020826967946 | 8.01935262280988 | 2.05718042416902e-10 | 4.11831696453836e-09 | 13.3805890832047 |

| | | | | | | |
|---|---|---|---|---|---|---|
| ***PIM2*** | 2.71550250609548 | 6.15255837077442 | 7.92213625190116 | 2.88563897130481e-10 | 5.54362199608454e-09 | 13.0436515246536 |
| ***CKS2*** | 2.00678603315408 | 9.3096457419611 | 7.91746763342237 | 2.93296844371785e-10 | 5.61511751707637e-09 | 13.0274553645974 |
| ***NEB*** | -4.11714386970485 | 5.44604642890316 | -7.91204411673921 | 2.98893229522187e-10 | 5.69607051592854e-09 | 13.008638616508 |
| ***GULP1*** | -2.25610951156113 | 4.99119458991965 | -7.8977680107018 | 3.14143291050352e-10 | 5.94587574515303e-09 | 12.9590992109171 |
| ***SYNPO2*** | -2.62179561332833 | 5.84008862291203 | -7.87778429283991 | 3.36814141273135e-10 | 6.33895631304559e-09 | 12.8897325765783 |
| ***ABI3BP*** | -3.00542241814722 | 6.05831443020489 | -7.8345260517002 | 3.91687592215771e-10 | 7.21118494155667e-09 | 12.7394930100659 |
| ***TIGIT*** | 2.00571730755604 | 4.85702950423234 | 7.75714394942291 | 5.13254999689576e-10 | 9.19223147831137e-09 | 12.4704643535342 |
| ***MT1M*** | -2.46005207885799 | 5.95123002498895 | -7.68318408923879 | 6.64799458692446e-10 | 1.15704281964278e-08 | 12.2130230189094 |
| ***CEP55*** | 2.25808752185759 | 7.86955438695579 | 7.67198379183674 | 6.91382792283696e-10 | 1.19396043286396e-08 | 12.1740115947634 |
| ***IRF1*** | 2.37873048131667 | 7.54685920089794 | 7.61645235371845 | 8.39829967752213e-10 | 1.41010160714525e-08 | 11.9804985184376 |
| ***KRT36*** | -2.16548889486194 | 3.92189605447652 | -7.61179597524295 | 8.53646505432314e-10 | 1.42754379462657e-08 | 11.9642653430249 |
| ***ANLN*** | 2.53070316450021 | 8.0352209481484 | 7.58617739011565 | 9.33850713964746e-10 | 1.54460948280008e-08 | 11.8749348190367 |
| ***OAS2*** | 2.64106928297935 | 7.27745008859287 | 7.5851962946527 | 9.37068426933067e-10 | 1.54839401974178e-08 | 11.8715131837243 |
| ***DDX58*** | 2.9991071808237 | 7.34897387285247 | 7.573470832797 | 9.76397613847929e-10 | 1.60096768530051e-08 | 11.8306164168217 |
| ***GABRE*** | 2.27028891099584 | 7.24267307731222 | 7.55457448918098 | 1.04329904570303e-09 | 1.69038802580055e-08 | 11.7646954146779 |
| ***SAMD9*** | 2.59046082968485 | 9.79844370486239 | 7.55064823927513 | 1.05776738525768e-09 | 1.71050228823805e-08 | 11.7509964453261 |
| ***PARP9*** | 2.54661765794615 | 7.73402490299945 | 7.51307316902825 | 1.20684454811286e-09 | 1.92724858996814e-08 | 11.619860094314 |
| ***DDX60*** | 2.88909240842574 | 8.06487254992665 | 7.49543687008577 | 1.28392216643401e-09 | 2.02893810285815e-08 | 11.55828888577 |
| ***LIFR*** | -2.07867802473201 | 5.53535870825693 | -7.459927227179 | 1.45444006536926e-09 | 2.27466232195214e-08 | 11.4342801597935 |
| ***IL12RB2*** | 2.55805340528585 | 5.71180489834863 | 7.45528831420665 | 1.47833496895418e-09 | 2.30496885398445e-08 | 11.4180761722374 |
| ***CRYAB*** | -3.44818036180387 | 9.55597766534151 | -7.45509219552333 | 1.47935380938363e-09 | 2.30496885398445e-08 | 11.4173910999173 |
| ***ICOS*** | 2.89934559749099 | 5.1555129360299 | 7.40066356062144 | 1.79118872338857e-09 | 2.7295552952205e-08 | 11.2272070944387 |
| ***IFI44*** | 3.53365593200439 | 7.56919621725605 | 7.37090234720585 | 1.98878899181345e-09 | 2.99234593022989e-08 | 11.1231701985625 |
| ***PLAU*** | 2.39260215278159 | 8.57020412917519 | 7.36231018575495 | 2.0498060225024e-09 | 3.075817037009e-08 | 11.0931287131013 |
| ***PDGFD*** | -2.11876665571311 | 5.07167879564174 | -7.35958521643061 | 2.06954705854176e-09 | 3.10169231809949e-08 | 11.0836006577809 |
| ***LYN*** | 2.12204466091415 | 7.98057824877848 | 7.34728939610554 | 2.16102418779578e-09 | 3.21950079355336e-08 | 11.0406043523236 |
| ***CILP*** | -3.13924489649823 | 4.9482775836485 | -7.31706860833915 | 2.4034774253596e-09 | 3.52707665169952e-08 | 10.934906867372 |

| | | | | | | |
|---|---|---|---|---|---|---|
| ***STAC3*** | -2.06644601225098 | 5.63630149986437 | -7.30719162941241 | 2.48849473472808e-09 | 3.64862397021399e-08 | 10.9003558609317 |
| ***CKMT2*** | -2.63612806022477 | 4.83515373184243 | -7.2997305264264 | 2.55471217441114e-09 | 3.74241741222444e-08 | 10.8742539593745 |
| ***GBP5*** | 4.44427558551104 | 6.65113499294338 | 7.27311211408585 | 2.80568991668329e-09 | 4.04252346472983e-08 | 10.7811188832862 |
| ***ZNF667-AS1*** | -2.128332763259 | 4.87188084099555 | -7.26692838409712 | 2.86745048786017e-09 | 4.11910949710106e-08 | 10.7594797626951 |
| ***DLX3*** | 2.02199058886459 | 6.65732084813927 | 7.24781007715923 | 3.06715728561903e-09 | 4.37759826471898e-08 | 10.6925712299758 |
| ***IL24*** | 3.11083394567403 | 6.37862107758704 | 7.24495761860451 | 3.09812694593645e-09 | 4.41423459465504e-08 | 10.6825876125792 |
| ***HCAR3*** | 2.34721816686528 | 7.13722736190716 | 7.22661111132876 | 3.30495448015791e-09 | 4.68089471271345e-08 | 10.6183697158508 |
| ***DTL*** | 2.01089470192016 | 7.04175576090158 | 7.2012665654535 | 3.61365050139691e-09 | 5.06214993702833e-08 | 10.5296429313087 |
| ***ANKRD1*** | -3.1133407690426 | 4.64863450754387 | -7.17481861817168 | 3.96666699125246e-09 | 5.46927197072028e-08 | 10.4370373034396 |
| ***OGN*** | -3.5002274272681 | 4.73044310481258 | -7.15619445612595 | 4.23581480370173e-09 | 5.76874336634964e-08 | 10.371816932978 |
| ***CXCL10*** | 6.14648033692434 | 8.7910234744076 | 7.09868888584299 | 5.18801504610774e-09 | 6.89087548707899e-08 | 10.1703940127992 |
| ***LYVE1*** | -2.42733252286539 | 5.60923414561174 | -7.0490734128766 | 6.18040210479997e-09 | 8.01766541151674e-08 | 9.99656297639755 |
| ***KLHL41*** | -4.75403959098151 | 5.34685620455355 | -7.00050144619454 | 7.33602553905835e-09 | 9.34880194174108e-08 | 9.82635653937075 |
| ***ENO3*** | -2.19069445226263 | 4.90410063925229 | -6.99871160887869 | 7.38251800658439e-09 | 9.40085779187077e-08 | 9.820084083315 |
| ***TNFAIP3*** | 2.09129164435193 | 9.09919551548545 | 6.99621803382357 | 7.44778316097835e-09 | 9.46226364067547e-08 | 9.81134533856448 |
| ***STAT1*** | 2.51159486194126 | 9.62514058348407 | 6.98040846550729 | 7.8752368618293e-09 | 9.93711705838097e-08 | 9.75593932134752 |
| ***CXCL11*** | 5.67481804273164 | 6.54242117905535 | 6.9725511179362 | 8.09673826020155e-09 | 1.01780582990126e-07 | 9.72840177123022 |
| ***PI3*** | 3.95006704815152 | 12.1410057361531 | 6.95940337470203 | 8.48142485775013e-09 | 1.06260971810013e-07 | 9.68232198915522 |
| ***CHRDL1*** | -3.04124376591616 | 5.52064245561558 | -6.95156847513116 | 8.71931536694209e-09 | 1.08818379546713e-07 | 9.65486192252815 |
| ***GLDC*** | 2.36227232871455 | 5.35494741502665 | 6.93445267395785 | 9.26250776389669e-09 | 1.14703590569118e-07 | 9.59487251146365 |
| ***PDLIM3*** | -2.9010253827232 | 5.89916981439912 | -6.88384678539439 | 1.10750000527882e-08 | 1.32879828091282e-07 | 9.41749618543559 |
| ***APOBEC3G*** | 2.0158691042325 | 5.90001307179194 | 6.88368585333462 | 1.10812969031455e-08 | 1.32879828091282e-07 | 9.41693210009474 |
| ***XAF1*** | 2.76995869579837 | 8.34100141913849 | 6.86820832602089 | 1.17039491306181e-08 | 1.39542574602416e-07 | 9.36268149618174 |
| ***IDO1*** | 4.43794527276434 | 6.41043964986764 | 6.83781812332654 | 1.30302865317734e-08 | 1.53809242001037e-07 | 9.2561604284125 |
| ***CFB*** | 2.18614160149299 | 7.27443210371463 | 6.80044951582045 | 1.48692096444557e-08 | 1.72706803234348e-07 | 9.12518243111272 |
| ***SLAMF7*** | 2.4115556417846 | 6.40880267421816 | 6.78167679140411 | 1.58888656030692e-08 | 1.83526314483163e-07 | 9.05938600356695 |
| ***MGST1*** | -3.1253701447685 | 5.70340938323627 | -6.77451132898459 | 1.62962617358911e-08 | 1.8732266078192e-07 | 9.034272401777 |

| | | | | | | |
|---|---|---|---|---|---|---|
| ***WIF1*** | -2.38913798898364 | 4.4327768748711 | -6.76455295566896 | 1.68798660287342e-08 | 1.93363857341538e-07 | 8.99937079388636 |
| ***CASP14*** | 3.51666486694306 | 5.02789351268293 | 6.7508245858958 | 1.77188501579668e-08 | 2.01405159218579e-07 | 8.95125761072293 |
| ***USP18*** | 2.69481412144982 | 6.55300863587281 | 6.74456799269614 | 1.81149283955554e-08 | 2.05532866046574e-07 | 8.92933097442297 |
| ***MMP3*** | 3.72671587164298 | 8.09973724881271 | 6.67637368958414 | 2.30509896641929e-08 | 2.52923111888535e-07 | 8.69036819932537 |
| ***CXCL13*** | 3.44340154693718 | 7.29997954906581 | 6.65246289223342 | 2.50833092513468e-08 | 2.72351759380986e-07 | 8.60659660166476 |
| ***IL2RB*** | 2.57619511110226 | 6.68319073882314 | 6.63510384200223 | 2.66701357423316e-08 | 2.86592116725339e-07 | 8.54578513767227 |
| ***LDOC1*** | -2.47560791723524 | 6.0102367819243 | -6.6215417564695 | 2.7979398239043e-08 | 2.99117366540116e-07 | 8.4982788965138 |
| ***FHL1*** | -3.1768406620769 | 7.93114278062943 | -6.57021725354818 | 3.35429337919196e-08 | 3.52264252987524e-07 | 8.31853027011315 |
| ***CCL18*** | 3.16000906102123 | 7.9842583407813 | 6.46099032536883 | 4.9338866015163e-08 | 4.9775175793371e-07 | 7.9362210917024 |
| ***IFIT2*** | 3.15644671382227 | 6.92881319003976 | 6.45239812512246 | 5.08591942950258e-08 | 5.11540302039824e-07 | 7.90616270975056 |
| ***PAX9*** | -2.76586710442697 | 6.18159246329995 | -6.44427767862341 | 5.23390277780374e-08 | 5.24839763197465e-07 | 7.87775700979689 |
| ***UBD*** | 3.27779117634138 | 7.06719086498024 | 6.4359417135772 | 5.39028735164346e-08 | 5.37930653858439e-07 | 7.84859982271502 |
| ***MYBPC1*** | -4.3297894754747 | 5.51114037326454 | -6.4191229220256 | 5.72015894565323e-08 | 5.67450669438953e-07 | 7.78977939753093 |
| ***STATH*** | -3.89818410736826 | 4.12173564771416 | -6.40053980869771 | 6.10812130177271e-08 | 6.01128533119587e-07 | 7.72480091084709 |
| ***DCT*** | -2.54323314127458 | 4.40464388651133 | -6.39650956000433 | 6.1956682610662e-08 | 6.07386995622829e-07 | 7.71071033544124 |
| ***CRISP3*** | -6.14523633697503 | 5.6163693943407 | -6.37736318973669 | 6.62899422787478e-08 | 6.43429649530783e-07 | 7.64377955289409 |
| ***BPGM*** | 2.05545709470369 | 8.0233527847173 | 6.35321238412364 | 7.21898216876748e-08 | 6.94623726187124e-07 | 7.55937617372796 |
| ***TAP1*** | 2.26388226568018 | 10.1641663515317 | 6.28454968724508 | 9.19846303698153e-08 | 8.62181206212517e-07 | 7.31955342283632 |
| ***IL1B*** | 3.06265796787491 | 8.28354416840212 | 6.24682166076255 | 1.05078917239094e-07 | 9.7017430461993e-07 | 7.18787653591996 |
| ***MMP10*** | 4.88584808224441 | 7.78996212898902 | 6.24235098358852 | 1.06748912264606e-07 | 9.82971610070448e-07 | 7.17227810781421 |
| ***IL36G*** | 3.41684294402861 | 8.34753958512536 | 6.23895646149571 | 1.08034571526466e-07 | 9.93059505157184e-07 | 7.16043516243041 |
| ***SMPX*** | -4.13329039706096 | 4.98527522977155 | -6.21245220660464 | 1.18618552917378e-07 | 1.07829678746949e-06 | 7.06798789438189 |
| ***ERAP2*** | 2.42826091352747 | 6.99121968412214 | 6.20530525543106 | 1.21645467858722e-07 | 1.10055780155072e-06 | 7.04306598909366 |
| ***TXLNB*** | -2.14593018782709 | 4.90135623703117 | -6.20103545676777 | 1.23490457253476e-07 | 1.11482767263626e-06 | 7.02817831422277 |
| ***EPSTI1*** | 2.90637341598366 | 7.96287895812683 | 6.17139469310445 | 1.37090670128315e-07 | 1.22260554306044e-06 | 6.924858326096 |
| ***FPR3*** | 2.3216945893293 | 7.11678405335903 | 6.14958693784332 | 1.4804094088847e-07 | 1.31157974012678e-06 | 6.84887628058029 |
| ***FAM3B*** | -3.20833676670653 | 5.58659857047138 | -6.12795267678875 | 1.5976502230141e-07 | 1.40498744004872e-06 | 6.77352844843731 |

| | | | | | | |
|---|---|---|---|---|---|---|
| ***CXCL8*** | 3.07136565911674 | 8.28916215143004 | 6.09524514816846 | 1.79269460138182e-07 | 1.55435300784048e-06 | 6.65967314421294 |
| ***AIM2*** | 2.8960202877438 | 6.73598107506574 | 6.0895269698613 | 1.82915198605455e-07 | 1.58266781712855e-06 | 6.63977555589029 |
| ***RNASE7*** | 2.92312995774393 | 7.14510856188076 | 6.07987384483766 | 1.89238084307181e-07 | 1.62733525986e-06 | 6.60619066408911 |
| ***RRAGD*** | -2.41085480369318 | 7.55610374551101 | -6.07970930335996 | 1.8934772656152e-07 | 1.62733525986e-06 | 6.60561825243956 |
| ***ABCA12*** | 3.04084119754192 | 6.39776726160443 | 6.06308467439476 | 2.00758053326945e-07 | 1.71390370897673e-06 | 6.54779392250336 |
| ***FLNC*** | -2.37210728699231 | 6.41455647898699 | -6.05492225501381 | 2.06608460351794e-07 | 1.75485492892376e-06 | 6.5194103988682 |
| ***SLCO1B3*** | 3.37774099690223 | 4.97365905852374 | 6.0133421329525 | 2.39152166747775e-07 | 1.98966957510037e-06 | 6.37489883038023 |
| ***RSAD2*** | 3.75271271230119 | 7.06701184140673 | 6.00773343062525 | 2.43915970741866e-07 | 2.02022098889932e-06 | 6.35541588272347 |
| ***CMPK2*** | 2.52162637549764 | 7.43913324191089 | 5.97854416353215 | 2.70275921617773e-07 | 2.20564220992926e-06 | 6.25406118754907 |
| ***OLR1*** | 2.29122037381127 | 5.36535198365359 | 5.96922642025428 | 2.7927336952649e-07 | 2.26244029320682e-06 | 6.22172136329204 |
| ***MGP*** | -2.70437855226224 | 7.56543062693426 | -5.96537926999179 | 2.83074657290362e-07 | 2.287671757316e-06 | 6.20837083216212 |
| ***RYR1*** | -2.34557668450804 | 6.15579506836231 | -5.9642584463537 | 2.84191774369207e-07 | 2.29558593302304e-06 | 6.20448153583964 |
| ***CASQ2*** | -3.46169376472014 | 5.20907556393222 | -5.94617892816188 | 3.02829749381496e-07 | 2.43211285492052e-06 | 6.14175945174637 |
| ***MX2*** | 2.91115308002722 | 8.10605263904007 | 5.91398083101102 | 3.39083831946166e-07 | 2.68175702986483e-06 | 6.0301256668501 |
| ***MYOZ2*** | -2.67870665268935 | 4.27777708646794 | -5.88641781382487 | 3.73528097716296e-07 | 2.92425515580233e-06 | 5.93463465000426 |
| ***GXYLT2*** | -2.26122171701819 | 6.41501507457923 | -5.88135945998758 | 3.80217146043278e-07 | 2.97040186890096e-06 | 5.91711763909013 |
| ***HCP5*** | 2.10336096407039 | 7.68202645366022 | 5.88010744438997 | 3.81891091621298e-07 | 2.97928712976316e-06 | 5.91278228827308 |
| ***CD38*** | 2.3081916382523 | 5.95900530501394 | 5.8612582081044 | 4.07996570993114e-07 | 3.15194382488929e-06 | 5.84753064295831 |
| ***SAMD9L*** | 2.39103761007093 | 7.71955187804554 | 5.85387288151225 | 4.18702176117538e-07 | 3.22567226892401e-06 | 5.82197344516555 |
| ***IFI35*** | 2.29346602221058 | 7.89439687389463 | 5.8320694458528 | 4.51964778467473e-07 | 3.45001161785253e-06 | 5.74655222705667 |
| ***S100P*** | 3.47981184504035 | 8.29427990492246 | 5.83148635110872 | 4.52889495337606e-07 | 3.45548668545266e-06 | 5.74453584905425 |
| ***IL1A*** | 2.65082298467713 | 7.81560984183177 | 5.81720383117825 | 4.76136443754539e-07 | 3.61462322031927e-06 | 5.69515640770371 |
| ***TRIM22*** | 2.15479371784367 | 9.06382948124515 | 5.8114068270555 | 4.85906916448301e-07 | 3.6804300138076e-06 | 5.67511997790157 |
| ***THBS4*** | -3.44544239435144 | 6.29970912017982 | -5.8016221413973 | 5.02852442783801e-07 | 3.79497520933711e-06 | 5.64130837390718 |
| ***MAL*** | -5.02152625018916 | 9.51705410764117 | -5.79593786848648 | 5.1296507085327e-07 | 3.86254349915554e-06 | 5.62167043630481 |
| ***CRNN*** | -5.23620363486683 | 9.59167088048632 | -5.79557202455063 | 5.13622806225954e-07 | 3.86574851355603e-06 | 5.62040663653414 |
| ***CDSN*** | 4.49227934863046 | 7.57880884256124 | 5.79021069438374 | 5.23358408624802e-07 | 3.92660254687148e-06 | 5.60188760302175 |

| | | | | | | |
|---|---|---|---|---|---|---|
| ***GPX3*** | -2.41093543290722 | 9.51194152700226 | -5.77294109002883 | 5.55982047000926e-07 | 4.1315345725156e-06 | 5.54225523604151 |
| ***MYF6*** | -2.77435927631678 | 4.72557257475065 | -5.76795357016729 | 5.65774425783709e-07 | 4.19402063081848e-06 | 5.52503896609746 |
| ***MYL2*** | -4.58045767410217 | 5.79577615043568 | -5.76794067056745 | 5.65799973429942e-07 | 4.19402063081848e-06 | 5.52499444173024 |
| ***IFIT3*** | 2.86656079851046 | 7.49953730825425 | 5.73841087049965 | 6.27392729106995e-07 | 4.60751908994978e-06 | 5.42311535271769 |
| ***TGFBR3*** | -2.1399961383356 | 7.2890021960777 | -5.7362688390585 | 6.32111278163448e-07 | 4.6360393875343e-06 | 5.41572888759445 |
| ***XIRP2*** | -3.68998815005695 | 4.78168660621679 | -5.71681878442871 | 6.76602248717282e-07 | 4.92762879520553e-06 | 5.34868130367583 |
| ***MYOT*** | -3.78544847109899 | 5.32366361309682 | -5.71219237251555 | 6.87634725672653e-07 | 4.9970523520086e-06 | 5.33273942919621 |
| ***TREM2*** | 2.04785061434833 | 6.36782195022641 | 5.65587238496617 | 8.37152866246068e-07 | 5.94103883263507e-06 | 5.13886444604089 |
| ***TNNT1*** | -3.11832937963005 | 7.06727421683125 | -5.6507928804681 | 8.52128717852656e-07 | 6.0191076864096e-06 | 5.12139694512801 |
| ***PTHLH*** | 2.60135792340075 | 8.34883023485729 | 5.64361919210051 | 8.73732781508015e-07 | 6.14305327513613e-06 | 5.09673314605178 |
| ***TYRP1*** | -3.08368991211002 | 4.8778680545418 | -5.60924956854115 | 9.85019967921357e-07 | 6.830346621856e-06 | 4.97865341794664 |
| ***GBP1*** | 2.27221187636029 | 9.12987510573345 | 5.54685451889026 | 1.22416441643878e-06 | 8.28512089402859e-06 | 4.76466733257762 |
| ***PSMB9*** | 2.23240798356274 | 7.98831690655825 | 5.54065758374573 | 1.25085079319369e-06 | 8.42993921944354e-06 | 4.74344219143896 |
| ***CA3*** | -2.48121813291349 | 4.90501225117326 | -5.51925385168407 | 1.34753709645752e-06 | 9.01675541649613e-06 | 4.67017149992306 |
| ***MMP9*** | 2.38197096514445 | 8.10005790349088 | 5.51634309841413 | 1.36124553068521e-06 | 9.09098057702198e-06 | 4.66021196725879 |
| ***KRT4*** | -4.25580547049899 | 8.90143884146788 | -5.51128409960198 | 1.38540098148854e-06 | 9.2374854874592e-06 | 4.6429046573037 |
| ***KRT16*** | 2.04157085073444 | 12.5278865458623 | 5.49534877207907 | 1.46429921598795e-06 | 9.70101191167326e-06 | 4.58841125231501 |
| ***NELL2*** | 3.56789068997647 | 7.24984317010403 | 5.47250626732569 | 1.58522003898089e-06 | 1.03868705622603e-05 | 4.51035896745288 |
| ***CSAG3*** | 2.43850053653988 | 5.44073629830312 | 5.4533409003331 | 1.6942697855561e-06 | 1.09932830339783e-05 | 4.44492810713463 |
| ***CD2*** | 2.55419851114771 | 7.25006827379438 | 5.4175693144561 | 1.91805800236773e-06 | 1.22449881515665e-05 | 4.32294514541747 |
| ***MYOZ1*** | -3.19586621147189 | 5.0970411631291 | -5.38530288770467 | 2.14489151358482e-06 | 1.34761648624175e-05 | 4.21307752787865 |
| ***DKK 1,00*** | 3.53127089158366 | 6.25058424805927 | 5.37633743775322 | 2.21250273100243e-06 | 1.38279345169142e-05 | 4.18257805027755 |
| ***ASB5*** | -3.16454546870852 | 4.96918723483117 | -5.35711203271354 | 2.36468029245749e-06 | 1.46253676016234e-05 | 4.11721710926387 |
| ***ACTA1*** | -4.71016029724701 | 6.94131790613176 | -5.34210301118929 | 2.49064293475344e-06 | 1.5307803956182e-05 | 4.06623078333452 |
| ***CTSV*** | 2.46898257437361 | 8.93855925369403 | 5.3287128175023 | 2.60860939008178e-06 | 1.59387373445349e-05 | 4.02077377423396 |
| ***IFI44L*** | 3.2288233479564 | 7.61798936144832 | 5.32090451262419 | 2.67993383777113e-06 | 1.62908678839109e-05 | 3.99427938316309 |
| ***SLC6A14*** | 2.95849968977243 | 6.67922110124174 | 5.28393815827238 | 3.04466018438042e-06 | 1.82002764524688e-05 | 3.86898317221553 |

| | | | | | | |
|---|---|---|---|---|---|---|
| ***MAGEA3*** | 3.51385984048467 | 5.98496149320308 | 5.2476245987057 | 3.45062444672797e-06 | 2.02905618814303e-05 | 3.74612071180426 |
| ***SLN*** | -4.04959056337463 | 6.07360913404399 | -5.24544209613593 | 3.47666024762562e-06 | 2.04042470346907e-05 | 3.73874361008982 |
| ***TMPRSS11B*** | -4.81981867029929 | 6.76354360130201 | -5.23932735418346 | 3.55064350227747e-06 | 2.07871768625425e-05 | 3.71807945756439 |
| ***WFDC5*** | 2.49719552096466 | 7.64133697070581 | 5.2312264275956 | 3.65105867998709e-06 | 2.12555167332628e-05 | 3.6907131262844 |
| ***CA2*** | 2.36519910199402 | 9.07196257992232 | 5.22507998371002 | 3.72911461010494e-06 | 2.16343200786861e-05 | 3.66995696928339 |
| ***MX1*** | 2.54944051735359 | 10.2294444815646 | 5.20536695057316 | 3.99075046713997e-06 | 2.2889097720621e-05 | 3.60343189597026 |
| ***CD79A*** | 2.1177615931358 | 5.75469283010934 | 5.19714568185865 | 4.10514972436897e-06 | 2.34805541926818e-05 | 3.57570806825986 |
| ***PPBP*** | 2.04879737774742 | 5.18615702068424 | 5.19445376491249 | 4.14330722243223e-06 | 2.36500771407921e-05 | 3.56663297978654 |
| ***IL36RN*** | 2.181171539002 | 7.44725036231257 | 5.18616222085543 | 4.26305348083344e-06 | 2.42587696538304e-05 | 3.53868837284771 |
| ***GNLY*** | 2.47154291896151 | 6.19575219821934 | 5.18432775398437 | 4.29000508075583e-06 | 2.4378821093507e-05 | 3.53250742178914 |
| ***SPP1*** | 2.32744281627207 | 6.57258970629775 | 5.1539265871805 | 4.76195186686192e-06 | 2.67234064334408e-05 | 3.43016433532135 |
| ***CPA4*** | 2.34437024694064 | 7.59644736968855 | 5.14892007573137 | 4.84444267047948e-06 | 2.71041441449467e-05 | 3.41332655794246 |
| ***IFI6*** | 2.74706764843894 | 9.4762903438676 | 5.11709621126593 | 5.40277814219615e-06 | 2.98173203235318e-05 | 3.30640657887157 |
| ***ITK*** | 2.17606067961102 | 6.23054714959727 | 5.10153327566436 | 5.69848755572008e-06 | 3.12217133973927e-05 | 3.25418880702048 |
| ***MYH2*** | -4.26136173732711 | 5.52137722238742 | -5.08849855332106 | 5.95840310005495e-06 | 3.24642335736065e-05 | 3.21048962379002 |
| ***APOD*** | -2.75862707823498 | 7.99542625476502 | -5.07302106766517 | 6.28224747756321e-06 | 3.39861918516171e-05 | 3.15864383566455 |
| ***EGFL6*** | 2.03709339635392 | 7.02476026877395 | 5.0582494691202 | 6.60748742626816e-06 | 3.55472579366674e-05 | 3.10920641566114 |
| ***LRRN1*** | -2.04341709087972 | 4.49099385170702 | -5.05460116375238 | 6.69033910408399e-06 | 3.59697508449396e-05 | 3.09700295426455 |
| ***S100A7*** | 3.24748941364646 | 11.4293770773837 | 5.0055358347744 | 7.90929154714446e-06 | 4.15968298103057e-05 | 2.93314112998795 |
| ***IFIT1*** | 3.26336689926335 | 8.77121842859815 | 5.00424861688489 | 7.94405464632316e-06 | 4.1740118040744e-05 | 2.92884885614928 |
| ***CKM*** | -3.70948750801132 | 5.62319305881382 | -4.96129610024353 | 9.19446456532467e-06 | 4.73686983606705e-05 | 2.78582020495238 |
| ***HERC5*** | 2.00051952579304 | 7.16267478852758 | 4.9589409802962 | 9.26836898391943e-06 | 4.77051773164901e-05 | 2.77798907842223 |
| ***CCL20*** | 2.18919981198299 | 7.0215256134595 | 4.91001618390613 | 1.0942949719172e-05 | 5.50651874690421e-05 | 2.61557719464488 |
| ***MYL1*** | -4.45559996759489 | 6.03132641182728 | -4.85418127845055 | 1.32196969151926e-05 | 6.50095281427365e-05 | 2.43087276858192 |
| ***MB*** | -2.72791423723085 | 5.69017103478786 | -4.83349983509578 | 1.41762766159995e-05 | 6.92637322722464e-05 | 2.36263760949648 |
| ***CCL5*** | 2.5559054621089 | 8.29374316744032 | 4.82935305302699 | 1.43761110554217e-05 | 7.00551508891468e-05 | 2.34896788380286 |
| ***BST2*** | 2.73107650293963 | 8.2008167962488 | 4.81825217263832 | 1.49247941467455e-05 | 7.24111189362635e-05 | 2.31239397825578 |

| | | | | | | |
|---|---|---|---|---|---|---|
| ***SELL*** | 2.18425486607339 | 6.40756685543264 | 4.81793918102868 | 1.49405590554821e-05 | 7.24454007650973e-05 | 2.31136318813935 |
| ***TNNT3*** | -2.41998206445139 | 6.20846477089669 | -4.79940505109615 | 1.59038582044772e-05 | 7.6537030411376e-05 | 2.25036507345584 |
| ***CSRP3*** | -3.19850501016651 | 5.14578982577969 | -4.79178183277428 | 1.6317553728764e-05 | 7.82512763824907e-05 | 2.22529978516077 |
| ***OAS1*** | 2.10543935292424 | 8.32822203022629 | 4.7486313185086 | 1.88663473544255e-05 | 8.86601765068433e-05 | 2.08368412393164 |
| ***KRT76*** | -2.18429701921149 | 4.91899507720787 | -4.71228825767051 | 2.13131851938482e-05 | 9.84995595418246e-05 | 1.96476476729163 |
| ***SERPINB7*** | 2.58514310966009 | 5.99287693163596 | 4.68646391408452 | 2.32384761660793e-05 | 0.000106276787210933 | 1.8804658440899 |
| ***KRT10*** | 2.41007499076638 | 10.8174969265075 | 4.67701530423179 | 2.39847084284171e-05 | 0.000109030923467171 | 1.84966509422357 |
| ***SPRR2G*** | 3.98891354799094 | 9.71932402463414 | 4.67155047134886 | 2.44269852708694e-05 | 0.000110799527960676 | 1.83186122524144 |
| ***KRTDAP*** | 3.31102530092002 | 12.0584722584023 | 4.66877405510094 | 2.46547387515339e-05 | 0.000111741313917156 | 1.82281890070152 |
| ***ALOX12B*** | 3.16082287173912 | 7.17169223051124 | 4.65400452379263 | 2.59017556231939e-05 | 0.000116662964213066 | 1.77475065297599 |
| ***TNNC1*** | -3.33948009959789 | 5.76684479738028 | -4.6383167671912 | 2.72940251986024e-05 | 0.000122371274214784 | 1.72375647359637 |
| ***C10orf99*** | 2.71899728730503 | 10.6937766859678 | 4.61414128819468 | 2.9584358181731e-05 | 0.00013119197813496 | 1.64529982017525 |
| ***KRT23*** | 2.58364765983831 | 7.00382572736071 | 4.56713550704513 | 3.45892005510125e-05 | 0.000151093030259026 | 1.49320267463263 |
| ***CXCL9*** | 3.63341023146077 | 8.34736972352256 | 4.56378686993494 | 3.49758053110769e-05 | 0.000152501835932277 | 1.48239051318119 |
| ***KRT13*** | -3.7429485516103 | 11.7468128517743 | -4.55572297145183 | 3.59242354957058e-05 | 0.000155699731047743 | 1.45636633828855 |
| ***GZMB*** | 2.13516732999583 | 7.32415578004553 | 4.4820705955566 | 4.58363304345847e-05 | 0.000192985318432367 | 1.2195196197644 |
| ***ACTC1*** | -3.17156794208385 | 6.71153887186792 | -4.40142648896815 | 5.97610501562598e-05 | 0.000243547847174618 | 0.96199933947827 |
| ***ASPRV1*** | 2.90140856744647 | 6.95635316941558 | 4.34102645826738 | 7.28172266133026e-05 | 0.000288841087513972 | 0.770416566173514 |
| ***GZMA*** | 2.13177516641803 | 6.78565040894562 | 4.33672340315499 | 7.3846856570541e-05 | 0.000292159915211147 | 0.756811133233198 |
| ***S100A12*** | 2.3231349699826 | 9.22022466473405 | 4.31308376093913 | 7.97615442858074e-05 | 0.000312516650582077 | 0.68217187846295 |
| ***LCE1B*** | 2.50615851674213 | 5.22712494633727 | 4.19000869121604 | 0.000118830015703645 | 0.000441497377104598 | 0.296522792654737 |
| ***MFAP4*** | -2.1148903714451 | 7.30890522169041 | -4.14086600492325 | 0.000139165605377949 | 0.000506432668379969 | 0.143967587684743 |
| ***TSPAN7*** | -2.06821934525068 | 7.07275062948946 | -4.13940288654493 | 0.000139820150647536 | 0.000508592362783439 | 0.13943849437606 |
| ***KLK5*** | 2.33372740993617 | 8.86611110503014 | 4.10281248273137 | 0.000157196512519421 | 0.000564768143339835 | 0.0264192136757213 |
| ***FST*** | 2.1113857759888 | 7.6482495235631 | 4.0558488373674 | 0.000182591014299896 | 0.000645149752689663 | -0.117935303721 |
| ***TMEM45A*** | 2.02422776241755 | 8.85973680112553 | 4.00105586999315 | 0.000217260620068781 | 0.00075092195224437 | -0.285328794843555 |
| ***PIP*** | -2.03605756415588 | 4.37072311752362 | -3.91146277470854 | 0.000288091640640129 | 0.000960640376637263 | -0.556577208033522 |

| | | | | | | |
|---|---|---|---|---|---|---|
| ***WFDC12*** | 2.90384137627965 | 7.13881145718728 | 3.87188400175828 | 0.00032606149926675 | 0.0010718137619473 | -0.675401812773321 |
| ***ALOX12*** | -2.33550233140769 | 6.6797485082754 | -3.8682765774352 | 0.000329753046751448 | 0.00108245304428304 | -0.686200913943157 |
| ***LCE2B*** | 2.56021870639626 | 5.8169656708233 | 3.86558727507799 | 0.000332531264965806 | 0.00108985453547234 | -0.694248127990666 |
| ***KRT1*** | 3.62401320289026 | 9.95384884638287 | 3.78160128021344 | 0.000431592563380824 | 0.00137004111600362 | -0.944068721385507 |
| ***HORMAD1*** | 2.26899769276991 | 5.32636511138706 | 3.73950896390491 | 0.000491382803672318 | 0.00153410908677903 | -1.06816484073323 |
| ***TNNI1*** | -2.48184928922148 | 5.59661967976072 | -3.73411632593801 | 0.000499595551762983 | 0.00155624902004194 | -1.08400870261625 |
| ***FMO2*** | -2.30742105518845 | 6.62685081804097 | -3.60515028133308 | 0.000740243182854266 | 0.00219581308167777 | -1.45911712738896 |
| ***CLCA4*** | -2.49060111911126 | 7.27518764725467 | -3.50208894189365 | 0.00100885313940266 | 0.0028787832602177 | -1.75344241278638 |
| ***MYH1*** | -2.36587789513063 | 4.76783029204238 | -3.49770673526983 | 0.00102212525122939 | 0.00290868241661997 | -1.76584638521056 |
| ***MYLPF*** | -2.13783543001888 | 6.03579835020563 | -3.4537290999134 | 0.00116487341543835 | 0.00324939400561735 | -1.88981468773183 |
| ***KRT75*** | 2.16388610566687 | 6.93676443222859 | 3.38821329770674 | 0.00141322887569512 | 0.00384871487141562 | -2.07274236320087 |
| ***SPRR3*** | -2.02857257035371 | 12.4053346114345 | -3.38116888040546 | 0.00144274790622022 | 0.00391333794878441 | -2.0922841107362 |
| ***LTF*** | -2.42906420647759 | 5.82982453138562 | -3.36090305686088 | 0.00153097270010031 | 0.00411155777053705 | -2.14836365301514 |
| ***GZMK*** | 2.06498603707607 | 6.39625992561945 | 3.34597518519489 | 0.00159921130006633 | 0.00426729628547017 | -2.18953893986401 |
| ***KLK6*** | 2.13331294680632 | 8.24685549391381 | 3.27733252505519 | 0.00195185402454463 | 0.0050741502470447 | -2.37740075834965 |
| ***CEACAM7*** | -2.17632238270575 | 5.18425623840581 | -3.27630595855976 | 0.00195764860029098 | 0.00508762600818326 | -2.3801916845384 |
| ***FLG*** | 2.46958161978005 | 6.52026751010572 | 3.10561794644631 | 0.00318345116090318 | 0.00776660631992066 | -2.83637452171024 |
| ***DSC1*** | 2.48208719162928 | 6.68797732724549 | 2.96961454412368 | 0.00464376956264177 | 0.0108556667839104 | -3.18820843382069 |
| ***FLG2*** | 2.03950511856142 | 4.8171382907359 | 2.80157607961931 | 0.00730997663149368 | 0.0160945103468815 | -3.60778041379389 |
| ***SPINK6*** | 2.03671231555239 | 5.91913010050111 | 2.6947453695707 | 0.00967938272656132 | 0.0205009916955246 | -3.86539363519263 |
| ***TGM3*** | -2.49257062246349 | 9.80247791531332 | -2.60366143976137 | 0.0122370118061711 | 0.0250854976792106 | -4.0791940653021 |

Table S3 - Differentially expressed genes in advanced OSCC (stages III and IV) in comparison to normal samples (healthy oral mucosa).

| Gene | log2 Fold Change | Average Expression | t | p-value | adjusted p-value | B |
|---|---|---|---|---|---|---|
| *GPD1* | -2.08136717503126 | 4.34764942208053 | -40.4054515583771 | 1.00166705284491e-38 | 1.66837664321848e-34 | 76.9767068201681 |
| *FGF10* | -2.07807112156927 | 3.98665990216155 | -39.1696710772738 | 4.26217011400672e-38 | 3.5495352709448e-34 | 75.6545766522419 |
| *PI16* | -2.48206274843036 | 5.4044534042165 | -26.1429448301638 | 4.79074602577852e-30 | 1.32991109675612e-26 | 58.134937225245 |
| *PEBP4* | -2.4878240125763 | 4.39126680783299 | -22.2127554039154 | 6.63485639647917e-27 | 6.39261633436821e-24 | 51.0836456921561 |
| *GPRC5B* | -2.50704751008409 | 5.27332415686965 | -21.1559052916665 | 5.56923517260463e-26 | 3.71044724139611e-23 | 48.9941857937809 |
| *ADIPOQ* | -2.84590894995462 | 4.32248603708525 | -19.9016330416279 | 7.80054493393114e-25 | 4.19115730385668e-22 | 46.3936678131834 |
| *PRKAA2* | -3.03551700832938 | 4.26300361290586 | -19.6294460172783 | 1.40745468534338e-24 | 6.51182367752206e-22 | 45.8110712112236 |
| *HSPB6* | -3.62422649780632 | 4.87322560445144 | -19.4550853734636 | 2.06106564179694e-24 | 9.03397613941312e-22 | 45.4343188589318 |
| *MYH11* | -2.16281076208699 | 5.12377460935976 | -18.7805374956597 | 9.24769927453584e-24 | 3.75682144186998e-21 | 43.95008011015 |
| *EPB41L4A* | -2.01898797478866 | 5.37963954369296 | -18.754081060835 | 9.81677352162674e-24 | 3.89305189943369e-21 | 43.8909861491536 |
| *PYGM* | -4.13831457984942 | 4.70821972413499 | -18.4354910264137 | 2.02532265412431e-23 | 7.33342915806402e-21 | 43.1740415209571 |
| *ANK2* | -2.3522400028299 | 4.50064250052564 | -17.8980443917124 | 7.02309509600412e-23 | 2.29366023370676e-20 | 41.9418642827752 |
| *SORBS1* | -2.24768297988752 | 5.33599733786061 | -17.3228314215899 | 2.74196320230296e-22 | 7.61168984959303e-20 | 40.5905475745259 |
| *NPY6R* | -2.86801699858592 | 4.18982602637732 | -16.7459923323941 | 1.11135277627362e-21 | 2.87761199341179e-19 | 39.2004365572321 |
| *CASQ1* | -3.04371725653104 | 4.29836716287083 | -16.7182151764612 | 1.18986398705478e-21 | 3.0027840255128e-19 | 39.132592303555 |
| *ADH1B* | -3.95364042663156 | 4.17835191661223 | -16.4241894581426 | 2.46301331681914e-21 | 5.26919460052017e-19 | 38.4092628042487 |
| *RBP4* | -2.03122133393179 | 4.10461929499294 | -16.1411395851625 | 5.00530609800486e-21 | 1.00443829359481e-18 | 37.7038760861349 |
| *ABCA8* | -3.39874492280139 | 4.725423402863 | -16.088128481201 | 5.72171656562189e-21 | 1.13453465615474e-18 | 37.5707676307454 |
| *ADH1C* | -2.36865313141433 | 4.30073332316218 | -16.0764736511206 | 5.89273460243936e-21 | 1.15469867692035e-18 | 37.5414603121387 |
| *ACACB* | -2.38444983048822 | 5.18372263587103 | -15.9986607226115 | 7.17591852680124e-21 | 1.35820567025456e-18 | 37.3453970698404 |
| *AMPD1* | -3.25454259723796 | 4.38398875798019 | -15.9880741671748 | 7.37123717228938e-21 | 1.3794980487826e-18 | 37.3186692949685 |
| *SMYD1* | -2.89558207875655 | 4.18494189107435 | -15.8739969970683 | 9.85269731245633e-21 | 1.80336842237662e-18 | 37.0298497965761 |
| *MAOB* | -2.63954480452642 | 5.24177456169648 | -15.7461665020065 | 1.36616799292495e-20 | 2.39525200949031e-18 | 36.7044404201632 |
| *LDB3* | -2.67900864462037 | 4.59365084194251 | -15.559514939015 | 2.20897242238996e-20 | 3.71642875427547e-18 | 36.2259096007241 |

| | | | | | | |
|---|---|---|---|---|---|---|
| ***ATP2A1*** | -3.11862174093512 | 4.4482577826067 | -15.5070259987797 | 2.53035447820798e-20 | 4.13192001853256e-18 | 36.0906117390625 |
| ***NRAP*** | -3.77964032368 | 4.59777507517331 | -15.3829190487275 | 3.49300859776613e-20 | 5.24140100940474e-18 | 35.7694278584912 |
| ***SH3BGRL2*** | -3.7505519338858 | 6.58993461191685 | -15.3080495966226 | 4.24653686542452e-20 | 6.31520696700989e-18 | 35.5747946877122 |
| ***PADI2*** | -2.43862480568581 | 5.24952075527924 | -15.2688437103985 | 4.70511361200278e-20 | 6.81464107143638e-18 | 35.4726102694666 |
| ***GSTM5*** | -2.11579815440819 | 4.74658663363261 | -14.9180448015049 | 1.18704802212234e-19 | 1.60743673629835e-17 | 34.5501810706534 |
| ***TENM2*** | 2.89733246488233 | 8.09174558368225 | 14.8621485939911 | 1.37745859057469e-19 | 1.82214101217568e-17 | 34.4018417134883 |
| ***CAB39L*** | -2.01180570598866 | 5.01798758255627 | -14.8059130594856 | 1.60044584962373e-19 | 2.06643612956068e-17 | 34.252221820776 |
| ***TMOD1*** | -2.46792622907823 | 5.57232474043829 | -14.4184026307205 | 4.54636599764764e-19 | 5.40887657548707e-17 | 33.2107696407969 |
| ***KRT17*** | 2.62780674921354 | 11.6662867931092 | 14.2873039150811 | 6.4985767867577e-19 | 7.54420248663076e-17 | 32.8542699323831 |
| ***GATM*** | -2.79604643074336 | 5.5625312855606 | -14.2858764307379 | 6.52397960118514e-19 | 7.54420248663076e-17 | 32.8503764612793 |
| ***ANO5*** | -2.31535043429228 | 3.82590451937045 | -13.8593731143372 | 2.11613073799654e-18 | 2.13613779224669e-16 | 31.6757291675747 |
| ***CAPN6*** | -2.3675436322562 | 4.55210977407805 | -13.3767244428707 | 8.2324470017713e-18 | 7.1416477740366e-16 | 30.3188216081532 |
| ***IGF2BP2*** | 2.07145423346012 | 6.61646521498684 | 13.1403590454111 | 1.61828733263385e-17 | 1.30213496678016e-15 | 29.6434650585908 |
| ***C10orf71*** | -2.02082708616181 | 4.43370166483694 | -13.1232153931989 | 1.70005813973845e-17 | 1.3548405921284e-15 | 29.5942018409424 |
| ***PTGIS*** | -2.34487443933367 | 4.84570441005922 | -12.9111491302201 | 3.13772101275102e-17 | 2.39733399946701e-15 | 28.9816799316544 |
| ***TCEA3*** | -2.53832682124683 | 6.63687290162367 | -12.8698681787797 | 3.53761776719349e-17 | 2.67829825138067e-15 | 28.8617693817733 |
| ***CTTNBP2*** | -2.06314868394882 | 4.88373757448073 | -12.7551749041469 | 4.94250898895785e-17 | 3.54199851181629e-15 | 28.5274542804274 |
| ***MYL3*** | -2.13601646418903 | 4.46130239624743 | -12.6176457490686 | 7.39708797089602e-17 | 5.07020153264379e-15 | 28.1243208076122 |
| ***CDH3*** | 2.68016824519016 | 10.3515659739691 | 12.6036941042349 | 7.70697139983283e-17 | 5.23948227084145e-15 | 28.0832873247582 |
| ***INHBA*** | 3.26741693688398 | 7.49495296213354 | 12.5918216001935 | 7.98103396596149e-17 | 5.403743973051e-15 | 28.0483488660363 |
| ***LAMC2*** | 3.98701945102124 | 8.54873235040382 | 12.5770985058905 | 8.3346825530835e-17 | 5.51300612519692e-15 | 28.004996152477 |
| ***ANGPTL1*** | -3.58423756051139 | 4.53545990954199 | -12.5523941529256 | 8.96419763016536e-17 | 5.90148915921084e-15 | 27.9321897314869 |
| ***BARX2*** | -2.2001087823036 | 5.91988461468087 | -12.4550974660402 | 1.19504527729451e-16 | 7.56831716297239e-15 | 27.6446712890189 |
| ***PPARG*** | -2.18972872556272 | 4.48386359234611 | -12.3385320432174 | 1.68920194037793e-16 | 1.03820470549575e-14 | 27.2985841609745 |
| ***PRELP*** | -2.03230176827988 | 5.28144078803052 | -12.2861419363654 | 1.97460077664374e-16 | 1.19162864260066e-14 | 27.1424572349274 |
| ***TTN*** | -4.15571125352112 | 5.2155187712561 | -12.2675074036329 | 2.08752221213753e-16 | 1.25522635254017e-14 | 27.0868381753915 |
| ***CMYA5*** | -4.08514370505666 | 4.90452941250617 | -12.1993934341047 | 2.55904092926813e-16 | 1.49555739361017e-14 | 26.8831496333662 |

| | | | | | | |
|---|---|---|---|---|---|---|
| ***SYNPO2*** | -3.28205366308829 | 5.84008862291203 | -12.1290367733001 | 3.16015195570834e-16 | 1.81501693014752e-14 | 26.6721167362925 |
| ***MMP12*** | 5.03250841723111 | 8.30543557759231 | 12.1128196372318 | 3.31793631363845e-16 | 1.89909097044543e-14 | 26.6233819380076 |
| ***PODN*** | -2.17889758449074 | 5.53254694538584 | -12.0408179773076 | 4.12091360308615e-16 | 2.3267097278984e-14 | 26.4065906899933 |
| ***NDNF*** | -2.58303208607076 | 4.61978848335705 | -12.0236634642721 | 4.33972189707922e-16 | 2.42558415831381e-14 | 26.3548396400198 |
| ***SLC7A2*** | -2.05418220046858 | 4.91542722181237 | -11.802092443067 | 8.49475769927987e-16 | 4.53489372561556e-14 | 25.6829483332323 |
| ***PLN*** | -2.59955001258343 | 4.37505645530077 | -11.744519220922 | 1.01249898919785e-15 | 5.28657779438223e-14 | 25.507311076636 |
| ***DEPTOR*** | -2.80348237711016 | 5.65434555352311 | -11.6969933836892 | 1.17077305858871e-15 | 5.9634238727381e-14 | 25.3619982614801 |
| ***GPC3*** | -2.77988644842728 | 4.93965880028018 | -11.6511775282249 | 1.34710451946769e-15 | 6.75824484224512e-14 | 25.2216340700614 |
| ***DDX60L*** | 2.50047183406251 | 6.16269363947514 | 11.4909856982522 | 2.2047507601453e-15 | 1.05523932933851e-13 | 24.7287042897231 |
| ***MYOM1*** | -2.44877754789786 | 4.44913809666507 | -11.4897383078777 | 2.21325375781055e-15 | 1.05596319896529e-13 | 24.7248527707709 |
| ***STAC3*** | -2.62911678768718 | 5.63630149986437 | -11.4343555983258 | 2.62628052769899e-15 | 1.23568724489701e-13 | 24.5536452161094 |
| ***DES*** | -3.37948385752378 | 5.57057805060264 | -11.3685528666496 | 3.21999773475451e-15 | 1.47559479928716e-13 | 24.3497061230709 |
| ***CD80*** | 2.05377356283054 | 4.90239784139467 | 11.3597867367321 | 3.30876270937878e-15 | 1.50575824282549e-13 | 24.3224951461217 |
| ***ABCA6*** | -2.11772113984838 | 4.84121292749184 | -11.3558029855087 | 3.34991743012183e-15 | 1.52033309853158e-13 | 24.3101258644624 |
| ***AURKA*** | 2.03375853746558 | 6.589448343898 | 11.295293960508 | 4.04282503629918e-15 | 1.79566116812264e-13 | 24.1219956439913 |
| ***PPP1R3C*** | -3.4974844328399 | 6.24931355678531 | -11.2174420793201 | 5.15275546903838e-15 | 2.22342733399749e-13 | 23.87924474489 |
| ***MYO1B*** | 2.22640005653445 | 8.47014060797121 | 11.1758817928478 | 5.86705942838316e-15 | 2.48655831651781e-13 | 23.7493333370566 |
| ***ACTN2*** | -3.96969607706863 | 5.09151927976117 | -11.0724092070338 | 8.11354815926567e-15 | 3.34503114209725e-13 | 23.4249222920941 |
| ***ARHGEF26*** | -2.25164157329308 | 4.51141011070394 | -11.0413363193866 | 8.94554400431161e-15 | 3.66987637772941e-13 | 23.3272315161899 |
| ***FRZB*** | -2.39330849077291 | 5.08181173093489 | -10.9977331588372 | 1.02609989419604e-14 | 4.16846825310468e-13 | 23.189936750543 |
| ***MYPN*** | -2.20609877243341 | 4.33132487379578 | -10.9204135588352 | 1.30949912401097e-14 | 5.18076423029138e-13 | 22.9458767308604 |
| ***ITGA3*** | 2.30660654259562 | 7.92020826967946 | 10.9042943722435 | 1.37793183710291e-14 | 5.42572876567047e-13 | 22.8948997164123 |
| ***AQP1*** | -2.14298908903598 | 7.59308129544136 | -10.8962584705353 | 1.41338950341955e-14 | 5.52615388942628e-13 | 22.8694736956926 |
| ***COL14A1*** | -2.05141036502993 | 5.62167332821038 | -10.8276285357874 | 1.75648156264523e-14 | 6.72550733503885e-13 | 22.6519882500957 |
| ***ABI3BP*** | -3.37567622501558 | 6.05831443020489 | -10.8228932550811 | 1.78305727120584e-14 | 6.8116059424781e-13 | 22.6369601385489 |
| ***MMP1*** | 5.45392524375112 | 9.7704648852227 | 10.7834004807786 | 2.02118117229522e-14 | 7.63374004665516e-13 | 22.5115125767338 |
| ***CDK1*** | 2.10815757333638 | 6.53179469537793 | 10.7819070122518 | 2.03079295924188e-14 | 7.64891642147128e-13 | 22.5067647190752 |

| | | | | | | |
|---|---|---|---|---|---|---|
| ***PDK4*** | -3.33841206959673 | 5.90583464302015 | -10.7810288421 | 2.03646635485872e-14 | 7.64891642147128e-13 | 22.5039728122333 |
| ***MYH7*** | -4.0054164225434 | 5.07423292511225 | -10.7208594973181 | 2.46599810660477e-14 | 9.04689345057411e-13 | 22.3124467889864 |
| ***CKMT2*** | -3.14575537933255 | 4.83515373184243 | -10.7137289492461 | 2.52264309871306e-14 | 9.21428584477295e-13 | 22.2897189300597 |
| ***BNC1*** | 2.23891673434777 | 7.37178250300109 | 10.6827044233839 | 2.78485080500118e-14 | 1.00920740169957e-12 | 22.1907564873682 |
| ***FABP3*** | -2.13455659974931 | 4.63460743731224 | -10.6626565745151 | 2.96880442728437e-14 | 1.06340659227631e-12 | 22.1267427190688 |
| ***MELK*** | 2.36027769405767 | 7.80291326289654 | 10.6300358236122 | 3.29480929257969e-14 | 1.16514529887914e-12 | 22.0224743730094 |
| ***KYNU*** | 2.95327633547691 | 6.84414749578844 | 10.6222522049891 | 3.37781355349769e-14 | 1.19196742684444e-12 | 21.9975750928012 |
| ***CD36*** | -2.42033090654523 | 5.6453203645514 | -10.5491264075441 | 4.26886613226625e-14 | 1.48335415965015e-12 | 21.7632775721059 |
| ***FERMT1*** | 2.08416658875202 | 7.84072341498534 | 10.4807348363462 | 5.31705450834573e-14 | 1.81106052946844e-12 | 21.5435405713434 |
| ***LIFR*** | -2.35727622804866 | 5.53535870825693 | -10.4047877484252 | 6.78988821541675e-14 | 2.26184756231963e-12 | 21.2988429506361 |
| ***CHRDL1*** | -3.69478381762155 | 5.52064245561558 | -10.3871400877814 | 7.18758373747989e-14 | 2.37062167785079e-12 | 21.2418802401339 |
| ***GULP1*** | -2.37753844411176 | 4.99119458991965 | -10.2363971414404 | 1.17064158207908e-13 | 3.69284208164947e-12 | 20.7537451087738 |
| ***TBX15*** | -2.20855583240531 | 5.45683035472589 | -10.2338671781982 | 1.18029281065698e-13 | 3.7162489705676e-12 | 20.7455287147176 |
| ***NEB*** | -4.32498531521115 | 5.44604642890316 | -10.2223954334647 | 1.22507513775676e-13 | 3.84997198008991e-12 | 20.70826284549 |
| ***PRUNE2*** | -2.41821270798965 | 4.83737999859061 | -10.2180172113897 | 1.2426162793573e-13 | 3.8904166821382e-12 | 20.6940359763656 |
| ***ETNK2*** | -2.42607065275693 | 5.48562231017895 | -10.1515117396788 | 1.54259236908898e-13 | 4.78462169451511e-12 | 20.477640919454 |
| ***OCIAD2*** | 2.10549875815185 | 9.01416044729801 | 10.0896506649226 | 1.8872014308981e-13 | 5.73599033449611e-12 | 20.2758746664455 |
| ***LYVE1*** | -2.82378013987701 | 5.60923414561174 | -10.0857690867819 | 1.91125912659477e-13 | 5.78798763864772e-12 | 20.2631990428373 |
| ***EMCN*** | -2.2270202490142 | 5.28308334530805 | -10.0378415191661 | 2.23517957448922e-13 | 6.68386911897531e-12 | 20.1065376662043 |
| ***TRDN*** | -3.74531637714174 | 4.27122912065231 | -9.98051369112048 | 2.69649802011589e-13 | 7.81416651542336e-12 | 19.918786867914 |
| ***RBP1*** | 2.15466316063539 | 7.64571247957669 | 9.96270159321966 | 2.85860383063989e-13 | 8.25180336276222e-12 | 19.8603714897518 |
| ***PLP1*** | -2.09735302384916 | 4.71513386953937 | -9.92319791724108 | 3.25419104251396e-13 | 9.3451389662263e-12 | 19.7306829502039 |
| ***MMP11*** | 2.27261142031335 | 5.44587423673276 | 9.87983385333972 | 3.7525506941349e-13 | 1.07208377978578e-11 | 19.5881077018594 |
| ***LMOD3*** | -2.71923106180474 | 5.05447855637064 | -9.82575298600254 | 4.48374164037546e-13 | 1.26364129885099e-11 | 19.4099860214282 |
| ***LRRC39*** | -2.51728939495222 | 4.17196297015904 | -9.73387000290511 | 6.07215049062238e-13 | 1.67446587039414e-11 | 19.1065734576313 |
| ***PLA2G7*** | 3.00171506595549 | 7.01600668773773 | 9.6922019040431 | 6.96964651011008e-13 | 1.89994160838615e-11 | 18.9686555357833 |
| ***SLC16A1*** | 2.07052530370767 | 6.88618034934523 | 9.68196916860782 | 7.20985543481086e-13 | 1.95582006713697e-11 | 18.934755409475 |

| | | | | | | |
|---|---|---|---|---|---|---|
| ***IGSF10*** | -2.48601671080636 | 4.61081526549187 | -9.66947000912178 | 7.51465335272063e-13 | 2.02530851525752e-11 | 18.8933304902094 |
| ***SPARCL1*** | -2.22801496716039 | 9.72422364589674 | -9.63570974993195 | 8.40463864091099e-13 | 2.23622461985644e-11 | 18.7813521467712 |
| ***C12orf75*** | 2.07920156196493 | 8.55302002530123 | 9.59360497244621 | 9.66551046757315e-13 | 2.54729022702371e-11 | 18.6415138019086 |
| ***ENO3*** | -2.42221076891232 | 4.90410063925229 | -9.51751572528825 | 1.2449570171063e-12 | 3.23494603384128e-11 | 18.3882969512314 |
| ***SLC28A3*** | 2.29805877897348 | 5.99601594336461 | 9.42935811104224 | 1.67062784871447e-12 | 4.21605718911943e-11 | 18.0941069989916 |
| ***PBX1*** | -2.31003478049692 | 6.51473297380601 | -9.40778028712275 | 1.79556310564477e-12 | 4.49727805828862e-11 | 18.0219686425765 |
| ***XDH*** | 2.07817147054642 | 6.02651755582369 | 9.37907467626524 | 1.97652901608391e-12 | 4.92093681493179e-11 | 17.9259215547909 |
| ***KRT36*** | -2.16120228759209 | 3.92189605447652 | -9.34333626584684 | 2.22781363056018e-12 | 5.51359046517242e-11 | 17.8062172891876 |
| ***MYBPC1*** | -5.09855702192972 | 5.51114037326454 | -9.29675904976116 | 2.60441796776966e-12 | 6.30511419639119e-11 | 17.6500008107533 |
| ***CDC6*** | 2.11619779019291 | 5.60455612334508 | 9.22469627101474 | 3.31790015124816e-12 | 7.77256609271299e-11 | 17.4078483344165 |
| ***TPM2*** | -2.10964540234607 | 6.94917881671078 | -9.09869705397481 | 5.07352742823157e-12 | 1.14380843111225e-10 | 16.983135266828 |
| ***CD274*** | 2.14319860653362 | 5.47190764626197 | 9.06603946575486 | 5.66549042010329e-12 | 1.26155626252995e-10 | 16.8727853844727 |
| ***PPARGC1A*** | -2.09626038161743 | 5.06154304890019 | -9.05344232312938 | 5.91204781438662e-12 | 1.31294757861898e-10 | 16.830190478038 |
| ***CRYAB*** | -3.39677448616974 | 9.55597766534151 | -9.03244117709015 | 6.34741480610779e-12 | 1.39844630966311e-10 | 16.7591430380213 |
| ***PDGFD*** | -2.11316471250018 | 5.07167879564174 | -9.0277378883975 | 6.44927109445966e-12 | 1.41901003103461e-10 | 16.7432255459624 |
| ***SPP1*** | 3.28544393994578 | 6.57258970629775 | 8.9480551571962 | 8.4488162830139e-12 | 1.82048491603984e-10 | 16.4732142091293 |
| ***ACPP*** | -2.21146086948667 | 6.01428802783524 | -8.91776274094668 | 9.36393819199669e-12 | 2.01246134872125e-10 | 16.3703998851759 |
| ***PARP12*** | 2.01466831922871 | 8.5599951114481 | 8.91397328091075 | 9.48524493178361e-12 | 2.03328493672829e-10 | 16.3575318530026 |
| ***GGTA1P*** | -2.05179442649466 | 6.16636083805701 | -8.91197014188566 | 9.55000796536968e-12 | 2.04453640965549e-10 | 16.35072913723 |
| ***CEP55*** | 2.12909702144496 | 7.86955438695579 | 8.89688016243489 | 1.00524372879899e-11 | 2.141092013667e-10 | 16.2994704890842 |
| ***TXLNB*** | -2.49650874781782 | 4.90135623703117 | -8.87272661230352 | 1.09127645692844e-11 | 2.31250644613233e-10 | 16.2173777598026 |
| ***PLAU*** | 2.34233637040769 | 8.57020412917519 | 8.86478621279782 | 1.12115240618366e-11 | 2.36678257001204e-10 | 16.1903776440198 |
| ***IL24*** | 3.08090068168274 | 6.37862107758704 | 8.82494689911322 | 1.28398134735288e-11 | 2.66658270841765e-10 | 16.0548181358188 |
| ***TPX2*** | 2.06504506411557 | 7.73276443015548 | 8.80309345998864 | 1.38322849677591e-11 | 2.85844340475181e-10 | 15.9803936631762 |
| ***CLU*** | -2.28140796528019 | 6.53915125236276 | -8.78445989837517 | 1.47394498837065e-11 | 3.02340242934748e-10 | 15.9168989384558 |
| ***TGFBR3*** | -2.66199861497244 | 7.2890021960777 | -8.77606265593817 | 1.51676558399329e-11 | 3.09978497754506e-10 | 15.8882741903981 |
| ***ZNF667-AS1*** | -2.08721563760733 | 4.87188084099555 | -8.7650430237676 | 1.57486852787526e-11 | 3.19500733255667e-10 | 15.8507000703622 |

| | | | | | | |
|---|---|---|---|---|---|---|
| ***MMP13*** | 4.40668428420738 | 6.36886909977083 | 8.76192010932157 | 1.59173924609616e-11 | 3.22138625552583e-10 | 15.8400496532733 |
| ***HLF*** | -2.52348332468166 | 5.72749924403696 | -8.71232620226999 | 1.88546076998038e-11 | 3.74752202682496e-10 | 15.6707918467966 |
| ***FCER1A*** | -2.06101493935298 | 6.56111560497719 | -8.6203059037696 | 2.58318110153398e-11 | 5.0204742622112e-10 | 15.3561381949974 |
| ***APOBEC2*** | -2.43937265748253 | 4.99538039473499 | -8.5377168093501 | 3.42906839702134e-11 | 6.45362296280083e-10 | 15.0730866889177 |
| ***CCL20*** | 3.08959780918086 | 7.0215256134595 | 8.52265390404763 | 3.61112955002924e-11 | 6.75050210833748e-10 | 15.0213982407828 |
| ***PDLIM3*** | -2.90879425569286 | 5.89916981439912 | -8.48922517785094 | 4.05075312185539e-11 | 7.47168815034588e-10 | 14.9066175167645 |
| ***APOL1*** | 2.42139679028676 | 7.33424397000342 | 8.47903741548842 | 4.19517368756771e-11 | 7.69550823130458e-10 | 14.8716178621491 |
| ***KLHL41*** | -4.67764097410293 | 5.34685620455355 | -8.4716620614424 | 4.30295388199259e-11 | 7.8499452199856e-10 | 14.8462746245051 |
| ***ANLN*** | 2.29705653051146 | 8.0352209481484 | 8.46893601257355 | 4.34349374014256e-11 | 7.91523323148955e-10 | 14.8369061922628 |
| ***PTHLH*** | 3.15433826304522 | 8.34883023485729 | 8.4166881668643 | 5.19935818593778e-11 | 9.31188278978277e-10 | 14.6572288874351 |
| ***LIMCH1*** | -2.01701027225028 | 6.3301296461594 | -8.38821582127987 | 5.73535725475612e-11 | 1.01733876927815e-09 | 14.5592187973534 |
| ***CILP*** | -2.92319397967452 | 4.9482775836485 | -8.38001794337045 | 5.89976765015188e-11 | 1.04427768311296e-09 | 14.5309870073713 |
| ***FAM83A*** | 2.07137838049149 | 7.5224298204213 | 8.35315713291045 | 6.47247997732592e-11 | 1.13839098735312e-09 | 14.4384457504304 |
| ***IL12RB2*** | 2.30734176006032 | 5.71180489834863 | 8.27069765923398 | 8.60516066653581e-11 | 1.46701695047923e-09 | 14.1539960710529 |
| ***SERPINE1*** | 2.91561363307366 | 8.26988271662929 | 8.24204924020407 | 9.50145134873764e-11 | 1.60340601483864e-09 | 14.0550473450905 |
| ***OGN*** | -3.24395692236483 | 4.73044310481258 | -8.15711022239619 | 1.27510082052183e-10 | 2.10278012540709e-09 | 13.7613130610089 |
| ***MGST1*** | -3.05146119197229 | 5.70340938323627 | -8.13504103288501 | 1.37651124788798e-10 | 2.24996774728383e-09 | 13.6849070264092 |
| ***DDX60*** | 2.54157803905419 | 8.06487254992665 | 8.10987853186526 | 1.50208285801374e-10 | 2.412083400228e-09 | 13.5977489721631 |
| ***RRAGD*** | -2.61393719252033 | 7.55610374551101 | -8.10741286617059 | 1.51499168404475e-10 | 2.42631745090859e-09 | 13.5892059617764 |
| ***FAM3B*** | -3.44394922217547 | 5.58659857047138 | -8.09035496590153 | 1.60740871844745e-10 | 2.55711553146713e-09 | 13.5300922192378 |
| ***MYL2*** | -5.20064688243788 | 5.79577615043568 | -8.05461264434963 | 1.81987828493135e-10 | 2.86772873356826e-09 | 13.4061623679844 |
| ***TNC*** | 2.19461418751348 | 8.59328585705563 | 7.99559205217759 | 2.23442231718382e-10 | 3.48469458005746e-09 | 13.2013292521813 |
| ***IFI30*** | 2.03636700987216 | 9.7267663579285 | 7.98730105434562 | 2.29982074046475e-10 | 3.57998264048419e-09 | 13.1725364853449 |
| ***MFAP4*** | -3.31478212559556 | 7.30890522169041 | -7.98240370735769 | 2.33935170820707e-10 | 3.61784977269238e-09 | 13.1555269900222 |
| ***XIRP2*** | -4.17422243489699 | 4.78168660621679 | -7.9539054655337 | 2.58333702606879e-10 | 3.96206827865578e-09 | 13.0565158859756 |
| ***APOD*** | -3.51001362645984 | 7.99542625476502 | -7.93885475064711 | 2.72236331428637e-10 | 4.12590385466367e-09 | 13.0042043682964 |
| ***PLEK2*** | 2.06316612156039 | 8.98623050470374 | 7.93285968060263 | 2.77981517229392e-10 | 4.19389506428691e-09 | 12.9833634094776 |

| | | | | | | |
|---|---|---|---|---|---|---|
| ***FHL1*** | -3.09855083613895 | 7.93114278062943 | -7.88167106369046 | 3.32279389340088e-10 | 4.94146920432903e-09 | 12.8053228513971 |
| ***PTPRZ1*** | 2.14469416762315 | 7.86476800010851 | 7.87385103423654 | 3.41466500878935e-10 | 5.05239396009313e-09 | 12.7781096712624 |
| ***MLIP*** | -2.3013176384848 | 4.42065695961331 | -7.83870419837157 | 3.86016816653589e-10 | 5.66475427152615e-09 | 12.6557559518458 |
| ***IFIH1*** | 2.00970837316567 | 6.78144741197759 | 7.79241017637619 | 4.53744074827113e-10 | 6.5095273990701e-09 | 12.4944866325524 |
| ***DCT*** | -2.50950294502393 | 4.40464388651133 | -7.76282813715812 | 5.03157730435363e-10 | 7.13849672753953e-09 | 12.3913716216695 |
| ***MYOZ2*** | -2.871794800805 | 4.27777708646794 | -7.76166181463058 | 5.05213147917624e-10 | 7.16155760997102e-09 | 12.3873051463493 |
| ***GBP5*** | 3.84404787841928 | 6.65113499294338 | 7.73719374086172 | 5.50333878959825e-10 | 7.74840328652143e-09 | 12.3019782638293 |
| ***AIM2*** | 2.98210226505534 | 6.73598107506574 | 7.71222823533348 | 6.00551241845474e-10 | 8.33565123681518e-09 | 12.2148837451667 |
| ***MGP*** | -2.81556874480776 | 7.56543062693426 | -7.63857058032807 | 7.77206334993229e-10 | 1.04144398355971e-08 | 11.9577357609352 |
| ***OAS3*** | 2.10456743050666 | 7.54502085561267 | 7.62842090657427 | 8.05338601302066e-10 | 1.07654251551262e-08 | 11.9222809843082 |
| ***ANKRD1*** | -2.67504029023864 | 4.64863450754387 | -7.58210820307846 | 9.47269368253855e-10 | 1.245281657272e-08 | 11.7604401712574 |
| ***MMP3*** | 3.43431877422612 | 8.09973724881271 | 7.56711511356906 | 9.984044668756e-10 | 1.30529237050863e-08 | 11.7080254249961 |
| ***OLR1*** | 2.35646538581677 | 5.36535198365359 | 7.55070733778484 | 1.0575481186266e-09 | 1.3750602235632e-08 | 11.6506536060012 |
| ***MMP9*** | 2.6462734262085 | 8.10005790349088 | 7.53745799585122 | 1.10786597749029e-09 | 1.43154505206193e-08 | 11.604317084413 |
| ***TGFBI*** | 2.01916938907555 | 11.2466313433113 | 7.51682740408387 | 1.19104655370183e-09 | 1.52718024622461e-08 | 11.5321514462639 |
| ***STAT1*** | 2.19015112445615 | 9.62514058348407 | 7.48653253431822 | 1.32469662290316e-09 | 1.68171851761243e-08 | 11.4261482955761 |
| ***IGFBP5*** | -2.17798254400263 | 8.00863380714789 | -7.37061215856151 | 1.99081970742003e-09 | 2.43637715259281e-08 | 11.0202136542121 |
| ***OAS2*** | 2.07845177581204 | 7.27745008859287 | 7.34179688232143 | 2.20318756497362e-09 | 2.67270881880558e-08 | 10.919234622869 |
| ***RYR1*** | -2.33203217583893 | 6.15579506836231 | -7.2931767555592 | 2.61433403822422e-09 | 3.1169898167976e-08 | 10.7487941601732 |
| ***CA3*** | -2.65678119412827 | 4.90501225117326 | -7.26853011520533 | 2.85132336290978e-09 | 3.37059204631833e-08 | 10.6623680501219 |
| ***PAX9*** | -2.53039555413222 | 6.18159246329995 | -7.25114742453632 | 3.03131706603962e-09 | 3.5656509217483e-08 | 10.6014039430207 |
| ***IL1A*** | 2.67220741733612 | 7.81560984183177 | 7.21238832446207 | 3.47477805087136e-09 | 4.05293439883147e-08 | 10.4654416766492 |
| ***CXCL10*** | 5.06417950953938 | 8.7910234744076 | 7.19343163463332 | 3.71481083068964e-09 | 4.29978382181839e-08 | 10.3989309973192 |
| ***CXCL1*** | 2.01526955248312 | 7.78239209693998 | 7.1852340531995 | 3.82369515369537e-09 | 4.41048936841759e-08 | 10.3701668295881 |
| ***TNNT3*** | -2.94225203432099 | 6.20846477089669 | -7.17679450998668 | 3.93913804890265e-09 | 4.53422828904786e-08 | 10.3405521526291 |
| ***SMPX*** | -3.86361873084411 | 4.98527522977155 | -7.14227860553897 | 4.44880741282679e-09 | 5.07182315318569e-08 | 10.2194195411147 |
| ***CASQ2*** | -3.37306927749396 | 5.20907556393222 | -7.12606814972673 | 4.71049914323964e-09 | 5.35174668966701e-08 | 10.1625216186852 |

| | | | | | | |
|---|---|---|---|---|---|---|
| ***XAF1*** | 2.32938543423543 | 8.34100141913849 | 7.10373749703009 | 5.09645268796449e-09 | 5.74013259286714e-08 | 10.0841346639549 |
| ***IFI44*** | 2.76816474609184 | 7.56919621725605 | 7.10172274306308 | 5.13279672167275e-09 | 5.7725767856976e-08 | 10.0770619021143 |
| ***SGCG*** | -2.11894935218981 | 4.34797904382266 | -7.08845946552637 | 5.37862390512701e-09 | 6.02058869380346e-08 | 10.0304998056398 |
| ***ATP1A2*** | -2.2851914105853 | 4.83917220781525 | -7.07010025136393 | 5.73849529937452e-09 | 6.37202518042546e-08 | 9.96604362289639 |
| ***MYOT*** | -3.78283338287261 | 5.32366361309682 | -7.02066227102957 | 6.83216189868994e-09 | 7.46696119321388e-08 | 9.79245343857368 |
| ***WIF1*** | -2.01269175473823 | 4.4327768748711 | -7.00891054545645 | 7.12148856792056e-09 | 7.72739502197296e-08 | 9.7511859771799 |
| ***CXCL13*** | 2.939431279451 | 7.29997954906581 | 6.98447045308232 | 7.76311872772587e-09 | 8.34209713090336e-08 | 9.6653579598787 |
| ***SLN*** | -4.3360283376276 | 6.07360913404399 | -6.90777965485445 | 1.01773582935249e-08 | 1.05748022293793e-07 | 9.39601304492939 |
| ***CXCL11*** | 4.54515695648291 | 6.54242117905535 | 6.86853370659728 | 1.16905054912209e-08 | 1.20195715717145e-07 | 9.25817128005641 |
| ***MYOZ1*** | -3.27779664371871 | 5.0970411631291 | -6.79327089399796 | 1.52511518525221e-08 | 1.52292077491372e-07 | 8.99383738960363 |
| ***MANSC1*** | -2.03646641302041 | 7.43931704291647 | -6.75939638839449 | 1.71902350235009e-08 | 1.70024082275197e-07 | 8.87487579231522 |
| ***KRT4*** | -4.24007356457859 | 8.90143884146788 | -6.75335849942506 | 1.75609148430133e-08 | 1.72767039353355e-07 | 8.85367278623111 |
| ***LRRN1*** | -2.21513437737785 | 4.49099385170702 | -6.73915323257781 | 1.8464857152033e-08 | 1.80381619193116e-07 | 8.80379019366621 |
| ***GPX3*** | -2.28411645851039 | 9.51194152700226 | -6.72674821049399 | 1.92922373356303e-08 | 1.87693636134497e-07 | 8.76023107868212 |
| ***DDX58*** | 2.16451576114917 | 7.34897387285247 | 6.72262839738014 | 1.95751410768232e-08 | 1.90334821818778e-07 | 8.74576512953105 |
| ***CRISP3*** | -5.26445972046814 | 5.6163693943407 | -6.71941942627189 | 1.97983702384588e-08 | 1.92393030741989e-07 | 8.73449757612036 |
| ***FLNC*** | -2.10074460375628 | 6.41455647898699 | -6.5951227450105 | 3.07172075159201e-08 | 2.82510109544542e-07 | 8.2981850246673 |
| ***ASB5*** | -3.16058280799104 | 4.96918723483117 | -6.5805462350037 | 3.23407922899122e-08 | 2.95646671998231e-07 | 8.24703790760494 |
| ***S100A7A*** | 2.76832151066461 | 5.80529484563635 | 6.55993028230002 | 3.47845711520883e-08 | 3.14534102665137e-07 | 8.17470797957622 |
| ***HPGD*** | -2.79979780877984 | 6.49157939902976 | -6.54311819497409 | 3.69134333825707e-08 | 3.32700295681871e-07 | 8.11573182309264 |
| ***EPSTI1*** | 2.49167890669156 | 7.96287895812683 | 6.5072781167877 | 4.1896372120709e-08 | 3.72571262168996e-07 | 7.99003255136206 |
| ***MYH2*** | -4.37733990154507 | 5.52137722238742 | -6.42875547203089 | 5.5288427721021e-08 | 4.74682501093467e-07 | 7.71477846641243 |
| ***TSPAN7*** | -2.6090058171244 | 7.07275062948946 | -6.42231395473235 | 5.65605930609679e-08 | 4.84107522108675e-07 | 7.69220797870912 |
| ***MMP10*** | 4.08596610735742 | 7.78996212898902 | 6.42064069582086 | 5.68958082118269e-08 | 4.86227081362847e-07 | 7.6863452901222 |
| ***CXCL8*** | 2.62854692242054 | 8.28916215143004 | 6.41579930298662 | 5.78769318567862e-08 | 4.93093696678584e-07 | 7.66938284247976 |
| ***MAMDC2*** | -2.20710938484631 | 7.04465097578457 | -6.37388531658973 | 6.71089308104139e-08 | 5.64243488934e-07 | 7.52257077585142 |
| ***MAL*** | -4.47003545099764 | 9.51705410764117 | -6.34562398464685 | 7.41496933437035e-08 | 6.16901744421941e-07 | 7.42362161979526 |

| | | | | | | |
|---|---|---|---|---|---|---|
| ***EEF1A2*** | -2.06125397414874 | 6.116201207692 | -6.32424061960617 | 7.996274905421e-08 | 6.60317078952366e-07 | 7.34877761174537 |
| ***IFI6*** | 2.76011197117757 | 9.4762903438676 | 6.3234823636058 | 8.01770263334936e-08 | 6.61758449261977e-07 | 7.34612402739729 |
| ***THBS4*** | -3.02288912530917 | 6.29970912017982 | -6.26039949284998 | 1.0016506132896e-07 | 8.09485328236368e-07 | 7.12545907223154 |
| ***IDO1*** | 3.28868795998331 | 6.41043964986764 | 6.23208909724825 | 1.10682964759256e-07 | 8.80389427426063e-07 | 7.02649635402835 |
| ***CTSV*** | 2.32487328855566 | 8.93855925369403 | 6.17133271086752 | 1.37120619602083e-07 | 1.06325932965191e-06 | 6.81426901137352 |
| ***MYF6*** | -2.38049412727616 | 4.72557257475065 | -6.0869754738005 | 1.84565696724356e-07 | 1.3866153561754e-06 | 6.51998690480613 |
| ***TYRP1*** | -2.71477304431151 | 4.8778680545418 | -6.07355507653378 | 1.93494358705575e-07 | 1.44522064511213e-06 | 6.47321434777151 |
| ***CRNN*** | -4.45246976699004 | 9.59167088048632 | -6.06116548410204 | 2.02118619178183e-07 | 1.50222566757332e-06 | 6.43004585880184 |
| ***LPL*** | -2.00025600354021 | 6.05597386349988 | -6.05474708989919 | 2.06735853286574e-07 | 1.52903746551562e-06 | 6.40768695990886 |
| ***BST2*** | 2.78458376909564 | 8.2008167962488 | 6.04214759965693 | 2.16107300461471e-07 | 1.58847449094716e-06 | 6.36380470646773 |
| ***CASP14*** | 2.54770497922429 | 5.02789351268293 | 6.01520398098899 | 2.37591357235799e-07 | 1.73111183119837e-06 | 6.27000430614103 |
| ***IL36G*** | 2.66705123148771 | 8.34753958512536 | 5.98954288648591 | 2.60025901802591e-07 | 1.87570005215416e-06 | 6.18072162144739 |
| ***ALDH1A1*** | -2.58369014109258 | 5.99365825512674 | -5.94329848886759 | 3.05909472364757e-07 | 2.17187901607306e-06 | 6.01995933055851 |
| ***IL1B*** | 2.34051967946356 | 8.28354416840212 | 5.87148950577963 | 3.93613829524734e-07 | 2.72034520521327e-06 | 5.77069388668714 |
| ***CKM*** | -3.54967208227128 | 5.62319305881382 | -5.83908573528002 | 4.40983895265513e-07 | 3.02015944060131e-06 | 5.65836918259386 |
| ***MMRN1*** | -2.15534027885331 | 5.23828023726853 | -5.80160819722312 | 5.02877006177368e-07 | 3.38420986460212e-06 | 5.52858484714314 |
| ***LMOD2*** | -2.29162666748815 | 5.14950735521069 | -5.75780813868957 | 5.86225636028021e-07 | 3.89321140098992e-06 | 5.37708756605228 |
| ***MYL1*** | -4.23713440346948 | 6.03132641182728 | -5.67750364454461 | 7.76248630903898e-07 | 4.99003969423402e-06 | 5.09986783466665 |
| ***TNNC1*** | -3.32337038284924 | 5.76684479738028 | -5.67721933135126 | 7.77019874992598e-07 | 4.99114656300683e-06 | 5.09888765371113 |
| ***MUC15*** | -2.38145296727609 | 6.3266423356262 | -5.66762413804035 | 8.03499174579239e-07 | 5.13943250836859e-06 | 5.06581338966249 |
| ***PI3*** | 2.61034985367088 | 12.1410057361531 | 5.65641991300525 | 8.35554274556482e-07 | 5.32605893494557e-06 | 5.02720658183223 |
| ***DPT*** | -2.41228726048179 | 6.97917041307807 | -5.61784629074652 | 9.55930768439265e-07 | 5.99536916928512e-06 | 4.89440744869624 |
| ***TNNT1*** | -2.50047322144978 | 7.06727421683125 | -5.57294803337245 | 1.1178525745847e-06 | 6.91383307919896e-06 | 4.74006670955742 |
| ***FABP4*** | -3.20211285240752 | 6.0432195143926 | -5.49180941406967 | 1.48241861735855e-06 | 8.8657682192905e-06 | 4.46181662068498 |
| ***IFI27*** | 2.27922503690522 | 11.3446349931428 | 5.42771761366255 | 1.8517524064686e-06 | 1.08258294426609e-05 | 4.24267207369891 |
| ***EPHX3*** | -2.35651585251305 | 6.88708695907475 | -5.42692066967929 | 1.85687620008485e-06 | 1.08443653536512e-05 | 4.23995089404076 |
| ***CXCL9*** | 3.51172574617756 | 8.34736972352256 | 5.42508933408468 | 1.86870367128899e-06 | 1.09019714006968e-05 | 4.23369812192675 |

| | | | | | | |
|---|---|---|---|---|---|---|
| ***POSTN*** | 2.00715607136344 | 7.69188227487931 | 5.39747220130019 | 2.05638029705248e-06 | 1.18721213960853e-05 | 4.1394650603974 |
| ***RSAD2*** | 2.68439131078318 | 7.06701184140673 | 5.28550566846595 | 3.02824236366877e-06 | 1.68464945922735e-05 | 3.7586381158359 |
| ***MMP7*** | 2.44243207633765 | 5.64837784344334 | 5.27732385469562 | 3.11491185829576e-06 | 1.72652152784606e-05 | 3.73088945183357 |
| ***SLC6A14*** | 2.40138975832522 | 6.67922110124174 | 5.27502150842922 | 3.13973957779075e-06 | 1.73854728748945e-05 | 3.72308305876202 |
| ***GNLY*** | 2.02461157799497 | 6.19575219821934 | 5.22325696601977 | 3.75258121560987e-06 | 2.03460262783848e-05 | 3.547807854417 |
| ***TMPRSS11B*** | -3.86138730534098 | 6.76354360130201 | -5.16254180584283 | 4.62323559462069e-06 | 2.45550421122456e-05 | 3.34282707677442 |
| ***IFIT3*** | 2.08907097369808 | 7.49953730825425 | 5.14350374808856 | 4.9352748859657e-06 | 2.59557747081291e-05 | 3.27869096800719 |
| ***IFIT2*** | 2.0382973318867 | 6.92881319003976 | 5.12466655882409 | 5.26446136936707e-06 | 2.74272344598617e-05 | 3.2152982489504 |
| ***ENDOU*** | -2.13044352312212 | 6.3043780191719 | -5.08303677519353 | 6.07075770670268e-06 | 3.12081914700123e-05 | 3.07544182923041 |
| ***CSRP3*** | -2.75490247659101 | 5.14578982577969 | -5.07611688041648 | 6.21611254006033e-06 | 3.18570986053061e-05 | 3.05222684571653 |
| ***ALOX12*** | -2.48292083912205 | 6.6797485082754 | -5.05796091578513 | 6.61400335438614e-06 | 3.36683495937211e-05 | 2.99136181168375 |
| ***ABCA12*** | 2.05399255732752 | 6.39776726160443 | 5.03702540389291 | 7.10405378436046e-06 | 3.58669656963649e-05 | 2.92126043679447 |
| ***ACTA1*** | -3.6089550427137 | 6.94131790613176 | -5.03423431361048 | 7.17203378400815e-06 | 3.61882443824416e-05 | 2.91192130492491 |
| ***IFI44L*** | 2.46842873890449 | 7.61798936144832 | 5.00307665504783 | 7.97583601421991e-06 | 3.97146560995058e-05 | 2.80777424009387 |
| ***SLCO1B3*** | 2.27954919749796 | 4.97365905852374 | 4.99130284423492 | 8.30215237040543e-06 | 4.10938038280752e-05 | 2.76847151364149 |
| ***KRT78*** | -2.52404632680908 | 8.13143438302369 | -4.98973433568187 | 8.34660543122457e-06 | 4.12892961278517e-05 | 2.76323778456582 |
| ***MYH1*** | -2.70729979290404 | 4.76783029204238 | -4.92269353555662 | 1.04824247463273e-05 | 5.04027905816476e-05 | 2.54002859818403 |
| ***IL36A*** | -2.0526038524831 | 5.10256334044961 | -4.9211693632142 | 1.05367674854726e-05 | 5.06202478332941e-05 | 2.5349652463925 |
| ***CCL5*** | 2.09242210462389 | 8.29374316744032 | 4.86260342705968 | 1.28485957584794e-05 | 6.04195965424149e-05 | 2.34079947166053 |
| ***FMO2*** | -2.52499977885691 | 6.62685081804097 | -4.85213917009661 | 1.33112532999538e-05 | 6.24014171021758e-05 | 2.30618876442391 |
| ***CLCA4*** | -2.7848801853344 | 7.27518764725467 | -4.81620418202053 | 1.50282455817761e-05 | 6.94727889009333e-05 | 2.18752659986619 |
| ***MYLPF*** | -2.38965157847974 | 6.03579835020563 | -4.74814491972003 | 1.88971977134511e-05 | 8.52523632489822e-05 | 1.96362811030231 |
| ***KRT13*** | -3.14663920177128 | 11.7468128517743 | -4.71048669255463 | 2.14422603116204e-05 | 9.50858061103166e-05 | 1.84023046333108 |
| ***TNNI1*** | -2.54252346448976 | 5.59661967976072 | -4.70492592249891 | 2.18455247234699e-05 | 9.66284589690428e-05 | 1.82203928057338 |
| ***DKK 1,00*** | 2.49052239506722 | 6.25058424805927 | 4.66360026360175 | 2.50847217375278e-05 | 0.000109203116900226 | 1.68709703339992 |
| ***PLA2G2A*** | -2.50395043951046 | 6.39579806757148 | -4.55430723614789 | 3.60933244513381e-05 | 0.00015116178326917 | 1.33239708180682 |
| ***ACTC1*** | -2.61295300187456 | 6.71153887186792 | -4.45991328172971 | 4.9309873192403e-05 | 0.000199068426907651 | 1.02872691921064 |

| | | | | | | |
|---|---|---|---|---|---|---|
| ***CEACAM7*** | -2.39726960697133 | 5.18425623840581 | -4.43867640942212 | 5.28799018684548e-05 | 0.000211621250725849 | 0.96076432017191 |
| ***CWH43*** | -2.03468247379754 | 6.45835021700164 | -4.41574841700246 | 5.7017698632696e-05 | 0.000226394864660532 | 0.88754123634118 |
| ***MB*** | -2.01673167238956 | 5.69017103478786 | -4.39495565274993 | 6.10423997123278e-05 | 0.000240359860427549 | 0.821274939147001 |
| ***ADAMDEC1*** | 2.0040214179425 | 6.17350683044663 | 4.36526572200972 | 6.72736274448299e-05 | 0.000261251932553296 | 0.726883727460066 |
| ***S100P*** | 2.11234336883133 | 8.29427990492246 | 4.35374912046088 | 6.98541685061273e-05 | 0.000269888895995838 | 0.690343534229884 |
| ***SPRR3*** | -2.10927795077265 | 12.4053346114345 | -4.32399825290373 | 7.69754690625072e-05 | 0.000294127876280138 | 0.596142579734323 |
| ***ADH7*** | -2.50909584037733 | 7.4469705992375 | -4.20793856544389 | 0.000112155127228846 | 0.000410922965051401 | 0.231405431209826 |
| ***SERPINB11*** | -2.15438493045492 | 5.03601568411162 | -4.16897707107185 | 0.000127152533281027 | 0.000458906304296595 | 0.10997605661445 |
| ***ALDH3A1*** | -2.12728356831354 | 9.00503717158638 | -4.11242971296743 | 0.000152436379325536 | 0.000538261677771067 | -0.0653208868372319 |
| ***IFIT1*** | 2.17045972734348 | 8.77121842859815 | 4.09354989562703 | 0.000161917287203831 | 0.000567647723777524 | -0.123595084201845 |
| ***NELL2*** | 2.16704427251784 | 7.24984317010403 | 4.08806595514504 | 0.000164776749581 | 0.000576580155676708 | -0.140497642351951 |
| ***STATH*** | -2.02345472943874 | 4.12173564771416 | -4.08623304475066 | 0.000165743350680944 | 0.000579110813707111 | -0.146144594188502 |
| ***CDSN*** | 2.39887743875393 | 7.57880884256124 | 3.80286628415452 | 0.000404113364414229 | 0.00127721294073689 | -1.00376877156459 |
| ***TGM3*** | -2.90132062831351 | 9.80247791531332 | -3.72741842481624 | 0.000509979995213183 | 0.00156604476406172 | -1.22664437979321 |
| ***SPINK7*** | -2.74596806486867 | 8.56236229259054 | -3.30528531737656 | 0.00180017268452947 | 0.00475780327412295 | -2.42571792594062 |

Table S4 - Overlapping DEGs between OLP and OSCC

| Gene | Overlap |
|---|---|
| *TMEM45A* | OLP and early OSCC (overexpressed) |
| *ASPRV1* | OLP and early OSCC (overexpressed) |
| *LCE2B* | OLP and early OSCC (overexpressed) |
| *LCE1B* | OLP and early OSCC (overexpressed) |
| *S100A7* | OLP and early OSCC (overexpressed) |
| *SPRR2G* | OLP and early OSCC (overexpressed) |
| *RNASE7* | OLP and early OSCC (overexpressed) |
| *ALOX12B* | OLP and early OSCC (overexpressed) |
| *KRT16* | OLP and early OSCC (overexpressed) |
| *KRT75* | OLP and early OSCC (overexpressed) |
| *SERPINB7* | OLP and early OSCC (overexpressed) |
| *WFDC5* | OLP and early OSCC (overexpressed) |
| *SPINK6* | OLP and early OSCC (overexpressed) |
| *CA2* | OLP and early OSCC (overexpressed) |
| *WFDC12* | OLP and early OSCC (overexpressed) |
| *KRT10* | OLP and early OSCC (overexpressed) |
| *FLG* | OLP and early OSCC (overexpressed) |

| | |
|---|---|
| *S100A12* | OLP and early OSCC (overexpressed) |
| *IFI27* | OLP and advanced OSCC (overexpressed) |
| *KRT17* | OLP, early OSCC, and advanced OSCC (overexpressed) |
| *PTPRZ1* | OLP, early OSCC, and advanced OSCC (overexpressed) |
| *IL36G* | OLP, early OSCC, and advanced OSCC (overexpressed) |
| *CDSN* | OLP, early OSCC, and advanced OSCC (overexpressed) |
| *S100P* | OLP, early OSCC, and advanced OSCC (overexpressed) |
| *PTHLH* | OLP, early OSCC, and advanced OSCC (overexpressed) |
| *IL1B* | OLP, early OSCC, and advanced OSCC (overexpressed) |
| *PI3* | OLP, early OSCC, and advanced OSCC (overexpressed) |
| *BST2* | OLP, early OSCC, and advanced OSCC (overexpressed) |
| *IFI6* | OLP, early OSCC, and advanced OSCC (overexpressed) |
| *MAMDC2* | OLP and advanced OSCC (underexpressed) |
| *ETNK2* | OLP, early OSCC, and advanced OSCC (underexpressed) |
| *MAOB* | OLP, early OSCC, and advanced OSCC (underexpressed) |
| *PPARGC1A* | OLP, early OSCC, and advanced OSCC (underexpressed) |
| *KRT4* | OLP, early OSCC, and advanced OSCC (underexpressed) |
| *ALOX12* | OLP, early OSCC, and advanced OSCC (underexpressed) |

Table S5 - Enriched Reactome pathways in OLP using GSEA

| Gene Set | Description | Normalized enrichment score | FDR | Genes |
|---|---|---|---|---|
| R-HSA-1474290 | Collagen formation | 2.32 | 0 | *COL12A1, COL15A1, COL16A1, COL1A1, COL1A2, COL22A1, COL3A1, COL4A1, COL4A2, COL4A5, COL4A6, COL5A1, COL5A2, COL6A1, COL6A3, COL7A1, CTSL, CTSS, CTSV, ITGA6, LAMA3, LAMC2, LOX, MMP3, MMP9, P4HA2, PCOLCE, PLOD3, PXDN* |
| R-HSA-6809371 | Formation of the cornified envelope | 2.29 | 0 | *CDSN, DSC1, DSC2, DSG1, FLG, FURIN, IVL, KLK12, KLK13, KLK5, KLK8, KRT1, KRT10, KRT16, KRT17, KRT2, KRT23, KRT24, KRT36, KRT72, KRT73, KRT75, KRT76, KRT80, LCE1B, LCE1E, LCE2B, LCE3D, LOR, PI3, PKP2, RPTN, SPINK6, SPRR1B, SPRR2D, SPRR2G, TGM5* |
| R-HSA-8948216 | Collagen chain trimerization | 2.24 | 0 | *COL15A1, COL16A1, COL1A1, COL1A2, COL22A1, COL3A1, COL4A1, COL4A2, COL4A5, COL5A1, COL5A2, COL6A1, COL6A3, COL7A1* |
| R-HSA-6803157 | Antimicrobial peptides | 2.19 | 0 | *ATOX1, CTSG, DEFB1, LCN2, LYZ, PGLYRP4, PI3, PRSS2, PRSS3, RNASE6, RNASE7, S100A7, S100A7A* |
| R-HSA-909733 | Interferon alpha/beta signaling | 2.00 | 0.001 | *BST2, HLA-A, HLA-B, HLA-C, HLA-E, HLA-F, HLA-G, IFI27, IFI35, IFI6, IFIT1, IFNAR2, IRF4, IRF7, IRF8, IRF9, ISG15, ISG20, JAK1, MX1, MX2, OAS1, OAS2, OAS3, OASL, RSAD2, STAT1, XAF1* |
| R-HSA-1592389 | Activation of Matrix Metalloproteinases | 2.01 | 0.001 | *CTRB2, CTSG, CTSK, CTSV, ELANE, FURIN, MMP1, MMP10, MMP2, MMP25, MMP3, MMP9, PRSS2, TIMP1, TIMP2, TPSAB1* |
| R-HSA-2173782 | Binding and Uptake of Ligands by Scavenger Receptors | 1.97 | 0.002 | *APOE, CD163, CD36, COL1A1, COL1A2, COL3A1, COL4A1, COL4A2, HSPH1, SPARC, STAB1* |
| R-HSA-6798695 | Neutrophil degranulation | 1.94 | 0.004 | *ACAA1, ACLY, ACTR10, ADAM10, ADAM8, ALDOC, ANO6, ANPEP, AOC1, ARG1, ARHGAP9, ARSB, ATP6V0A1, ATP6V1D, B2M, BST2, C3AR1, CAB39, CALML5, CAP1, CAT, CCT2, CCT8, CD14, CD36, CD53, CD93, CDA, CFD, CFP, CHI3L1, CKAP4, CMTM6, COPB1, CRISPLD2, CSNK2B, CST3, CTSA, CTSC, CTSG, CTSH,* |

| | | | | |
|---|---|---|---|---|
| | | | | *CTSS, CTSZ, CXCR2, CYSTM1, DDX3X, DNAJC3, DOCK2, DSC1, DSG1, DYNLL1, ELANE, ENPP4, FABP5, FCER1G, FCGR2A, FCN1, FGL2, FGR, FLG2, FOLR3, FPR1, GGH, GLA, GLB1, GLIPR1, GM2A, GMFG, GNS, GYG1, HEXB, HLA-B, HLA-C, HPSE, HSPA8, IQGAP2, ITGAL, ITGAV, ITGB2, KRT1, LAMP2, LCN2, LRG1, LTF, LYZ, MANBA, METTL7A, MGST1, MMP25, MMP9, NCKAP1L, NEU1, NPC2, NRAS, OLR1, PECAM1, PGAM1, PIGR, PKM, PLAU, PLAUR, PLEKHO2, PNP, PRCP, PRSS2, PRSS3, PRTN3, PSMA5, PSMB7, PSMD12, PSMD14, PSMD2, PSMD6, PYCARD, PYGL, QPCT, QSOX1, RAB18, RAB31, RAB5C, RAB6A, RAP1B, S100A12, S100A7, S100P, SELL, SERPINA1, SERPINA3, SERPINB3, SIRPA, SLC2A3, SLPI, STK10, TBC1D10C, TCN1, TIMP2, TNFRSF1B, TYROBP, UBR4, VAT1, VCP, YPEL5* |
| R-HSA-3000171 | Non-integrin membrane-ECM interactions | 1.86 | 0.019 | *ACTN1, CASK, COL1A1, COL1A2, COL3A1, COL4A1, COL4A2, COL4A5, COL4A6, COL5A1, COL5A2, ITGA6, ITGAV, ITGB1, LAMA2, LAMA3, LAMA4, LAMC2, LAMC3, THBS1, TNC* |
| R-HSA-202433 | Generation of second messenger molecules | 1.83 | 0.021 | *CD247, CD3D, CD4, ENAH, HLA-DPA1, HLA-DPB1, HLA-DQA1, HLA-DQB1, HLA-DQB2, HLA-DRA, HLA-DRB5, ITK, NCK1, PAK1, WAS* |
| R-HSA-6783783 | Interleukin-10 signaling | 1.84 | 0.021 | *CCL19, CCL20, CCL22, CXCL10, IL10RA, IL1A, IL1B, JAK1, PTGS2, TIMP1, TNF, TNFRSF1B* |
| R-HSA-198933 | Immunoregulatory interactions between a Lymphoid and a non-Lymphoid cell | 1.84 | 0.021 | *B2M, CD19, CD1B, CD200, CD247, CD300A, CD300LF, CD300LG, CD3D, CD3G, CD81, CD8A, CD96, CDH1, CLEC2D, COL17A1, COL1A1, COL1A2, COL3A1, HCST, HLA-A, HLA-B, HLA-C, HLA-E, HLA-F, HLA-G, ICAM1, ICAM2, ITGAL, ITGB1, ITGB2, KIR2DL4, KLRB1, KLRG1, LAIR1, LAIR2, LILRB4, LILRB5, OSCAR, PILRA, SELL, SH2D1A, SIGLEC10, SLAMF6, SLAMF7, TREM2, TREML2, TYROBP, VCAM1* |
| R-HSA-2132295 | MHC class II antigen presentation | 1.84 | 0.021 | *ACTR10, AP1S1, AP1S2, AP2S1, ARF1, CANX, CLTC, CTSA, CTSC, CTSK, CTSL, CTSO, CTSS, CTSV, DCTN6, DYNC1I2, HLA-DMB, HLA-DOA, HLA-DOB, HLA-DPA1, HLA-DPB1, HLA-DQA1, HLA-DQB1, HLA-DQB2, HLA-DRB5, IFI30, KIF5B, KIFAP3, KLC1, LGMN, SAR1B, TUBA4A, TUBB2A, TUBB2B, TUBB3* |
| R-HSA-2453864 | Retinoid cycle disease events | 1.82 | 0.027 | *RBP1, RDH12* |
| R-HSA-8963899 | Plasma lipoprotein remodeling | 1.81 | 0.028 | *ABCG1, APOE, CIDEC, FURIN, LCAT, LIPG, LPL, PCSK5, PCSK6, PLTP* |

| | | | | |
|---|---|---|---|---|
| R-HSA-1236975 | Antigen processing-Cross presentation | 1.81 | 0.029 | *B2M, BTK, CALR, CD14, CD207, CD36, CTSL, CTSS, CTSV, CYBB, HLA-A, HLA-B, HLA-C, HLA-E, HLA-F, HLA-G, ITGAV, LY96, MYD88, NCF2, NCF4, PDIA3, PSMA3, PSMA4, PSMA5, PSMA6, PSMB10, PSMB2, PSMB5, PSMB7, PSMC2, PSMC6, PSMD1, PSMD12, PSMD14, PSMD2, PSMD6, PSMD7, PSMD9, PSME3, PSME4, SEC61A2, SEC61B, SEC61G, TAP1, TAP2, TLR1, TLR2, TLR6, VAMP3* |
| R-HSA-166663 | Initial triggering of complement | 1.77 | 0.043 | *C1QB, C1QC, C1S, C2, CFD, CFP, FCN1, FCN3* |

Table S6 - Enriched Reactome pathways in early OSCC using GSEA.

| Gene Set | Description | Normalized enrichment score | FDR | Genes |
|---|---|---|---|---|
| R-HSA-913531 | Interferon Signaling | 2.57 | 0 | *AAAS, ABCE1, ADAR, B2M, BST2, CIITA, DDX58, EIF2AK2, EIF4A2, EIF4A3, FCGR1A, FLNA, FLNB, GBP1, GBP2, GBP3, GBP4, GBP5, HERC5, HLA-A, HLA-B, HLA-C, HLA-DPA1, HLA-DQA1, HLA-DQB1, HLA-DQB2, HLA-DRA, HLA-E, HLA-F, HLA-G, ICAM1, IFI27, IFI30, IFI35, IFI6, IFIT1, IFIT2, IFIT3, IFITM1, IFITM3, IFNAR2, IFNG, IFNGR1, IRF1, IRF4, IRF6, IRF7, IRF8, ISG15, ISG20, JAK2, KPNA2, MX1, MX2, NDC1, NEDD4, NUP107, NUP205, NUP210, NUP37, NUP50, OAS1, OAS2, OAS3, OASL, PLCG1, PML, PSMB8, PTAFR, RNASEL, RSAD2, SAMHD1, SOCS1, SP100, STAT1, STAT2, TRIM14, TRIM21, TRIM22, TRIM25, TRIM38, TRIM5, TRIM6, TYK2, UBA7, UBE2L6, USP18, VCAM1, XAF1* |
| R-HSA-69306 | DNA Replication | 2.38 | 0 | *ANAPC1, ANAPC4, ANAPC5, CCNA1, CCNA2, CCNE1, CCNE2, CDC16, CDC23, CDC45, CDC6, CDC7, CDK2, CDT1, DNA2, E2F3, FEN1, GINS1, GINS2, GINS3, GINS4, GMNN, LIG1, MCM10, MCM2, MCM3, MCM4, MCM5, MCM6, MCM7, MCM8, ORC2, ORC5, ORC6, PCNA, POLA1, POLA2, POLD1, POLD3, POLE2, POLE3, PRIM1, PRIM2, PSMA1, PSMA2, PSMA3, PSMA4, PSMA5, PSMA6, PSMB1, PSMB10, PSMB2, PSMB3, PSMB4, PSMB7, PSMB8, PSMB9, PSMC2, PSMC4, PSMC6, PSMD10, PSMD11, PSMD14, PSMD2, PSMD3, PSMD5, PSMD6, PSMD9, PSME1, PSME2, PSME3, PSMF1, RFC1, RFC2, RFC3, RFC4, RFC5, RPA3,* |

| | | | | |
|---|---|---|---|---|
| | | | | *RPS27A, SKP2, UBE2C, UBE2D1* |
| R-HSA-1236975 | Antigen processing-Cross presentation | 2.34 | 0 | *B2M, BTK, CALR, CD14, CHUK, CTSL, CTSS, CTSV, CYBA, CYBB, FCGR1A, HLA-A, HLA-B, HLA-C, HLA-E, HLA-F, HLA-G, ITGAV, LY96, MRC1, MYD88, NCF2, NCF4, PDIA3, PSMA1, PSMA2, PSMA3, PSMA4, PSMA5, PSMA6, PSMB1, PSMB10, PSMB2, PSMB3, PSMB4, PSMB7, PSMB8, PSMB9, PSMC2, PSMC4, PSMC6, PSMD10, PSMD11, PSMD14, PSMD3, PSMD5, PSMD6, PSMD9, PSME1, PSME2, PSME3, RPS27A, SEC61A1, STX4, TAP1, TAP2, TAPBP, TLR1, TLR2, TLR6* |
| R-HSA-68882 | Mitotic Anaphase | 2.28 | 0 | *AHCTF1, ANAPC1, ANAPC4, ANAPC5, ANKLE2, AURKB, BIRC5, BUB1, BUB1B, BUB3, CDC16, CDC20, CDC23, CDCA5, CDCA8, CENPA, CENPC, CENPE, CENPF, CENPI, CENPK, CENPM, CENPN, CENPO, CENPQ, CENPT, CKAP5, DSN1, DYNC1I2, DYNC1LI1, DYNLL1, ERCC6L, KIF18A, KIF2A, KIF2C, KNTC1, LMNB1, MAD2L1, MIS12, NDC80, NDE1, NDEL1, NUF2, NUP107, NUP160, NUP37, NUP43, PLK1, PPP2R5B, PSMA1, PSMA2, PSMA3, PSMA4, PSMA5, PSMA6, PSMB1, PSMB10, PSMB2, PSMB3, PSMB4, PSMB7, PSMB8, PSMB9, PSMC2, PSMC4, PSMC6, PSMD10, PSMD11, PSMD14, PSMD2, PSMD3, PSMD5, PSMD6, PSMD7, PSMD9, PSME1, PSME2, PSME3, PSMF1, PTTG1, RAD21, RANGAP1, RCC2, RPS27A, SKA1, SKA2, SMC1A, SMC3, SPC25, SPDL1, STAG2, TMPO, TUBA1C, UBE2C, UBE2D1, VRK1, VRK2, XPO1, ZW10, ZWILCH, ZWINT* |
| R-HSA-174184 | Cdc20:Phospho-APC/C mediated degradation of Cyclin A | 2.20 | 0 | *ANAPC1, ANAPC4, ANAPC5, BUB1B, BUB3, CCNA1, CCNA2, CDC16, CDC20, CDC23, CDK1, MAD2L1, PSMA1, PSMA2, PSMA3, PSMA4, PSMA5, PSMA6, PSMB1, PSMB10, PSMB2, PSMB3, PSMB4, PSMB7, PSMB8, PSMB9, PSMC2, PSMC4, PSMC6, PSMD10, PSMD11, PSMD14, PSMD2, PSMD3, PSMD5, PSMD6, PSMD7, PSMD9, PSME1, PSME2, PSME3, PSMF1, RPS27A, UBE2C, UBE2D1* |
| R-HSA-141424 | Amplification of signal from the kinetochores | 2.17 | 0 | *AURKB, BIRC5, BUB1, BUB1B, BUB3, CDC20, CDCA8, CENPA, CENPC, CENPE, CENPF, CENPI, CENPK, CENPM, CENPN, CENPO, CENPQ, CENPT, CKAP5, DSN1, DYNC1LI1, DYNLL1, ERCC6L, KIF18A, KIF2A, KIF2C, KNTC1, MAD2L1, NDC80, NDE1, NUF2, NUP107, NUP37, PLK1, PPP2R5B, RANGAP1, RCC2, SKA1, SKA2, SPC25, SPDL1, XPO1, ZW10,* |

| | | | | |
|---|---|---|---|---|
| | | | | *ZWILCH, ZWINT* |
| R-HSA-5693567 | HDR through Homologous Recombination (HRR) or Single Strand Annealing (SSA) | 2.13 | 0 | *ATM, ATR, BLM, BRCA1, BRCA2, BRCC3, BRIP1, CCNA1, CCNA2, CDK2, CHEK1, CLSPN, DNA2, EME1, EME2, ERCC4, EXO1, GEN1, MDC1, NBN, PALB2, PCNA, POLD1, POLD2, POLD3, POLE2, POLE3, POLE4, POLK, PPP4C, RAD1, RAD17, RAD50, RAD51, RAD51AP1, RAD51C, RAD9A, RBBP8, RFC1, RFC2, RFC3, RFC4, RFC5, RHNO1, RMI1, RMI2, RPA1, RPA3, RPS27A, SIRT6, SLX4, SPIDR, TIMELESS, TIPIN, TOPBP1, UBE2I, UBE2V2, XRCC2, XRCC3* |
| R-HSA-202430 | Translocation of ZAP-70 to Immunological synapse | 2.11 | 0 | *CD247, CD3D, CD3E, CD3G, CD4, HLA-DPA1, HLA-DQA1, HLA-DQB1, HLA-DQB2, HLA-DRA, LCK, PTPN22, ZAP70* |
| R-HSA-69275 | G2/M Transition | 2.10 | 0 | *AJUBA, AURKA, BORA, CCNA1, CCNA2, CCNB1, CCNB2, CCP110, CDC25A, CDC25B, CDC25C, CDK1, CDK2, CDK7, CDKN1A, CENPF, CENPJ, CEP135, CEP152, CEP192, CEP250, CEP290, CEP57, CEP78, CKAP5, CNTRL, DCTN3, DYNC1I2, DYNLL1, EP300, FOXM1, HAUS1, HAUS3, HAUS6, HAUS7, HMMR, HSP90AA1, HSP90AB1, LIN9, MYBL2, NDE1, NEDD1, NEK2, NINL, OFD1, OPTN, PKMYT1, PLK1, PLK4, PPME1, PSMA1, PSMA2, PSMA3, PSMA4, PSMA5, PSMA6, PSMB1, PSMB10, PSMB2, PSMB3, PSMB4, PSMB7, PSMB8, PSMB9, PSMC2, PSMC4, PSMC6, PSMD1, PSMD10, PSMD11, PSMD12, PSMD14, PSMD2, PSMD3, PSMD5, PSMD6, PSMD7, PSMD9, PSME1, PSME2, PSME3, PSMF1, RAB8A, RPS27A, SDCCAG8, TPX2, TUBA1C, TUBA4A, TUBB3, TUBG1, TUBG2, TUBGCP3, TUBGCP5, XPO1, YWHAE* |
| R-HSA-397014 | Muscle contraction | -2.89 | 0 | *ABCC9, ACTA1, ACTA2, ACTC1, ACTG2, ACTN2, ANXA6, ATP1A2, ATP1B1, ATP1B2, ATP2A1, ATP2B2, CACNA1D, CACNA1S, CACNA2D1, CACNA2D3, CACNB1, CACNB4, CAMK2B, CASQ1, CASQ2, CORIN, DES, DMD, DMPK, FGF12, FGF14, FXYD1, FXYD2, FXYD6, HIPK2, KCNE4, KCNIP2, KCNIP3, KCNJ12, KCNK5, LMOD1, MYBPC1, MYBPC2, MYH11, MYH3, MYH6, MYH8, MYL1, MYL2, MYL3, MYL6B, MYL9, MYLPF, NEB, NOS1, NPR1, PLN, PRKACA, RYR1, SCN1B, SCN2B, SCN4A, SCN7A, SCN9A, SLC8A3, SLN, SORBS1, TCAP, TMOD1, TMOD4, TNNC1, TNNC2, TNNI1, TNNI2, TNNT1, TNNT3, TPM2, TPM3, TRDN, TRPC1, TTN* |

| | | | | |
|---|---|---|---|---|
| R-HSA-5693579 | Homologous DNA Pairing and Strand Exchange | 02.05 | 0 | *ATR, BLM, BRCA1, BRCA2, BRIP1, CHEK1, DNA2, EXO1, RAD50, RAD51, RAD51AP1, RAD51C, RAD9A, RBBP8, RFC2, RFC3, RFC4, RFC5, RMI1, TOPBP1, XRCC2* |
| R-HSA-211733 | Regulation of activated PAK-2p34 by proteasome mediated degradation | 02.06 | 0 | *PSMA1, PSMA2, PSMA3, PSMA4, PSMA5, PSMA6, PSMB1, PSMB10, PSMB2, PSMB3, PSMB4, PSMB7, PSMB8, PSMB9, PSMC2, PSMC4, PSMC6, PSMD1, PSMD10, PSMD11, PSMD12, PSMD14, PSMD2, PSMD3, PSMD5, PSMD6, PSMD7, PSMD9, PSME1, PSME2, PSME3, PSMF1, RPS27A* |
| R-HSA-1236977 | Endosomal/Vacuolar pathway | 2.00 | 0 | *B2M, CTSL, CTSS, CTSV, HLA-A, HLA-B, HLA-C, HLA-E, HLA-F, HLA-G* |
| R-HSA-68962 | Activation of the pre-replicative complex | 1.98 | 0 | *CDC45, CDC6, CDC7, CDK2, CDT1, GMNN, MCM10, MCM2, MCM3, MCM4, MCM5, MCM6, MCM7, MCM8, ORC2, ORC5, ORC6, POLA1, POLA2, POLE2, POLE3, PRIM1, PRIM2, RPA3* |
| R-HSA-6785807 | Interleukin-4 and Interleukin-13 signaling | 1.97 | 0 | *BATF, CCL22, CXCL8, FASLG, FN1, FSCN1, GATA3, HIF1A, HMOX1, HSP90B1, ICAM1, IL13RA2, IL18, IL1A, IL1B, IL2RG, IL4R, IL6, IRF4, ITGAM, ITGAX, ITGB2, JAK2, JAK3, LCN2, MMP1, MMP3, MMP9, SOCS1, STAT1, TIMP1, TNF, TNFRSF1B, TYK2, VCAM1* |
| R-HSA-6809371 | Formation of the cornified envelope | 1.94 | 0 | *CASP14, CDSN, DSC1, DSC3, DSG1, DSG2, DSG3, FLG, KLK5, KLK8, KRT1, KRT10, KRT14, KRT16, KRT17, KRT23, KRT24, KRT5, KRT6A, KRT6B, KRT75, KRT80, KRT9, LCE1B, LCE2B, LCE3D, LOR, PI3, SPINK6, SPRR1B, SPRR2D, SPRR2G* |
| R-HSA-109581 | Apoptosis | 1.91 | 0.001 | *APAF1, BAK1, BBC3, BCAP31, BCL2L1, BCL2L11, BID, BIRC2, BMF, CASP3, CASP6, CASP7, CASP8, CD14, CLSPN, CTNNB1, DAPK1, DIABLO, DNM1L, DSG1, DSG2, DSG3, DSP, DYNLL1, FAS, FASLG, FNTA, GZMB, HMGB2, KPNA1, KPNB1, LMNB1, LY96, NMT1, PMAIP1, PSMA1, PSMA2, PSMA3, PSMA4, PSMA5, PSMA6, PSMB1, PSMB10, PSMB2, PSMB3, PSMB4, PSMB7, PSMB8, PSMB9, PSMC2, PSMC4, PSMC6, PSMD1, PSMD10, PSMD11, PSMD12, PSMD14, PSMD2, PSMD3, PSMD5, PSMD6, PSMD7, PSMD9, PSME1, PSME2, PSME3, PSMF1, RIPK1, RPS27A, SFN, STAT3, TJP2, TNFRSF10B, TNFSF10, TP63, TP73, TRADD, UNC5B, XIAP, YWHAE* |

| | | | | |
|---|---|---|---|---|
| R-HSA-287183 7 | FCERI mediated NF-kB activation | 1.91 | 0.001 | *CARD11, CHUK, FCER1G, IKBKB, LYN, MALT1, NFKB1, NFKBIA, PSMA1, PSMA2, PSMA3, PSMA4, PSMA5, PSMA6, PSMB1, PSMB10, PSMB2, PSMB3, PSMB4, PSMB7, PSMB8, PSMB9, PSMC2, PSMC4, PSMC6, PSMD1, PSMD10, PSMD11, PSMD12, PSMD14, PSMD2, PSMD3, PSMD5, PSMD6, PSMD7, PSMD9, PSME1, PSME2, PSME3, PSMF1, RASGRP1, RELA, RPS27A, UBE2D1* |
| R-HSA-110313 | Translesion synthesis by Y family DNA polymerases bypasses lesions on DNA template | 1.91 | 0.001 | *ISG15, NPLOC4, PCNA, POLD1, POLD3, POLE2, POLE3, POLK, RFC1, RFC2, RFC3, RFC4, RFC5, RPA3, RPS27A, TRIM25, UBA7, UBE2L6, VCP* |
| R-HSA-569353 7 | Resolution of D-Loop Structures | 1.89 | 0.001 | *BLM, BRCA1, BRCA2, BRIP1, DNA2, EME1, EME2, EXO1, GEN1, NBN, PALB2, RAD50, RAD51, RAD51AP1, RAD51C, RBBP8, RMI1, SLX4, SPIDR, XRCC2, XRCC3* |
| R-HSA-566903 4 | TNFs bind their physiological receptors | 1.89 | 0.001 | *CD27, CD70, EDARADD, FASLG, TNFRSF11B, TNFRSF17, TNFRSF18, TNFRSF1B, TNFRSF9, TNFSF13B, TNFSF15, TNFSF18, TNFSF4, TNFSF9* |
| R-HSA-69205 | G1/S-Specific Transcription | 1.88 | 0.001 | *CCNA1, CCNE1, CDC45, CDC6, CDK1, DHFR, E2F4, FBXO5, HDAC1, PCNA, RBL1, RRM2, TK1, TYMS* |
| R-HSA-606279 | Deposition of new CENPA-containing nucleosomes at the centromere | 1.88 | 0.001 | *CENPA, CENPC, CENPI, CENPK, CENPM, CENPN, CENPO, CENPQ, CENPT, CENPW, HJURP, MIS18A, MIS18BP1, OIP5, RSF1, RUVBL1* |
| R-HSA-162587 | HIV Life Cycle | 1.88 | 0.001 | *AAAS, CCNK, CCNT1, CCNT2, CCR5, CD4, CDK7, CHMP2B, CHMP4A, CHMP4C, CHMP5, CXCR4, ERCC3, FEN1, FURIN, GTF2A1, GTF2A2, GTF2B, GTF2E1, GTF2E2, GTF2F2, GTF2H3, GTF2H4, HMGA1, KPNA1, LIG1, LIG4, NCBP1, NCBP2, NDC1, NEDD4L, NELFA, NELFCD, NMT1, NUP107, NUP133, NUP153, NUP155, NUP160, NUP188, NUP205, NUP210, NUP214, NUP35, NUP37, NUP43, NUP50, NUP54, NUP85, NUP88, NUP93, PDCD6IP, POLR2A, POLR2C, POLR2D, POLR2E, POLR2F, POLR2G, POLR2H, POLR2J, POLR2K, PPIA, PSIP1, RAE1, RAN, RANBP1, RANGAP1, RCC1, RNGTT, RPS27A, SSRP1, TAF11, TAF13, TAF2, TAF4B, TAF5, TBP, TCEA1, TPR, TSG101, UBAP1, VPS37C, VPS37D, VTA1, XPO1, XRCC4, XRCC6* |
| R-HSA-561078 | Degradation of GLI1 by the | 1.88 | 0.001 | *PSMA1, PSMA2, PSMA3, PSMA4, PSMA5, PSMA6, PSMB1, PSMB10,* |

| | | | | |
|---|---|---|---|---|
| 0 | proteasome | | | *PSMB2, PSMB3, PSMB4, PSMB7, PSMB8, PSMB9, PSMC2, PSMC4, PSMC6, PSMD10, PSMD11, PSMD14, PSMD2, PSMD3, PSMD5, PSMD6, PSMD7, PSMD9, PSME1, PSME2, PSME3, PSMF1, RPS27A* |
| R-HSA-174417 | Telomere C-strand (Lagging Strand) Synthesis | 1.87 | 0.001 | *DNA2, FEN1, LIG1, PCNA, POLA1, POLA2, POLD1, POLD3, POLE2, POLE3, PRIM1, PRIM2, RFC1, RFC2, RFC3, RFC4, RFC5, RPA3* |
| R-HSA-211945 | Phase I - Functionalization of compounds | -2.15 | 0.001 | *AADAC, ACSS2, ADH1A, ADH1B, ADH1C, ADH4, ADH7, ALDH1A1, ALDH2, ALDH3A1, AOC3, ARNT, BPHL, CES1, CES2, CYP11A1, CYP19A1, CYP1B1, CYP27A1, CYP2C18, CYP2C19, CYP2F1, CYP2J2, CYP2R1, CYP2U1, CYP3A4, CYP3A5, CYP46A1, CYP4B1, CYP4F12, CYP4F22, CYP4F3, CYP7B1, EPHX1, FMO2, MAOB, MARC1, MARC2, PTGIS* |
| R-HSA-195258 | RHO GTPase Effectors | 1.85 | 0.002 | *ACTR3, AHCTF1, ARPC1A, ARPC1B, ARPC2, ARPC4, ARPC5, AURKB, BIRC5, BTK, BUB1, BUB1B, BUB3, CALM3, CDC20, CDC25C, CDCA8, CENPA, CENPC, CENPE, CENPF, CENPI, CENPK, CENPL, CENPM, CENPN, CENPO, CENPQ, CENPT, CENPU, CFL1, CIT, CKAP5, CTNNB1, CYBA, CYBB, CYFIP1, CYFIP2, DIAPH1, DIAPH3, DSN1, DYNC1I2, DYNC1LI1, DYNLL1, ERCC6L, EVL, FLNA, FMNL1, FMNL2, FMNL3, GOPC, GRB2, IQGAP3, ITGB1, ITGB3BP, KDM1A, KIF14, KIF18A, KIF2A, KIF2C, KLC2, KNTC1, KTN1, LIMK1, LIMK2, MAD2L1, MAPRE1, MIS12, MYL6, NCF1, NCF2, NCF4, NCK1, NCKAP1L, NDC80, NDE1, NDEL1, NUF2, NUP107, NUP133, NUP160, NUP37, NUP43, NUP85, PAFAH1B1, PAK1, PFN1, PIK3R4, PLK1, PPP1CC, PPP2CA, PPP2R5B, PRC1, PTK2, RAC1, RAC2, RANGAP1, RCC2, RHOC, RHOG, RHPN2, RTKN, SFN, SKA1, SKA2, SPC25, SPDL1, SRGAP2, TUBA1C, TUBA4A, TUBB3, WAS, WASL, WIPF1, WIPF2, XPO1, YWHAB, YWHAE, ZW10, ZWILCH, ZWINT* |
| R-HSA-72172 | mRNA Splicing | 1.84 | 0.002 | *ALYREF, AQR, CASC3, CCAR1, CD2BP2, CLP1, CPSF1, CPSF3, CPSF7, CRNKL1, CSTF1, CSTF2, CSTF3, CTNNBL1, CWC15, CWC27, DDX23, DDX46, DDX5, DHX15, DHX16, DHX9, DNAJC8, EFTUD2, EIF4A3, ELAVL2, FUS, GTF2F2, HNRNPA1, HNRNPA2B1, HNRNPA3, HNRNPC, HNRNPD, HNRNPF, HNRNPH1, HNRNPK, HNRNPM, HNRNPR, HNRNPU, HSPA8, ISY1, LSM2, LSM3, LSM5, LSM6, LSM7, LSM8, MAGOH, MAGOHB, NCBP1, NCBP2, PAPOLA, PHF5A, PLRG1, POLR2A, POLR2C, POLR2D, POLR2E, POLR2F, POLR2G, POLR2H, POLR2J, POLR2K, PPIE,* |

| | | | | |
|---|---|---|---|---|
| | | | | *PPIH, PPIL1, PPIL4, PPWD1, PRCC, PRPF19, PRPF3, PRPF4, PRPF40A, PTBP1, RBM22, RBM5, RBMX, RNPS1, SART1, SF3B2, SF3B3, SF3B4, SF3B6, SLU7, SMNDC1, SNRNP40, SNRNP48, SNRPA, SNRPA1, SNRPB, SNRPB2, SNRPC, SNRPD1, SNRPD2, SNRPF, SNRPG, SRRT, SRSF1, SRSF10, SRSF11, SRSF2, SRSF3, SRSF4, SRSF6, SRSF7, SRSF9, SUGP1, TFIP11, TRA2B, U2AF1, U2AF1L4, U2AF2, U2SURP, UPF3B, WBP4, ZCRB1, ZRSR2* |
| R-HSA-918233 | TRAF3-dependent IRF activation pathway | 1.84 | 0.002 | *DDX58, EP300, IFIH1, IKBKE, IRF3, IRF7, MAVS, TBK1, TRAF3, TRIM25* |
| R-HSA-1474228 | Degradation of the extracellular matrix | 1.83 | 0.002 | *ADAM10, ADAM9, CASP3, COL12A1, COL17A1, COL4A1, COL4A2, COL4A5, COL7A1, CTSB, CTSL, CTSS, CTSV, FN1, KLK7, LAMA3, LAMB3, LAMC2, MMP1, MMP10, MMP11, MMP12, MMP13, MMP14, MMP3, MMP9, SPP1* |
| R-HSA-2029481 | FCGR activation | 1.83 | 0.002 | *CD247, CD3G, FCGR1A, FCGR2A, FYN, HCK, LYN* |
| R-HSA-2424491 | DAP12 signaling | 1.82 | 0.003 | *B2M, BTK, FYN, GRAP2, GRB2, HLA-E, KLRC2, KLRD1, KRAS, LAT, LCK, LCP2, PLCG1, RAC1, SYK, TREM2, TYROBP, VAV2, VAV3* |
| R-HSA-69109 | Leading Strand Synthesis | 1.79 | 0.004 | *PCNA, POLA2, POLD1, POLD3, PRIM1, PRIM2, RFC1, RFC2, RFC3, RFC4, RFC5* |
| R-HSA-5696397 | Gap-filling DNA repair synthesis and ligation in GG-NER | 1.79 | 0.004 | *LIG1, PCNA, POLD1, POLD2, POLD3, POLE2, POLE3, POLE4, POLK, RFC1, RFC2, RFC3, RFC4, RFC5, RPA1, RPA3, RPS27A* |
| R-HSA-3000157 | Laminin interactions | 1.79 | 0.004 | *COL4A1, COL4A2, COL4A5, COL4A6, COL7A1, ITGA2, ITGA3, ITGA6, ITGB4, LAMA1, LAMA3, LAMB1, LAMB3, LAMC2* |
| R-HSA-162658 | Golgi Cisternae Pericentriolar Stack Reorganization | 1.78 | 0.005 | *CCNB1, CCNB2, CDK1, PLK1* |
| R-HSA-6791312 | TP53 Regulates Transcription of Cell Cycle Genes | 1.78 | 0.005 | *ARID3A, AURKA, CCNA1, CCNA2, CCNB1, CCNE1, CCNE2, CDC25C, CDK1, CDK2, CDKN1A, CENPJ, CNOT6, E2F4, E2F7, E2F8, EP300, PCNA, PLK2, RBL1, SFN, TNKS1BP1, ZNF385A* |
| R-HSA-392154 | Nitric oxide stimulates guanylate | -2.05 | 0.005 | *KCNMA1, NOS1, PDE11A, PDE1A, PDE1B, PDE2A, PDE3A, PDE3B,* |

| | | | | |
|---|---|---|---|---|
| | cyclase | | | *PDE5A, PDE9A, PRKG1, PRKG2* |
| R-HSA-5218859 | Regulated Necrosis | 1.76 | 0.006 | *BIRC2, BIRC3, CASP8, FAS, FASLG, MLKL, RIPK3, RPS27A, TNFRSF10B, TNFSF10* |
| R-HSA-2559583 | Cellular Senescence | 1.75 | 0.007 | *ACD, ANAPC1, ANAPC10, ANAPC11, ANAPC4, ANAPC5, ATM, CABIN1, CBX8, CCNA1, CCNA2, CCNE1, CCNE2, CDC16, CDC23, CDC26, CDC27, CDK2, CDK4, CDK6, CDKN1A, CDKN2B, CDKN2C, CEBPB, CXCL8, E2F2, E2F3, EED, ETS1, EZH2, HIRA, HMGA1, HMGA2, ID1, IGFBP7, IL1A, IL6, KDM6B, LMNB1, MAP2K3, MAP2K4, MAP3K5, MAP4K4, MAPK10, MAPK14, MAPK9, MAPKAPK2, MAPKAPK5, MDM2, MDM4, MINK1, MOV10, NBN, NFKB1, POT1, RAD50, RB1, RBBP7, RELA, RNF2, RPS27A, RPS6KA1, RPS6KA3, STAT3, SUZ12, TERF1, TERF2IP, TFDP1, TINF2, TNRC6A, UBE2C, UBE2D1, UBN1* |
| R-HSA-622312 | Inflammasomes | 1.74 | 0.008 | *AIM2, APP, BCL2L1, CASP1, HSP90AB1, MEFV, NFKB1, NFKB2, NLRP1, NLRP3, P2RX7, PANX1, PSTPIP1, PYCARD, RELA, SUGT1* |
| R-HSA-71240 | Tryptophan catabolism | 1.74 | 0.008 | *IDO1, IDO2, KMO, KYNU, SLC3A2, SLC7A5, TDO2* |
| R-HSA-9020591 | Interleukin-12 signaling | 1.74 | 0.008 | *ANXA2, BOLA2, CAPZA1, CFL1, CNN2, GSTO1, HNRNPA2B1, HNRNPF, IFNG, IL12RB1, IL12RB2, JAK1, JAK2, LCP1, LMNB1, MIF, MSN, MTAP, PSME2, RALA, RAP1B, SERPINB2, SNRPA1, SOD2, STAT4, TYK2* |
| R-HSA-375276 | Peptide ligand-binding receptors | 1.74 | 0.008 | *APP, C3AR1, C5AR1, CCL20, CCL5, CCR1, CCR5, CCR6, CCR7, CXCL1, CXCL10, CXCL11, CXCL13, CXCL16, CXCL2, CXCL3, CXCL6, CXCL8, CXCL9, CXCR1, CXCR4, EDN1, F2R, F2RL1, F2RL2, FPR3, PPBP, XCL1* |
| R-HSA-380259 | Loss of Nlp from mitotic centrosomes | 1.73 | 0.009 | *ACTR1A, AKAP9, CCP110, CDK1, CENPJ, CEP135, CEP152, CEP164, CEP192, CEP250, CEP290, CEP57, CEP70, CEP78, CETN2, CKAP5, CLASP1, CNTRL, CSNK1E, DCTN3, DYNC1I2, DYNLL1, FGFR1OP, HAUS1, HAUS3, HAUS5, HAUS6, HAUS7, HSP90AA1, MAPRE1, NDE1, NEDD1, NEK2, NINL, OFD1, PAFAH1B1, PLK1, PLK4, SDCCAG8, TUBA4A, TUBB, TUBB4B, TUBG1, YWHAE* |
| R-HSA-1500620 | Meiosis | 1.71 | 0.011 | *ACD, ATM, ATR, BLM, BRCA1, BRCA2, CDK2, DIDO1, LMNB1, MLH1, MND1, MSH5, NBN, POT1, PSMC3IP, RAD21, RAD50, RAD51, RAD51C, RBBP8, REC8, RPA1, RPA3, SMC1A, SMC3, STAG1, STAG2, TERF1,* |

| | | | | |
|---|---|---|---|---|
| | | | | *TERF2IP, TINF2, UBE2I* |
| R-HSA-8852135 | Protein ubiquitination | 1.70 | 0.012 | *DERL1, HLA-A, LEO1, OTULIN, PCNA, PEX2, PRKDC, RAD18, RNF152, RPS27A, SHPRH, UBA6, UBE2A, UBE2C, UBE2D1, UBE2G1, UBE2L3, UBE2R2, UBE2S, UBE2T, UBE2V2, UBE2Z, UCHL3, VCP* |
| R-HSA-173623 | Classical antibody-mediated complement activation | 1.70 | 0.012 | *C1QA, C1QB, C1QC, C1R, C1S* |
| R-HSA-1606322 | ZBP1(DAI) mediated induction of type I IFNs | 1.70 | 0.012 | *CHUK, DHX9, DTX4, IKBKB, IRF3, MYD88, NFKB1, NFKB2, NFKBIA, NFKBIB, NKIRAS2, RELA, RIPK1, RIPK3, TBK1, ZBP1* |
| R-HSA-170822 | Regulation of Glucokinase by Glucokinase Regulatory Protein | 1.70 | 0.012 | *AAAS, NDC1, NUP107, NUP133, NUP153, NUP155, NUP160, NUP188, NUP205, NUP210, NUP214, NUP35, NUP37, NUP43, NUP50, NUP54, NUP85, NUP88, NUP93, RAE1, TPR* |
| R-HSA-6811434 | COPI-dependent Golgi-to-ER retrograde traffic | 1.70 | 0.013 | *ARCN1, ARFGAP1, CENPE, COPB1, COPB2, GBF1, KIF11, KIF15, KIF16B, KIF18A, KIF18B, KIF20A, KIF20B, KIF22, KIF23, KIF2A, KIF2C, KIF4A, KIFAP3, KIFC1, KLC2, NAPB, RACGAP1, TMED9, TUBA1C, ZW10* |
| R-HSA-209952 | Peptide hormone biosynthesis | 1.70 | 0.013 | *INHBA, PCSK1* |
| R-HSA-877312 | Regulation of IFNG signaling | 1.69 | 0.014 | *IFNG, IFNGR1, JAK2, PTPN1, PTPN2, PTPN6, SOCS1, STAT1* |
| R-HSA-202733 | Cell surface interactions at the vascular wall | 1.68 | 0.015 | *ANGPT2, ATP1B3, BSG, CAV1, CD177, CD2, CD244, CD47, CD74, CD84, COL1A1, DOK2, FCER1G, FN1, FYN, GAS6, GRB2, GRB7, INPP5D, ITGA3, ITGA4, ITGA6, ITGAL, ITGAM, ITGAV, ITGAX, ITGB2, KRAS, LCK, LYN, MERTK, MMP1, OLR1, PLCG1, PROCR, PSG4, PSG5, PTPN6, SDC4, SELE, SELL, SELP, SELPLG, SIRPA, SIRPG, SLC16A1, SLC16A3, SLC3A2, SLC7A11, SLC7A5, SLC7A7, SLC7A8, TNFRSF10B, TREM1* |
| R-HSA-448706 | Interleukin-1 processing | 1.67 | 0.017 | *CASP1, IL18, IL1A, IL1B, NFKB1, NFKB2* |
| R-HSA-6781827 | Transcription-Coupled Nucleotide Excision Repair (TC-NER) | 1.67 | 0.017 | *AQR, CDK7, COPS5, CUL4B, EP300, ERCC3, ERCC4, GTF2H3, GTF2H4, HMGN1, ISY1, LIG1, PCNA, POLD1, POLD2, POLD3, POLD4, POLE2, POLE3, POLE4, POLK, POLR2A, POLR2C, POLR2D, POLR2E, POLR2F, POLR2G, POLR2H, POLR2J, POLR2K, PPIE, PRPF19, RBX1, RFC1, RFC2, RFC3, RFC4, RFC5, RPA1, RPA3, RPS27A, TCEA1* |
| R-HSA-109688 | Cleavage of Growing Transcript in | 1.66 | 0.018 | *ALYREF, CASC3, CDC40, CHTOP, CLP1, CPSF1, CPSF3, CPSF7, CSTF1,* |

| | | | | |
|---|---|---|---|---|
| | the Termination Region | | | *CSTF2, CSTF3, DDX39A, EIF4A3, MAGOH, MAGOHB, NCBP1, NCBP2, PAPOLA, RNPS1, SLBP, SLU7, SNRPB, SNRPF, SNRPG, SRSF1, SRSF11, SRSF2, SRSF3, SRSF4, SRSF6, SRSF7, SRSF9, SYMPK, THOC2, THOC3, THOC6, THOC7, U2AF1, U2AF1L4, U2AF2, UPF3B, ZC3H11A* |
| R-HSA-912526 | Interleukin receptor SHC signaling | 1.66 | 0.020 | *CSF2, CSF2RA, CSF2RB, IL2RA, IL2RB, IL2RG, IL3RA, JAK2, JAK3* |
| R-HSA-8868773 | rRNA processing in the nucleus and cytosol | 1.66 | 0.020 | *C1D, CSNK1E, DCAF13, DDX21, DDX47, DDX52, DHX37, DIS3, DKC1, EBNA1BP2, EMG1, ERI1, EXOSC10, EXOSC2, EXOSC3, EXOSC4, EXOSC7, EXOSC8, EXOSC9, FCF1, FTSJ3, GNL3, HEATR1, ISG20L2, LAS1L, MPHOSPH6, NAT10, NCL, NIP7, NOB1, NOL11, NOL6, NOP10, NOP14, NOP2, NOP56, NOP58, PDCD11, PES1, PNO1, PWP2, RBM28, RCL1, RIOK1, RIOK2, RPL17, RPL22L1, RPL23, RPL28, RPL31, RPL32, RPL34, RPL35, RPL36AL, RPL37, RPL39, RPL39L, RPP40, RPS15A, RPS19, RPS20, RPS24, RPS25, RPS26, RPS27A, RPS27L, RPS3, RPS4Y1, RPS7, RPSA, RRP36, RRP7A, SENP3, TEX10, THUMPD1, TRMT112, TSR1, UTP14A, UTP15, UTP20, UTP3, UTP6, WDR12, WDR3, WDR75, XRN2* |
| R-HSA-15869 | Metabolism of nucleotides | 1.65 | 0.021 | *ADA, ADK, AMPD3, ATIC, CDA, DCK, DTYMK, ENTPD1, ENTPD7, GART, GDA, GLRX, HPRT1, ITPA, NME1, NT5C3A, NT5E, NUDT15, NUDT5, PAICS, PPAT, RRM1, RRM2, SAMHD1, TK1, TXNRD1, TYMP, TYMS, UMPS, UPP1, XDH* |
| R-HSA-5656121 | Translesion synthesis by POLI | 1.65 | 0.021 | *PCNA, RFC1, RFC2, RFC3, RFC4, RFC5, RPA3, RPS27A* |
| R-HSA-9013973 | TICAM1-dependent activation of IRF3/IRF7 | 1.64 | 0.025 | *IKBKE, IRF3, IRF7, RPS27A, TANK, TBK1, TRAF3* |
| R-HSA-5576893 | Phase 2 - plateau phase | -1.92 | 0.025 | *CACNA1D, CACNA1S, CACNA2D1, CACNA2D2, CACNA2D3, CACNB1, CACNB2, CACNB4, KCNE4, KCNQ1* |
| R-HSA-141405 | Inhibition of the proteolytic activity of APC/C required for the onset of anaphase by mitotic spindle checkpoint components | 1.63 | 0.026 | *ANAPC1, ANAPC4, ANAPC5, BUB1B, BUB3, CDC16, CDC20, CDC23, MAD2L1, UBE2C, UBE2D1* |

| | | | | |
|---|---|---|---|---|
| R-HSA-3229121 | Glycogen storage diseases | -1.90 | 0.029 | *EPM2A, GYG2, GYS2, PPP1R3C* |
| R-HSA-3656243 | Defective ST3GAL3 causes MCT12 and EIEE15 | -1.89 | 0.030 | *KERA, OGN, OMD, PRELP* |
| R-HSA-381042 | PERK regulates gene expression | 1.62 | 0.030 | *ATF3, ATF4, ATF6, CCL2, CXCL8, DCP2, DDIT3, DIS3, EIF2S1, EXOSC2, EXOSC3, EXOSC4, EXOSC7, EXOSC8, EXOSC9, HERPUD1, HSPA5, NFYA, PARN* |
| R-HSA-5358565 | Mismatch repair (MMR) directed by MSH2:MSH6 (MutSalpha) | 1.61 | 0.030 | *EXO1, LIG1, MLH1, MSH2, MSH6, PCNA, PMS2, POLD1, POLD2, POLD3, RPA1, RPA3* |
| R-HSA-193775 | Synthesis of bile acids and bile salts via 24-hydroxycholesterol | -1.86 | 0.040 | *AKR1C1, AKR1C2, AKR1C3, CYP27A1, CYP46A1, PTGIS* |
| R-HSA-164952 | The role of Nef in HIV-1 replication and disease pathogenesis | 1.57 | 0.045 | *ATP6V1H, B2M, CD247, CD28, CD4, DOCK2, ELMO1, FYN, HCK, LCK, RAC1* |
| R-HSA-8985947 | Interleukin-9 signaling | 1.57 | 0.047 | *IL2RG, JAK3, STAT1, STAT3* |
| R-HSA-936837 | Ion transport by P-type ATPases | -1.82 | 0.048 | *ATP13A4, ATP13A5, ATP1A2, ATP1B1, ATP1B2, ATP2A1, ATP2B2, ATP8A1, CAMK2B, CUTC, FXYD1, FXYD2, FXYD6, PLN, SLN* |
| R-HSA-166665 | Terminal pathway of complement | -1.85 | 0.049 | *C7, CLU* |

Table S7 - Enriched Reactome pathways in advanced OSCC using GSEA.

| Gene Set | Description | Normalized enrichment score | FDR | Genes |
|---|---|---|---|---|
| R-HSA-69306 | DNA Replication | 2.70 | 0 | *ANAPC1, ANAPC4, ANAPC5, CCNA1, CCNA2, CCNE1, CCNE2, CDC23, CDC45, CDC6, CDC7, CDK2, CDT1, DBF4, DNA2, E2F1, E2F3, FEN1, GINS1, GINS2, GINS3, GINS4, GMNN, LIG1, MCM10, MCM2, MCM3, MCM4, MCM5, MCM6, MCM7, MCM8, ORC1, ORC2, ORC5, ORC6, PCNA, POLA1, POLA2,* |

| | | | | |
|---|---|---|---|---|
| | | | | *POLD1, POLD2, POLD3, POLE, POLE2, POLE3, POLE4, PRIM1, PRIM2, PSMA1, PSMA2, PSMA3, PSMA4, PSMA5, PSMA6, PSMB1, PSMB10, PSMB2, PSMB3, PSMB4, PSMB5, PSMB7, PSMB8, PSMB9, PSMC1, PSMC2, PSMC3, PSMC4, PSMC6, PSMD1, PSMD10, PSMD11, PSMD12, PSMD13, PSMD14, PSMD2, PSMD3, PSMD5, PSMD9, PSME1, PSME2, PSME3, PSME4, RBX1, RFC2, RFC3, RFC4, RFC5, RPA3, SKP2, UBE2C* |
| R-HSA-913531 | Interferon Signaling | 2.68 | 0 | *ABCE1, ADAR, B2M, BST2, CD44, DDX58, EIF2AK2, EIF4A2, EIF4A3, EIF4E2, FCGR1A, FLNA, FLNB, GBP1, GBP2, GBP3, GBP4, GBP5, HERC5, HLA-A, HLA-B, HLA-C, HLA-DPA1, HLA-DQA1, HLA-DQB1, HLA-DRA, HLA-E, HLA-F, HLA-G, ICAM1, IFI27, IFI30, IFI35, IFI6, IFIT1, IFIT2, IFIT3, IFITM1, IFITM3, IFNAR2, IFNG, IFNGR1, IRF1, IRF3, IRF4, IRF6, IRF7, IRF8, IRF9, ISG15, ISG20, JAK2, KPNA2, MT2A, MX1, MX2, NDC1, NEDD4, NUP107, NUP155, NUP160, NUP188, NUP205, NUP210, NUP37, NUP50, NUP54, OAS1, OAS2, OAS3, OASL, PLCG1, PML, PSMB8, PTAFR, PTPN1, PTPN2, RAE1, RSAD2, SAMHD1, SOCS1, SP100, STAT1, STAT2, TRIM14, TRIM21, TRIM22, TRIM38, TRIM5, TRIM6, UBE2L6, USP18, XAF1* |
| R-HSA-68882 | Mitotic Anaphase | 2.61 | 0 | *AHCTF1, ANAPC1, ANAPC4, ANAPC5, ANKLE2, AURKB, BIRC5, BUB1, BUB1B, BUB3, CDC20, CDC23, CDCA5, CDCA8, CENPA, CENPE, CENPF, CENPH, CENPI, CENPK, CENPL, CENPN, CENPO, CENPQ, CENPT, CENPU, CKAP5, DSN1, ERCC6L, ITGB3BP, KIF18A, KIF2A, KIF2C, KNTC1, LMNB1, MAD2L1, MAPRE1, NDC80, NDE1, NUDC, NUF2, NUP107, NUP160, NUP37, NUP85, NUP98, PLK1, PSMA1, PSMA2, PSMA3, PSMA4, PSMA5, PSMA6, PSMB1, PSMB10, PSMB2, PSMB3, PSMB4, PSMB5, PSMB7, PSMB8, PSMB9, PSMC2, PSMC4, PSMC6, PSMD1, PSMD10, PSMD11, PSMD13, PSMD14, PSMD2, PSMD3, PSMD5, PSMD9, PSME1, PSME2, PSME3, PSME4, PTTG1, RAD21, RANGAP1, RCC2, SKA1, SKA2, SMC1A, SMC3, SPC25, SPDL1, TMPO, TUBA1B, TUBA1C, TUBA4A, TUBB3, TUBB4B, UBE2C, VRK1, VRK2, XPO1, ZW10, ZWILCH, ZWINT* |
| R-HSA-68949 | Orc1 removal from chromatin | 2.56 | 0 | *CCNA1, CCNA2, CDC6, CDK2, CDT1, MCM2, MCM3, MCM4, MCM5, MCM6, MCM7, MCM8, ORC1, ORC5, ORC6, PSMA1, PSMA2, PSMA3, PSMA4, PSMA5, PSMA6, PSMB1, PSMB10, PSMB2, PSMB3, PSMB4, PSMB5, PSMB7, PSMB8, PSMB9, PSMC4, PSMC6, PSMD10, PSMD11, PSMD13, PSMD14,* |

| | | | | |
|---|---|---|---|---|
| | | | | *PSMD2, PSMD3, PSMD5, PSMD9, PSME1, PSME2, PSME3, PSME4, SKP2* |
| R-HSA-141424 | Amplification of signal from the kinetochores | 2.50 | 0 | *AHCTF1, AURKB, BIRC5, BUB1, BUB1B, BUB3, CDC20, CDCA8, CENPA, CENPE, CENPF, CENPH, CENPI, CENPK, CENPL, CENPN, CENPO, CENPQ, CENPU, CKAP5, DSN1, ERCC6L, ITGB3BP, KIF18A, KIF2A, KIF2C, KNTC1, MAD2L1, NDC80, NDE1, NUF2, NUP107, NUP160, NUP37, PLK1, RANGAP1, RCC2, SKA1, SKA2, SPC25, SPDL1, XPO1, ZWILCH, ZWINT* |
| R-HSA-176814 | Activation of APC/C and APC/C:Cdc20 mediated degradation of mitotic proteins | 2.50 | 0 | *ANAPC1, ANAPC4, BUB1B, BUB3, CCNA1, CCNA2, CCNB1, CDC20, CDC23, CDK1, MAD2L1, NEK2, PLK1, PSMA1, PSMA2, PSMA3, PSMA4, PSMA5, PSMA6, PSMB1, PSMB10, PSMB2, PSMB3, PSMB4, PSMB5, PSMB7, PSMB8, PSMB9, PSMC2, PSMC4, PSMC6, PSMD10, PSMD11, PSMD13, PSMD14, PSMD2, PSMD3, PSMD5, PSMD9, PSME1, PSME2, PSME3, PSME4, PTTG1, UBE2C* |
| R-HSA-2022090 | Assembly of collagen fibrils and other multimeric structures | 2.42 | 0 | *CD151, COL10A1, COL11A1, COL12A1, COL17A1, COL1A1, COL1A2, COL3A1, COL4A1, COL4A2, COL4A5, COL5A2, COL6A3, COL7A1, CTSB, CTSL, CTSS, CTSV, ITGA6, ITGB4, LAMA3, LAMB3, LAMC2, LOXL2, MMP13, MMP3, MMP7, MMP9, PXDN* |
| R-HSA-5693538 | Homology Directed Repair | 2.41 | 0 | *ATM, ATR, BLM, BRCA1, BRCA2, BRIP1, CCNA1, CCNA2, CDK2, CHEK1, CLSPN, DNA2, EME1, EXO1, FEN1, GEN1, HUS1, MDC1, NBN, PALB2, PARP1, PARP2, PCNA, POLD1, POLD2, POLD3, POLE, POLE2, POLE3, POLE4, POLK, POLQ, PPP4C, RAD1, RAD17, RAD50, RAD51, RAD51AP1, RAD51B, RAD51C, RAD9A, RBBP8, RFC1, RFC2, RFC3, RFC4, RFC5, RMI1, RMI2, RPA3, RTEL1, TIMELESS, TIPIN, TOPBP1, UBE2I, UBE2V2, UIMC1, XRCC2, XRCC3* |
| R-HSA-1236975 | Antigen processing-Cross presentation | 2.41 | 0 | *B2M, CALR, CD14, CHUK, CTSL, CTSS, CTSV, CYBA, CYBB, FCGR1A, HLA-A, HLA-B, HLA-C, HLA-E, HLA-F, HLA-G, ITGAV, LY96, NCF1, NCF2, NCF4, PDIA3, PSMA1, PSMA2, PSMA3, PSMA4, PSMA5, PSMA6, PSMB1, PSMB10, PSMB2, PSMB3, PSMB4, PSMB5, PSMB7, PSMB8, PSMB9, PSMC2, PSMC4, PSMC6, PSMD1, PSMD10, PSMD11, PSMD13, PSMD14, PSMD2, PSMD3, PSMD5, PSMD9, PSME1, PSME2, PSME3, PSME4, SEC61A1, SEC61A2, SEC61B, SEC61G, STX4, TAP1, TAP2, TAPBP, TLR1, TLR2, TLR6* |
| R-HSA-68962 | Activation of the pre-replicative | 2.40 | 0 | *CDC45, CDC6, CDC7, CDK2, CDT1, DBF4, GMNN, MCM10, MCM2, MCM3,* |

| | | | | |
|---|---|---|---|---|
| | complex | | | *MCM4, MCM5, MCM6, MCM7, MCM8, ORC1, ORC2, ORC5, ORC6, POLA1, POLA2, POLE, POLE2, POLE3, POLE4, PRIM1, PRIM2, RPA3* |
| R-HSA-162906 | HIV Infection | 2.39 | 0 | *AAAS, AP1S1, AP1S3, AP2B1, AP2M1, AP2S1, APOBEC3G, ATP6V1H, B2M, CCNK, CCNT2, CD247, CD28, CD4, CD8B, CDK9, CHMP4A, CHMP5, CXCR4, DOCK2, ELL, ERCC2, ERCC3, FEN1, FURIN, FYN, GTF2A2, GTF2B, GTF2E1, GTF2E2, GTF2F2, GTF2H3, GTF2H4, HCK, KPNA1, KPNB1, LCK, LIG1, LIG4, MNAT1, NCBP1, NCBP2, NDC1, NEDD4L, NELFCD, NMT1, NUP107, NUP153, NUP155, NUP160, NUP188, NUP205, NUP210, NUP214, NUP35, NUP37, NUP43, NUP50, NUP54, NUP85, NUP88, NUP93, NUP98, PAK2, POLR2D, POLR2G, POLR2H, POLR2K, POLR2L, PPIA, PSMA1, PSMA2, PSMA3, PSMA4, PSMA5, PSMA6, PSMB1, PSMB10, PSMB2, PSMB3, PSMB4, PSMB5, PSMB7, PSMB8, PSMB9, PSMC1, PSMC2, PSMC3, PSMC4, PSMC6, PSMD1, PSMD10, PSMD11, PSMD12, PSMD13, PSMD14, PSMD2, PSMD3, PSMD5, PSMD6, PSMD7, PSMD8, PSMD9, PSME1, PSME2, PSME3, PSME4, RAC1, RAE1, RAN, RANBP1, RANGAP1, RBX1, RCC1, RNGTT, RNMT, SLC25A5, SSRP1, SUPT16H, TAF11, TAF12, TAF13, TAF2, TAF4B, TBP, TCEA1, TPR, UBAP1, XPO1, XRCC4, XRCC6* |
| R-HSA-453274 | Mitotic G2-G2/M phases | 2.34 | 0 | *AJUBA, AURKA, BORA, CCNA1, CCNA2, CCNB1, CCNB2, CDC25A, CDC25B, CDC25C, CDK1, CDK2, CDKN1A, CENPF, CENPJ, CEP135, CEP152, CEP192, CEP290, CEP78, CKAP5, CNTRL, CSNK1E, E2F1, E2F3, FOXM1, HAUS1, HAUS2, HAUS3, HAUS6, HMMR, HSP90AA1, LIN9, MAPRE1, MNAT1, MYBL2, NDE1, NEDD1, NEK2, NME7, ODF2, PHLDA1, PKMYT1, PLK1, PLK4, PSMA1, PSMA2, PSMA3, PSMA4, PSMA5, PSMA6, PSMB1, PSMB10, PSMB2, PSMB3, PSMB4, PSMB5, PSMB7, PSMB8, PSMB9, PSMC1, PSMC2, PSMC3, PSMC4, PSMC6, PSMD1, PSMD10, PSMD11, PSMD12, PSMD13, PSMD14, PSMD2, PSMD3, PSMD5, PSMD9, PSME1, PSME2, PSME3, PSME4, RAB8A, RBX1, TPX2, TUBA1B, TUBA1C, TUBA4A, TUBB, TUBB3, TUBB4B, TUBG1, TUBGCP3, TUBGCP5, XPO1, YWHAE* |
| R-HSA-1168372 | Downstream signaling events of B Cell Receptor (BCR) | 2.33 | 0 | *CALM1, CALM3, CARD11, CHUK, CUL1, FKBP1A, HRAS, KRAS, MALT1, MAP3K7, NFKB1, NFKBIA, NFKBIE, NRAS, PPIA, PRKCB, PSMA1, PSMA2, PSMA3, PSMA4, PSMA5, PSMA6, PSMB1, PSMB10, PSMB2, PSMB3, PSMB4, PSMB5, PSMB7, PSMB8, PSMB9, PSMC1, PSMC2, PSMC3, PSMC4, PSMC6,* |

| | | | | |
|---|---|---|---|---|
| | | | | *PSMD1, PSMD10, PSMD11, PSMD12, PSMD13, PSMD14, PSMD2, PSMD3, PSMD5, PSMD9, PSME1, PSME2, PSME3, PSME4, RASGRP1, REL* |
| R-HSA-1169410 | Antiviral mechanism by IFN-stimulated genes | 2.32 | 0 | *ABCE1, DDX58, EIF2AK2, EIF4A2, EIF4A3, EIF4E2, FLNA, FLNB, HERC5, IFIT1, IRF3, ISG15, KPNA1, KPNA2, MX1, MX2, NDC1, NEDD4, NUP107, NUP153, NUP155, NUP160, NUP188, NUP205, NUP210, NUP214, NUP37, NUP50, NUP54, NUP85, NUP93, NUP98, OAS1, OAS2, OAS3, OASL, PLCG1, RAE1, STAT1, TRIM25, UBA7, UBE2L6, USP18* |
| R-HSA-6783783 | Interleukin-10 signaling | 2.30 | 0 | *CCL20, CCL3, CCL4, CCL5, CCR1, CD80, CD86, CSF2, CXCL1, CXCL10, CXCL2, CXCL8, ICAM1, IL10RA, IL1A, IL1B, PTAFR, TIMP1* |
| R-HSA-4641258 | Degradation of DVL | 2.28 | 0 | *PSMA1, PSMA2, PSMA3, PSMA4, PSMA5, PSMA6, PSMB1, PSMB10, PSMB2, PSMB3, PSMB4, PSMB5, PSMB7, PSMB8, PSMB9, PSMC1, PSMC2, PSMC3, PSMC4, PSMC6, PSMD1, PSMD10, PSMD11, PSMD12, PSMD13, PSMD14, PSMD2, PSMD3, PSMD5, PSMD9, PSME1, PSME2, PSME3, PSME4, RBX1* |
| R-HSA-69205 | G1/S-Specific Transcription | 2.22 | 0 | *CCNA1, CCNE1, CDC45, CDC6, CDK1, CDT1, DHFR, E2F1, E2F4, E2F5, FBXO5, HDAC1, LIN9, ORC1, PCNA, POLA1, RBL1, RRM2, TK1, TYMS* |
| R-HSA-195258 | RHO GTPase Effectors | 2.20 | 0 | *ACTR2, ACTR3, AHCTF1, ARPC1A, ARPC1B, ARPC2, AURKB, BIRC5, BUB1, BUB1B, BUB3, CALM1, CALM3, CDC20, CDC25C, CDCA8, CENPA, CENPE, CENPF, CENPH, CENPI, CENPK, CENPL, CENPM, CENPN, CENPO, CENPQ, CENPT, CENPU, CFL1, CIT, CKAP5, CTTN, CYBA, CYBB, CYFIP2, DIAPH1, DIAPH3, DSN1, ERCC6L, EVL, FLNA, FMNL2, FMNL3, GOPC, GRB2, IQGAP3, ITGB1, ITGB3BP, KDM1A, KIF14, KIF18A, KIF2A, KIF2C, KLC2, KNTC1, KTN1, LIMK1, LIMK2, MAD2L1, MAPRE1, MYH10, MYH9, MYL6, NCF1, NCF2, NCF4, NCKAP1L, NDC80, NDE1, NDEL1, NUDC, NUF2, NUP107, NUP160, NUP37, NUP43, NUP85, NUP98, PAK1, PAK2, PFN1, PFN2, PIK3R4, PLK1, PPP2R5B, PRC1, PTK2, RAC2, RANGAP1, RCC2, RHOC, RHOD, RHOG, RHPN2, ROCK1, RTKN, SFN, SKA1, SKA2, SPC25, SPDL1, SRC, SRGAP2, TUBA1B, TUBA1C, TUBA4A, TUBB3, TUBB4B, WAS, WIPF1, XPO1, YWHAE, YWHAH, YWHAQ, YWHAZ, ZW10, ZWILCH, ZWINT* |
| R-HSA-194441 | Metabolism of non-coding RNA | 2.18 | 0 | *AAAS, CLNS1A, DDX20, GEMIN2, GEMIN4, GEMIN6, GEMIN7, NCBP1, NCBP2, NDC1, NUP107, NUP153, NUP155, NUP160, NUP188, NUP205, NUP210, NUP214, NUP35, NUP37, NUP43, NUP50, NUP54, NUP85, NUP88,* |

| | | | | |
|---|---|---|---|---|
| | | | | *NUP93, NUP98, PRMT5, RAE1, SMN1, SNRPB, SNRPD1, SNRPD2, SNRPE, SNRPF, SNRPG, TGS1, TPR, WDR77* |
| R-HSA-72172 | mRNA Splicing | 2.17 | 0 | *ALYREF, AQR, BUD31, CCAR1, CLP1, CPSF1, CPSF2, CPSF3, CSTF1, CSTF2, CSTF3, CTNNBL1, CWC15, CWC27, DDX23, DDX46, DHX15, DHX16, DHX38, DHX9, DNAJC8, EFTUD2, EIF4A3, ELAVL2, FIP1L1, FUS, GTF2F2, HNRNPA1, HNRNPA2B1, HNRNPA3, HNRNPC, HNRNPD, HNRNPF, HNRNPH1, HNRNPK, HNRNPL, HNRNPM, HNRNPR, HNRNPU, HSPA8, ISY1, LSM2, LSM3, LSM5, LSM6, LSM7, LSM8, MAGOH, MAGOHB, NCBP1, NCBP2, PAPOLA, PHF5A, PLRG1, POLR2A, POLR2D, POLR2G, POLR2H, POLR2J, POLR2K, POLR2L, PPIH, PPIL1, PPWD1, PRCC, PRPF19, PRPF3, PRPF31, PRPF4, PRPF40A, PTBP1, PUF60, RBM22, RBMX, RNPS1, SART1, SF3A3, SF3B2, SF3B3, SF3B4, SF3B6, SMNDC1, SNRNP40, SNRNP48, SNRPA, SNRPA1, SNRPB, SNRPB2, SNRPC, SNRPD1, SNRPD2, SNRPE, SNRPF, SNRPG, SNW1, SRRT, SRSF1, SRSF10, SRSF11, SRSF2, SRSF3, SRSF7, SRSF9, SYMPK, TFIP11, TRA2B, U2AF2, U2SURP, UPF3B, USP39, WBP11, WDR33, ZRSR2* |
| R-HSA-180786 | Extension of Telomeres | 2.16 | 0 | *DKC1, DNA2, FEN1, LIG1, PCNA, POLA1, POLA2, POLD1, POLD3, POLE, POLE2, POLE3, POLE4, PRIM1, PRIM2, RFC2, RFC3, RFC4, RFC5, RPA3, RUVBL1, WRAP53* |
| R-HSA-72202 | Transport of Mature Transcript to Cytoplasm | 2.16 | 0 | *AAAS, ALYREF, CPSF1, CPSF2, CPSF3, DDX39A, EIF4A3, FIP1L1, GLE1, MAGOH, MAGOHB, NCBP1, NCBP2, NDC1, NUP107, NUP153, NUP155, NUP160, NUP188, NUP205, NUP210, NUP214, NUP35, NUP37, NUP43, NUP50, NUP54, NUP85, NUP88, NUP93, NUP98, RAE1, RNPS1, SARNP, SLBP, SRSF1, SRSF2, SRSF3, SRSF7, SRSF9, SYMPK, THOC1, THOC2, THOC3, THOC6, TPR, U2AF2, UPF3B, ZC3H11A* |
| R-HSA-211945 | Phase I - Functionalization of compounds | -2.37 | 0 | *AADAC, ACSS2, ADH1A, ADH1B, ADH1C, ADH4, ADH7, ALDH1A1, ALDH2, ALDH3A1, AOC3, CES1, CES2, CYP11A1, CYP1B1, CYP27A1, CYP2C18, CYP2C19, CYP2F1, CYP2J2, CYP2R1, CYP2U1, CYP3A4, CYP3A5, CYP4B1, CYP4F12, CYP4F22, CYP4F3, CYP7B1, EPHX1, FMO2, MAOB, MARC1, MARC2, NCOA1, PTGIS* |
| R-HSA-397014 | Muscle contraction | -2.71 | 0 | *ABCC9, ACTA1, ACTA2, ACTC1, ACTG2, ACTN2, ANXA6, ATP1A2, ATP1B1,* |

| | | | | |
|---|---|---|---|---|
| | | | | *ATP1B2, ATP2A1, ATP2B2, CACNA1D, CACNA1S, CACNA2D1, CACNA2D3, CACNB1, CACNB2, CACNB4, CACNG1, CACNG6, CAMK2B, CASQ1, CASQ2, CAV3, CLIC2, CORIN, DES, DMD, DMPK, FGF12, FGF14, FXYD1, FXYD6, HIPK2, ITPR2, KAT2B, KCNIP2, KCNIP3, KCNJ2, LMOD1, MYBPC1, MYBPC2, MYH11, MYH3, MYH6, MYH8, MYL1, MYL2, MYL3, MYL6B, MYL9, MYLK, MYLPF, NEB, NOS1, NPR1, NPR2, PLN, PRKACA, RYR1, RYR3, SCN11A, SCN1B, SCN2B, SCN4A, SCN4B, SCN5A, SCN7A, SLC8A3, SLN, SORBS1, SORBS3, STIM1, TCAP, TMOD1, TMOD4, TNNC1, TNNC2, TNNI1, TNNI2, TNNT1, TNNT3, TPM2, TPM3, TRDN, TRPC1, TTN, WWTR1* |
| R-HSA-606279 | Deposition of new CENPA-containing nucleosomes at the centromere | 2.09 | 0 | *CENPA, CENPH, CENPI, CENPK, CENPL, CENPM, CENPN, CENPO, CENPQ, CENPT, CENPU, CENPW, HJURP, ITGB3BP, MIS18A, MIS18BP1, OIP5, RUVBL1, SMARCA5* |
| R-HSA-1538133 | G0 and Early G1 | 2.10 | 0 | *CCNA1, CCNA2, CCNE1, CCNE2, CDC25A, CDC6, CDK1, CDK2, E2F1, E2F4, E2F5, HDAC1, LIN9, MYBL2, PCNA, RBL1, TOP2A* |
| R-HSA-5693537 | Resolution of D-Loop Structures | 2.12 | 0 | *BLM, BRCA1, BRCA2, BRIP1, DNA2, EME1, EXO1, GEN1, NBN, PALB2, RAD50, RAD51, RAD51AP1, RBBP8, RMI1, RTEL1, XRCC2* |
| R-HSA-1236977 | Endosomal/Vacuolar pathway | 2.06 | 0 | *B2M, CTSL, CTSS, CTSV, HLA-A, HLA-B, HLA-C, HLA-E, HLA-F, HLA-G* |
| R-HSA-5685938 | HDR through Single Strand Annealing (SSA) | 2.04 | 0 | *ATM, ATR, BLM, BRCA1, BRIP1, DNA2, EXO1, HUS1, NBN, RAD1, RAD17, RAD50, RAD51, RBBP8, RFC2, RFC3, RFC4, RFC5, RMI1, RMI2, RPA3, TOPBP1* |
| R-HSA-210991 | Basigin interactions | 2.03 | 0 | *ATP1B3, CAV1, ITGA3, ITGA6, MMP1, SLC16A1, SLC3A2, SLC7A11, SLC7A5, SLC7A7, SLC7A8* |
| R-HSA-3000171 | Non-integrin membrane-ECM interactions | 2.03 | 0 | *ACTN1, AGRN, CASK, COL10A1, COL11A1, COL1A1, COL1A2, COL3A1, COL4A1, COL4A2, COL4A5, COL5A1, COL5A2, FN1, ITGA2, ITGA6, ITGAV, ITGB1, ITGB4, LAMA1, LAMA3, LAMB1, LAMB3, LAMC2, SDC4, THBS1, TNC* |
| R-HSA-110373 | Resolution of AP sites via the multiple-nucleotide patch replacement pathway | 2.02 | 0 | *ADPRHL2, FEN1, LIG1, PARG, PARP1, PARP2, PCNA, POLD1, POLD2, POLD3, POLE, POLE2, POLE3, POLE4, RFC1, RFC2, RFC3, RFC4, RFC5, RPA3* |
| R-HSA-450408 | AUF1 (hnRNP D0) binds and | 2.01 | 0 | *PSMA1, PSMA2, PSMA3, PSMA4, PSMA5, PSMA6, PSMB1, PSMB10, PSMB2,* |

| | | | | |
|---|---|---|---|---|
| | destabilizes mRNA | | | *PSMB3, PSMB4, PSMB5, PSMB7, PSMB8, PSMB9, PSMC2, PSMC4, PSMC6, PSMD10, PSMD11, PSMD13, PSMD14, PSMD2, PSMD3, PSMD5, PSMD9, PSME1, PSME2, PSME3, PSME4* |
| R-HSA-6783310 | Fanconi Anemia Pathway | 2.02 | 0 | *ATR, DCLRE1A, DCLRE1B, EME1, FANCA, FANCB, FANCD2, FANCE, FANCG, FANCI, FANCL, FANCM, RPA3, UBE2T, USP1* |
| R-HSA-6811434 | COPI-dependent Golgi-to-ER retrograde traffic | 2.00 | 0 | *ARF3, ARF4, ARFGAP1, CENPE, COPB1, COPB2, COPG1, KDELR2, KIF11, KIF15, KIF16B, KIF18A, KIF18B, KIF20A, KIF20B, KIF21B, KIF22, KIF23, KIF26B, KIF2A, KIF2C, KIF3A, KIF3C, KIF4A, KIFC1, KLC2, NAPB, NAPG, RACGAP1, SURF4, TMED9, TUBA1B, TUBA1C, TUBA4A, TUBB3, TUBB4B, ZW10* |
| R-HSA-8875878 | MET promotes cell motility | 1.99 | 0 | *COL11A1, COL1A1, COL1A2, COL27A1, COL3A1, COL5A1, COL5A2, FN1, ITGA2, ITGA3, ITGB1, LAMA1, LAMA3, LAMB1, LAMB3, LAMC2, MET, RAP1B, TNS4* |
| R-HSA-110313 | Translesion synthesis by Y family DNA polymerases bypasses lesions on DNA template | 1.98 | 0 | *ISG15, MAD2L2, NPLOC4, PCNA, POLD1, POLD2, POLD3, POLE, POLE2, POLE3, POLE4, RFC1, RFC2, RFC3, RFC4, RFC5, RPA3, TRIM25, UBA7, UBE2L6, USP10, USP43, VCP* |
| R-HSA-109581 | Apoptosis | 1.98 | 0 | *APAF1, BAK1, BAX, BCAP31, BCL2L1, BCL2L11, BID, BIRC2, CASP3, CASP6, CASP7, CASP8, CD14, CLSPN, DAPK1, DAPK3, DNM1L, DSG2, DSG3, E2F1, FADD, FASLG, GZMB, HMGB2, KPNA1, LMNB1, LY96, NMT1, PAK2, PLEC, PMAIP1, PSMA1, PSMA2, PSMA3, PSMA4, PSMA5, PSMA6, PSMB1, PSMB10, PSMB2, PSMB3, PSMB4, PSMB5, PSMB7, PSMB8, PSMB9, PSMC1, PSMC2, PSMC3, PSMC4, PSMC6, PSMD1, PSMD10, PSMD11, PSMD12, PSMD13, PSMD14, PSMD2, PSMD3, PSMD5, PSMD9, PSME1, PSME2, PSME3, PSME4, PTK2, RIPK1, SFN, STAT3, TFDP1, TNFRSF10A, TNFRSF10B, TNFSF10, TP63, TP73, YWHAE, YWHAH, YWHAZ* |
| R-HSA-69091 | Polymerase switching | 1.96 | 0.001 | *PCNA, POLA1, POLA2, POLD1, POLD3, PRIM1, PRIM2, RFC2, RFC3, RFC4, RFC5* |
| R-HSA-162658 | Golgi Cisternae Pericentriolar Stack Reorganization | 1.96 | 0.001 | *CCNB1, CCNB2, CDK1, PLK1* |
| R-HSA-912694 | Regulation of IFNA signaling | 1.94 | 0.001 | *IFNAR2, PTPN1, SOCS1, STAT1, STAT2, USP18* |

| | | | | |
|---|---|---|---|---|
| R-HSA-209822 | Glycoprotein hormones | 1.94 | 0.001 | *INHBA* |
| R-HSA-170822 | Regulation of Glucokinase by Glucokinase Regulatory Protein | 1.92 | 0.001 | *AAAS, NDC1, NUP107, NUP153, NUP155, NUP160, NUP188, NUP205, NUP210, NUP214, NUP35, NUP37, NUP43, NUP50, NUP54, NUP85, NUP88, NUP93, NUP98, RAE1, TPR* |
| R-HSA-8868773 | rRNA processing in the nucleus and cytosol | 1.89 | 0.001 | *BMS1, BOP1, BYSL, C1D, CSNK1E, DCAF13, DDX21, DDX47, DDX52, DIS3, DKC1, EBNA1BP2, EMG1, ERI1, EXOSC10, EXOSC2, EXOSC3, EXOSC4, EXOSC8, EXOSC9, FCF1, FTSJ3, GNL3, HEATR1, ISG20L2, MPHOSPH6, NAT10, NCL, NIP7, NOB1, NOL11, NOL6, NOL9, NOP10, NOP14, NOP2, NOP56, NOP58, PDCD11, PES1, PNO1, RBM28, RCL1, RIOK1, RIOK2, RPL22L1, RPL23, RPL34, RPL35, RPL39L, RPP30, RPP40, RPS20, RPS21, RPS24, RPS4Y1, RPS7, RRP36, RRP9, SENP3, TEX10, TRMT112, TSR1, UTP14A, UTP15, UTP20, UTP6, WDR12, WDR3, WDR36, WDR43, WDR75, XRN2* |
| R-HSA-9020591 | Interleukin-12 signaling | 1.88 | 0.002 | *ANXA2, BOLA2, CAPZA1, CFL1, CNN2, GSTO1, HNRNPA2B1, HNRNPF, IFNG, IL12RB1, IL12RB2, JAK2, LCP1, LMNB1, MIF, MSN, MTAP, PAK2, PSME2, RALA, RAP1B, SNRPA1, SOD2, STAT4* |
| R-HSA-69183 | Processive synthesis on the lagging strand | 1.88 | 0.002 | *DNA2, FEN1, LIG1, PCNA, POLA1, POLA2, POLD1, POLD2, POLD3, PRIM1, PRIM2, RPA3* |
| R-HSA-202430 | Translocation of ZAP-70 to Immunological synapse | 1.88 | 0.002 | *CD247, CD3D, CD3E, CD3G, CD4, HLA-DPA1, HLA-DQA1, HLA-DQB1, HLA-DRA, LCK, PTPN22, ZAP70* |
| R-HSA-168928 | DDX58/IFIH1-mediated induction of interferon-alpha/beta | 1.87 | 0.002 | *APP, CASP10, CASP8, CHUK, CYLD, DDX58, DHX58, FADD, HERC5, IFIH1, IKBKE, IRF3, IRF7, ISG15, NFKB1, NFKB2, NFKBIA, NLRC5, RIPK1, S100A12, TANK, TBK1, TNFAIP3, TRAF3, TRIM25, UBA7, UBE2L6* |
| R-HSA-109688 | Cleavage of Growing Transcript in the Termination Region | 1.87 | 0.002 | *ALYREF, CHTOP, CLP1, CPSF1, CPSF2, CPSF3, CSTF1, CSTF2, CSTF3, DDX39A, DHX38, EIF4A3, FIP1L1, MAGOH, MAGOHB, NCBP1, NCBP2, PAPOLA, RNPS1, SARNP, SLBP, SNRPB, SNRPE, SNRPF, SNRPG, SRSF1, SRSF11, SRSF2, SRSF3, SRSF7, SRSF9, SYMPK, THOC1, THOC2, THOC3, THOC6, U2AF2, UPF3B, WDR33, ZC3H11A* |
| R-HSA-6781827 | Transcription-Coupled Nucleotide Excision Repair | 1.87 | 0.002 | *AQR, COPS2, COPS5, COPS7B, ELL, ERCC2, ERCC3, ERCC6, GTF2H1, GTF2H3, GTF2H4, HMGN1, ISY1, LIG1, MNAT1, PCNA, POLD1, POLD2,* |

| | | | | |
|---|---|---|---|---|
| | (TC-NER) | | | *POLD3, POLE, POLE2, POLE3, POLE4, POLK, POLR2A, POLR2D, POLR2G, POLR2H, POLR2J, POLR2K, POLR2L, PRPF19, RBX1, RFC1, RFC2, RFC3, RFC4, RFC5, RPA3, TCEA1* |
| R-HSA-2424491 | DAP12 signaling | 1.85 | 0.002 | *B2M, FYN, GRAP2, GRB2, HLA-E, HRAS, KLRC2, KLRD1, KRAS, LAT, LCK, LCP2, NRAS, PLCG1, SHC1, TREM2, TYROBP, VAV2* |
| R-HSA-168643 | Nucleotide-binding domain, leucine rich repeat containing receptor (NLR) signaling pathways | 1.82 | 0.004 | *AIM2, APP, BCL2L1, BIRC2, BIRC3, CASP1, CASP2, CASP4, CASP8, CHUK, CYLD, HSP90AB1, IRAK1, IRAK2, ITCH, MAP3K7, MEFV, NFKB1, NFKB2, NLRP1, NLRP3, NOD1, NOD2, PANX1, PSTPIP1, PYCARD, RIPK2, TAB2, TNFAIP3, TXN* |
| R-HSA-73863 | RNA Polymerase I Transcription Termination | 1.82 | 0.004 | *CD3EAP, ERCC2, ERCC3, GTF2H1, GTF2H3, GTF2H4, MNAT1, POLR1A, POLR1E, POLR2H, POLR2K, POLR2L, TAF1A, TAF1B, TAF1D, TBP, TTF1, TWISTNB* |
| R-HSA-114604 | GPVI-mediated activation cascade | 1.82 | 0.004 | *COL1A1, COL1A2, FCER1G, FYN, LAT, LCK, LCP2, LYN, PDPN, PIK3CG, PIK3R5, RAC2, RHOG, VAV1, VAV2* |
| R-HSA-75205 | Dissolution of Fibrin Clot | 1.81 | 0.004 | *PLAT, PLAU, PLAUR, SERPINE1, SERPINE2* |
| R-HSA-164952 | The role of Nef in HIV-1 replication and disease pathogenesis | 1.80 | 0.005 | *AP1S1, AP1S3, AP2B1, AP2M1, ATP6V1H, B2M, CD247, CD28, CD4, CD8B, DOCK2, FYN, HCK, LCK, PAK2* |
| R-HSA-5358508 | Mismatch Repair | 1.80 | 0.005 | *EXO1, LIG1, MSH2, MSH6, PCNA, PMS2, POLD1, POLD2, POLD3, RPA3* |
| R-HSA-3229121 | Glycogen storage diseases | -1.96 | 0.005 | *EPM2A, GBE1, GYG2, GYS2, PPP1R3C* |
| R-HSA-6804114 | TP53 Regulates Transcription of Genes Involved in G2 Cell Cycle Arrest | 1.78 | 0.006 | *AURKA, BAX, CCNB1, CDC25C, CDK1, E2F4, PCNA, RBL1, SFN* |
| R-HSA-1500620 | Meiosis | 1.78 | 0.006 | *ATR, BLM, BRCA1, BRCA2, CDK2, CDK4, LMNB1, MND1, MSH5, NBN, POT1, PSMC3IP, RAD21, RAD50, RAD51, RBBP8, REC8, RPA3, SMC1A, SMC3, SYNE2* |
| R-HSA-167152 | Formation of HIV elongation | 1.78 | 0.006 | *CCNK, CCNT2, CDK9, ELL, ERCC2, ERCC3, GTF2F2, GTF2H1, GTF2H3,* |

| | | | | |
|---|---|---|---|---|
| | complex in the absence of HIV Tat | | | *GTF2H4, MNAT1, NCBP1, NCBP2, NELFCD, POLR2A, POLR2D, POLR2G, POLR2H, POLR2J, POLR2K, POLR2L, SSRP1, SUPT16H, TCEA1* |
| R-HSA-8852135 | Protein ubiquitination | 1.76 | 0.008 | *CTR9, DERL1, HLA-A, LEO1, OTULIN, PCNA, PEX13, PEX14, PEX2, PRKDC, RAD18, RNF152, RPS27A, RTF1, UBA1, UBA6, UBE2A, UBE2C, UBE2D1, UBE2D2, UBE2G1, UBE2H, UBE2K, UBE2L3, UBE2Q2, UBE2R2, UBE2S, UBE2T, UBE2V2, UBE2Z, UCHL3, USP9X, VCP* |
| R-HSA-8978868 | Fatty acid metabolism | -1.92 | 0.010 | *ACAA2, ACACB, ACADL, ACADM, ACADS, ACADVL, ACOT2, ACOX1, ACOX2, ACOX3, ACSL1, ACSM3, AKR1C3, ALDH3A2, ALOX12, CRAT, CYP1B1, CYP2C19, CYP2J2, CYP2U1, CYP4B1, CYP4F22, CYP4F3, ECHS1, ELOVL4, ELOVL6, EPHX2, GPX2, GPX4, HADH, HADHB, HPGD, HPGDS, HSD17B8, MLYCD, NUDT7, PHYH, PON3, PRKAA2, PTGDS, PTGIS, PTGR1, SCP2, SLC25A20, TECR, TECRL, THRSP* |
| R-HSA-8953750 | Transcriptional Regulation by E2F6 | 1.74 | 0.010 | *APAF1, BRCA1, CBX3, CDC7, CHEK1, E2F1, EED, EPC1, EZH2, PCGF6, RAD51, RBBP8, RRM2, TFDP1* |
| R-HSA-390471 | Association of TriC/CCT with target proteins during biosynthesis | 1.72 | 0.012 | *CCNE1, CCNE2, CCT2, CCT3, CCT4, CCT5, CCT6A, CCT7, FBXO6, FBXW10, FBXW2, FKBP9, GBA, KIFC3, NOP56, SKIV2L, SPHK1, STAT3, TCP1, WRAP53, XRN2* |
| R-HSA-1834949 | Cytosolic sensors of pathogen-associated DNA | 1.72 | 0.012 | *CHUK, DDX41, DHX36, DHX9, IFI16, IRF3, IRF7, MYD88, NFKB1, NFKB2, NFKBIA, NFKBIB, NLRC3, POLR2H, POLR2K, POLR2L, POLR3A, POLR3F, POLR3G, POLR3K, PRKDC, RIPK1, TBK1, TRIM21, TRIM32, XRCC6, ZBP1* |
| R-HSA-68952 | DNA replication initiation | 1.72 | 0.013 | *POLA1, POLA2, POLE, POLE2, POLE3, POLE4, PRIM1, PRIM2* |
| R-HSA-918233 | TRAF3-dependent IRF activation pathway | 1.71 | 0.013 | *DDX58, IFIH1, IKBKE, IRF3, IRF7, TBK1, TRAF3, TRIM25* |
| R-HSA-8948216 | Collagen chain trimerization | 1.71 | 0.014 | *COL10A1, COL11A1, COL12A1, COL17A1, COL1A1, COL1A2, COL27A1, COL3A1, COL4A1, COL4A2, COL4A5, COL4A6, COL5A1, COL5A2, COL6A1, COL6A3, COL7A1, COL8A1* |
| R-HSA-380259 | Loss of Nlp from mitotic centrosomes | 1.70 | 0.015 | *CDK1, CENPJ, CEP135, CEP152, CEP192, CEP290, CEP57, CEP72, CEP78, CKAP5, CNTRL, CSNK1E, DYNLL1, FGFR1OP, HAUS1, HAUS2, HAUS3, HAUS6, HSP90AA1, MAPRE1, NDE1, NEDD1, NEK2, ODF2, PLK1, PLK4, SDCCAG8, SFI1, SSNA1, TUBA4A, TUBB, TUBB4B, TUBG1, YWHAE* |

| | | | | |
|---|---|---|---|---|
| R-HSA-72086 | mRNA Capping | 1.70 | 0.015 | *ERCC2, ERCC3, GTF2F2, GTF2H1, GTF2H3, GTF2H4, MNAT1, NCBP1, NCBP2, POLR2A, POLR2D, POLR2G, POLR2H, POLR2J, POLR2K, POLR2L, RNGTT, RNMT* |
| R-HSA-936837 | Ion transport by P-type ATPases | -1.88 | 0.016 | *ATP12A, ATP13A4, ATP13A5, ATP1A2, ATP1B1, ATP1B2, ATP2A1, ATP2B2, ATP7A, ATP8A1, ATP9A, CAMK2B, CUTC, FXYD1, FXYD6, PLN, SLN* |
| R-HSA-5576893 | Phase 2 - plateau phase | -1.86 | 0.016 | *CACNA1D, CACNA1S, CACNA2D1, CACNA2D2, CACNA2D3, CACNB1, CACNB2, CACNB4, CACNG1, CACNG6* |
| R-HSA-418457 | cGMP effects | -1.86 | 0.016 | *ITPR1, KCNMA1, KCNMB4, PDE11A, PDE1A, PDE1B, PDE2A, PDE3A, PDE3B, PDE5A, PDE9A, PRKG1, PRKG2* |
| R-HSA-113507 | E2F-enabled inhibition of pre-replication complex formation | 1.70 | 0.016 | *CCNB1, CDK1, MCM8, ORC1, ORC2, ORC5, ORC6* |
| R-HSA-5250913 | Positive epigenetic regulation of rRNA expression | 1.70 | 0.016 | *ACTB, BAZ1B, CBX3, CD3EAP, CHD3, CHD4, DDX21, DEK, ERCC6, GATAD2A, GSK3B, HDAC1, HDAC2, KAT2A, MTA2, MYBBP1A, POLR1A, POLR1C, POLR1E, POLR2H, POLR2K, POLR2L, SMARCA5, TAF1A, TAF1B, TAF1D, TBP, TTF1, TWISTNB* |
| R-HSA-193368 | Synthesis of bile acids and bile salts via 7alpha-hydroxycholesterol | -1.87 | 0.016 | *ACOX2, AKR1C1, AKR1C2, AKR1C3, AMACR, BAAT, CYP27A1, CYP7B1, NCOA1, PTGIS, SCP2, SLC27A2* |
| R-HSA-2559583 | Cellular Senescence | 1.69 | 0.017 | *ANAPC1, ANAPC10, ANAPC4, ANAPC5, ATM, CCNA1, CCNA2, CCNE1, CCNE2, CDC23, CDC27, CDK2, CDK4, CDK6, CDKN1A, CDKN2A, CEBPB, CXCL8, E2F1, E2F3, EED, ETS1, ETS2, EZH2, HMGA2, ID1, IGFBP7, IL1A, LMNB1, MAP2K4, MAP3K5, MAP4K4, MAPK9, MDM2, MINK1, MOV10, NBN, NFKB1, POT1, RAD50, RB1, RNF2, RPS6KA1, STAT3, SUZ12, TFDP1, TINF2, TXN, UBE2C, UBE2D1, UBN1* |
| R-HSA-450604 | KSRP (KHSRP) binds and destabilizes mRNA | 1.69 | 0.017 | *DCP2, DIS3, EXOSC2, EXOSC3, EXOSC4, EXOSC8, EXOSC9, PARN, YWHAZ* |
| R-HSA-5633007 | Regulation of TP53 Activity | 1.68 | 0.019 | *ATM, ATR, AURKA, AURKB, BLM, BRCA1, BRIP1, CCNA1, CCNA2, CDK1, CDK2, CDK5, CDKN2A, CHEK1, CSNK2A2, DAXX, DNA2, EXO1, HDAC1, HUS1, MDM2, NBN, NOC2L, PIP4K2A, PIP4K2C, PML, PRDM1, PRMT5,* |

| | | | | |
|---|---|---|---|---|
| | | | | *RAD1, RAD17, RAD50, RBBP8, RFC2, RFC3, RFC4, RFC5, RICTOR, RMI1, RMI2, RNF34, RPA3, SMYD2, SSRP1, SUPT16H, TAF11, TAF12, TAF13, TAF2, TAF4B, TBP, TOPBP1, TP53INP1, TP63, TP73, TPX2* |
| R-HSA-15869 | Metabolism of nucleotides | 1.68 | 0.020 | *ADA, ADK, AMPD3, ATIC, CAD, CDA, CTPS1, DCK, DCTPP1, DTYMK, ENTPD7, GART, GLRX, GMPS, HPRT1, NME1, NT5C3A, NT5E, NUDT15, NUDT5, PAICS, PNP, PPAT, RRM1, RRM2, SAMHD1, TK1, TYMP, TYMS, UMPS, UPP1, XDH* |
| R-HSA-6782315 | tRNA modification in the nucleus and cytosol | 1.67 | 0.021 | *ADAT1, ADAT2, ALKBH8, CTU2, DUS2, FTSJ1, METTL1, NSUN2, OSGEP, PUS1, PUS7, THADA, THG1L, TP53RK, TPRKB, TRIT1, TRMT1, TRMT11, TRMT112, TRMT5, TRMT6, TYW1, TYW3, TYW5, URM1, WDR4* |
| R-HSA-8877330 | RUNX1 and FOXP3 control the development of regulatory T lymphocytes (Tregs) | 1.66 | 0.022 | *CBFB, CTLA4, FOXP3, IFNG, IL2RA, RUNX1* |
| R-HSA-6804116 | TP53 Regulates Transcription of Genes Involved in G1 Cell Cycle Arrest | 1.66 | 0.022 | *CCNA1, CCNA2, CCNE1, CCNE2, CDK2, E2F1, E2F7, E2F8* |
| R-HSA-173623 | Classical antibody-mediated complement activation | 1.66 | 0.023 | *C1QA, C1QB, C1QC* |
| R-HSA-8862803 | Deregulated CDK5 triggers multiple neurodegenerative pathways in Alzheimer's disease models | 1.65 | 0.024 | *APP, BCL2L11, CDC25A, CDC25B, CDC25C, CDK5, FASLG, LMNB1, SOD2, STAT4, YWHAE* |
| R-HSA-8863678 | Neurodegenerative Diseases | 1.65 | 0.024 | *APP, BCL2L11, CDC25A, CDC25B, CDC25C, CDK5, FASLG, LMNB1, SOD2, STAT4, YWHAE* |
| R-HSA-167161 | HIV Transcription Initiation | 1.65 | 0.024 | *ERCC2, ERCC3, GTF2A2, GTF2B, GTF2E1, GTF2E2, GTF2F2, GTF2H1, GTF2H3, GTF2H4, MNAT1, POLR2A, POLR2D, POLR2G, POLR2H, POLR2J, POLR2K, POLR2L, TAF11, TAF12, TAF13, TAF2, TAF4B, TAF6, TBP* |
| R-HSA-211979 | Eicosanoids | -1.80 | 0.026 | *CYP4B1, CYP4F12, CYP4F22, CYP4F3, PTGIS* |
| R-HSA-435354 | Zinc transporters | 1.64 | 0.026 | *SLC30A1, SLC30A6, SLC30A7, SLC39A1, SLC39A10, SLC39A14, SLC39A6,* |

| | | | | |
|---|---|---|---|---|
| | | | | *SLC39A7, SLC39A8* |
| R-HSA-5218859 | Regulated Necrosis | 1.64 | 0.027 | *BIRC2, BIRC3, CASP8, FADD, FASLG, MLKL, RIPK1, TNFRSF10A, TNFRSF10B, TNFSF10* |
| R-HSA-166665 | Terminal pathway of complement | -1.79 | 0.030 | *C7, CLU* |
| R-HSA-448706 | Interleukin-1 processing | 1.62 | 0.031 | *CASP1, IL18, IL1A, IL1B, NFKB1, NFKB2* |
| R-HSA-110312 | Translesion synthesis by REV1 | 1.62 | 0.032 | *MAD2L2, PCNA, RFC1, RFC2, RFC3, RFC4, RFC5, RPA3* |
| R-HSA-6798695 | Neutrophil degranulation | 1.62 | 0.033 | *ABCA13, ACLY, ACTR2, ADAM10, ADAM8, AMPD3, ANPEP, ANXA2, APAF1, ARHGAP9, ARPC5, ATAD3B, ATG7, ATP11A, ATP6AP2, ATP8B4, B2M, B4GALT1, BIN2, BST2, C3AR1, C5AR1, CAB39, CAND1, CAP1, CCT2, CD14, CD177, CD300A, CD44, CD47, CD53, CD58, CD68, CDA, CEP290, CHI3L1, CHIT1, CKAP4, CLEC12A, CLEC5A, CMTM6, CNN2, COPB1, COTL1, CTSA, CTSB, CTSC, CTSD, CTSS, CTSZ, CXCL1, CXCR1, CYBA, CYBB, DDOST, DERA, DIAPH1, DOCK2, DOK3, DSC1, DSN1, DYNC1LI1, DYNLT1, ERP44, FABP5, FAF2, FCER1G, FCGR2A, FOLR3, FPR1, FPR2, FTL, GALNS, GGH, GLA, GLB1, GLIPR1, GM2A, GMFG, GNS, GPR84, GSDMD, GSTP1, HEXB, HLA-B, HLA-C, HMOX2, HSP90AA1, HSP90AB1, HSPA8, IMPDH1, ITGAL, ITGAM, ITGAV, ITGAX, ITGB2, KCNAB2, LAIR1, LCN2, LILRA3, LPCAT1, LRMP, MAN2B1, MIF, MMP8, MMP9, MNDA, MVP, NCKAP1L, NEU1, NFKB1, NIT2, NPC2, NRAS, OLFM4, OLR1, ORM2, PA2G4, PAFAH1B2, PDAP1, PGAM1, PGM2, PKM, PLAC8, PLAU, PLAUR, PNP, PPIA, PRDX4, PSMA2, PSMA5, PSMB1, PSMB7, PSMC2, PSMC3, PSMD1, PSMD11, PSMD12, PSMD13, PSMD14, PSMD2, PSMD3, PTAFR, PTPN6, PTPRC, PTPRJ, PYCARD, PYGB, PYGL, QSOX1, RAB14, RAB18, RAB31, RAC1, RAP1B, RAP2C, RHOF, RHOG, ROCK1, S100A11, S100A12, S100A7, S100A8, S100P, SELL, SERPINA1, SERPINA3, SERPINB3, SIRPA, SIRPB1, SLC11A1, SLC15A4, SLC2A3, STK10, STOM, SURF4, TBC1D10C, TCIRG1, TCN1, TLR2, TNFAIP6, TNFRSF1B, TRAPPC1, TRPM2, TUBB, TUBB4B, TYROBP, VCP, VNN1, XRCC6* |
| R-HSA-141430 | Inactivation of APC/C via direct inhibition of the APC/C complex | 1.61 | 0.034 | *ANAPC1, ANAPC4, ANAPC5, BUB1B, BUB3, CDC20, CDC23, MAD2L1, UBE2C* |

| | | | | |
|---|---|---|---|---|
| R-HSA-5654695 | PI-3K cascade:FGFR2 | -1.77 | 0.034 | *FGF10, FGF16, FGF18, FGF2, FGF6, FGF7, FGF9, FGFR2, GAB1, PIK3R1* |
| R-HSA-9008059 | Interleukin-37 signaling | 1.61 | 0.035 | *CASP1, IL18BP, IL18R1, PTPN12, PTPN13, PTPN2, PTPN4, PTPN6, PTPN7, PTPN9, STAT3, TBK1* |
| R-HSA-5368287 | Mitochondrial translation | 1.60 | 0.038 | *CHCHD1, DAP3, ERAL1, GFM1, GFM2, MRPL1, MRPL11, MRPL13, MRPL14, MRPL15, MRPL16, MRPL17, MRPL18, MRPL21, MRPL22, MRPL23, MRPL27, MRPL28, MRPL3, MRPL32, MRPL37, MRPL42, MRPL47, MRPL48, MRPL49, MRPL50, MRPL52, MRPL9, MRPS10, MRPS12, MRPS15, MRPS17, MRPS18A, MRPS18C, MRPS2, MRPS21, MRPS22, MRPS23, MRPS28, MRPS33, MRPS5, MRRF, MTFMT, MTIF2, PTCD3, TSFM* |
| R-HSA-4090294 | SUMOylation of intracellular receptors | -1.76 | 0.040 | *AR, NR3C2, NR4A2, NR5A2, PGR, PIAS2, PPARA, PPARG, RORA, THRA, THRB* |
| R-HSA-5423646 | Aflatoxin activation and detoxification | -1.75 | 0.040 | *AKR7A2, CYP3A4, CYP3A5, GGT6, GGT7, MGST1, MGST2, MGST3* |
| R-HSA-912526 | Interleukin receptor SHC signaling | 1.59 | 0.041 | *CSF2, CSF2RA, IL2RA, IL2RB, IL2RG, JAK2, JAK3, PIK3CD, SHC1* |
| R-HSA-8984722 | Interleukin-35 Signalling | 1.59 | 0.041 | *IL12RB2, JAK2, STAT1, STAT4* |
| R-HSA-69231 | Cyclin D associated events in G1 | 1.58 | 0.043 | *CCND2, CDK4, CDK6, CDKN1A, CDKN2A, CKS1B, CUL1, E2F1, E2F3, E2F4, E2F5, JAK2, LYN, MNAT1, RB1, RBL1, SKP2, SRC, TFDP1* |
| R-HSA-420029 | Tight junction interactions | -1.75 | 0.043 | *CLDN1, CLDN10, CLDN11, CLDN17, CLDN3, CLDN4, CLDN5, CLDN7, CLDN8, MPP5, PARD3, PARD6A* |
| R-HSA-3781860 | Diseases associated with N-glycosylation of proteins | 1.58 | 0.045 | *ALG1, ALG13, ALG2, ALG3, ALG6, ALG8, B4GALT1, DPAGT1, MAN1B1, MGAT2, MOGS, MPDU1* |
| R-HSA-379724 | tRNA Aminoacylation | 1.57 | 0.046 | *AARS2, AIMP1, AIMP2, DARS2, EARS2, EEF1E1, FARS2, FARSA, FARSB, HARS2, LARS2, MARS2, NARS2, PPA1, RARS2, YARS2* |
| R-HSA-2995383 | Initiation of Nuclear Envelope Reformation | 1.57 | 0.048 | *ANKLE2, LMNB1, TMPO, VRK1, VRK2* |

| | | | | |
|---|---|---|---|---|
| R-HSA-2995410 | Nuclear Envelope Reassembly | 1.57 | 0.048 | *ANKLE2, LMNB1, TMPO, VRK1, VRK2* |
| R-HSA-5669034 | TNFs bind their physiological receptors | 1.57 | 0.048 | *CD27, EDARADD, FASLG, TNFRSF17, TNFRSF9, TNFSF13B, TNFSF18, TNFSF4, TNFSF9* |

Table S8 - Magenta module genes' categories identified using DGIdb. Gene names in bold indicate hubs.

| Gene | Categories |
|---|---|
| *ABCA12* | Druggable Genome |
| *ACE2* | Druggable Genome, Neutral Zinc Metallopeptidase |
| *ADIPOR1* | Kinase |
| *AKR1B10* | Druggable Genome |
| *ALDH3B2* | Druggable Genome |
| *ALOX12B* | Clinically Actionable, Druggable Genome |
| *ALOXE3* | Druggable Genome, Nuclear Hormone Receptor |
| *AOC1* | Druggable Genome, Ion Channel |
| *APH1A* | Clinically Actionable, Druggable Genome |
| *ASPRV1* | Druggable Genome |
| *ATF6* | Transcription Factor |
| *AVPI1* | Kinase |
| *BPIFC* | Druggable Genome |
| *BRAT1* | Kinase |

| | |
|---|---|
| *CARD6* | Transcription Factor, Kinase |
| *CDA* | Druggable Genome |
| *CDK7* | Druggable Genome, Transcription Factor, Serine Threonine Kinase |
| *CIB1* | Kinase |
| *CITED2* | Transcription Factor |
| *CLDN17* | Ion Channel |
| *CPS1* | Clinically Actionable, Druggable Genome |
| *CST6* | Druggable Genome |
| *CXCR2* | Druggable Genome, G Protein Coupled Receptor |
| *CYP4F22* | Druggable Genome |
| *DEFB1* | Druggable Genome |
| *DHRS1* | Druggable Genome |
| *DHRS9* | Druggable Genome |
| *DLX5* | Transcription Factor |
| *DMKN* | Druggable Genome |
| ***DSC1*** | Hub |
| ***DSG1*** | Hub |
| ***DSP*** | Hub |
| *DUOX1* | Druggable Genome |

| | |
|---|---|
| *DUSP14* | Druggable Genome, Kinase |
| *DUSP18* | Druggable Genome, Kinase |
| *EPHA4* | Druggable Genome, Tyrosine Kinase |
| *ETHE1* | Transcription Factor |
| *FAM20B* | Kinase |
| *FIBCD1* | Druggable Genome |
| ***FLG*** | Hub |
| *FLVCR2* | Druggable Genome |
| *GADD45A* | Kinase |
| *GFOD2* | Druggable Genome |
| *GJA1* | Ion Channel |
| *GJB2* | Ion Channel |
| *GJB6* | Ion Channel |
| *HMOX1* | Druggable Genome |
| *HPSE* | Druggable Genome |
| *HSPB1* | Druggable Genome |
| *HYAL1* | Druggable Genome |
| *IDE* | Druggable Genome, Neutral Zinc Metallopeptidase |
| *IDS* | Druggable Genome |

| | |
|---|---|
| *IL22RA1* | Kinase |
| *IL36G* | Druggable Genome, Kinase |
| *IL36RN* | Druggable Genome |
| *IRAK2* | Druggable Genome, Serine Threonine Kinase |
| ***IVL*** | Hub |
| ***JUP*** | Hub |
| *KCNK6* | Druggable Genome, Ion Channel |
| *KLHL18* | Druggable Genome |
| *KLK10* | Druggable Genome |
| *KLK14* | Druggable Genome |
| *KLK6* | Druggable Genome |
| *KLK7* | Druggable Genome |
| *KLK8* | Druggable Genome |
| *KRT1* | Kinase |
| *KRTDAP* | Druggable Genome |
| *LAD1* | Druggable Genome |
| *LCN2* | Druggable Genome |
| *LYPD5* | Druggable Genome |
| *LZTR1* | Clinically Actionable |

| | |
|---|---|
| *MAEA* | Transcription Factor |
| *MAFF* | Transcription Factor |
| *MBOAT2* | Druggable Genome |
| *MED8* | Transcription Factor |
| *MOK* | Druggable Genome, Transcription Factor, Serine Threonine Kinase |
| *MPZL3* | Druggable Genome |
| *MXD1* | Transcription Factor |
| *NLRX1* | Kinase |
| *NME4* | Kinase |
| *NRBP1* | Druggable Genome, Serine Threonine Kinase, Nuclear Hormone Receptor |
| *PDPK1* | Clinically Actionable, Druggable Genome, Serine Threonine Kinase |
| *PELI1* | Kinase |
| *PFKFB2* | Kinase |
| *PGBD1* | Druggable Genome |
| *PGLYRP4* | Druggable Genome |
| *PHF3* | Transcription Factor |
| *PHKA2* | Kinase |
| ***PI3*** | Hub, Druggable Genome |
| *PIP5K1C* | Druggable Genome, Kinase |

| | |
|---|---|
| ***PKP1*** | Hub |
| *PNLIPRP3* | Druggable Genome |
| *QSOX1* | Druggable Genome |
| *RAB9A* | Druggable Genome |
| *RAPGEF4* | Druggable Genome |
| *RDH12* | Druggable Genome |
| *RNASE7* | Druggable Genome |
| *RNF39* | Druggable Genome |
| *ROCK2* | Druggable Genome, Serine Threonine Kinase |
| ***RPTN*** | Hub, Druggable Genome |
| *RTN4R* | Druggable Genome |
| *S100A12* | Druggable Genome, Kinase |
| *S100A7* | Druggable Genome |
| *S100A9* | Druggable Genome, Kinase |
| *SDR9C7* | Druggable Genome |
| *SERPINB3* | Druggable Genome, Kinase |
| *SERPINB4* | Druggable Genome |
| *SERPINB7* | Druggable Genome |
| *SLC15A1* | Druggable Genome |

| | |
|---|---|
| *SLC25A29* | Druggable Genome |
| *SLC27A1* | Druggable Genome |
| *SLC5A1* | Druggable Genome |
| *SLC5A10* | Druggable Genome |
| *SLPI* | Druggable Genome |
| *SPAG9* | Kinase |
| *SPINK6* | Druggable Genome |
| ***SPRR1B*** | Hub |
| ***SPRR2G*** | Hub |
| *STRIP1* | Kinase |
| *STYXL1* | Druggable Genome |
| *TAB1* | Druggable Genome, Kinase |
| *TAF8* | Transcription Factor |
| *TCF7L1* | Transcription Factor |
| *TGM5* | Druggable Genome |
| *TMPRSS13* | Druggable Genome |
| *TP53INP2* | Kinase |
| *TPRA1* | G Protein Coupled Receptor |
| *TRIM45* | Kinase |

| | |
|---|---|
| *TSG101* | Druggable Genome |
| *TUBB2A* | Druggable Genome |
| *UBQLN2* | G Protein Coupled Receptor |
| *ULBP2* | Druggable Genome |
| *VASN* | Druggable Genome |
| *ZMYM3* | Clinically Actionable |
| *ZSCAN31* | Transcription Factor |

Table S9 - Drug interactions identified based on magenta module's genes. Gene names in bold represent hubs.

| Gene | Drug | Interaction type | Tested in OSCC | Tested in other neoplasms | Approved in OSCC | Approved in other neoplasms |
|---|---|---|---|---|---|---|
| *CDSN* | Carboplatin | | No | Yes | No | Yes |
| *CDSN* | Gemcitabine | | No | Yes | No | Yes |
| *S100A12* | Methotrexate | | No | Yes | No | Yes |
| ***PI3*** | Progesterone | | No | Yes | No | Yes |
| ***IVL*** | Adalimumab | | No | Yes | No | Yes |
| ***IVL*** | Infliximab | | No | Yes | No | Yes |
| *GJB6* | Carbenoxolone | inhibitor | No | No | No | No |
| *CDA* | Cytarabine | | No | Yes | No | Yes |

| | | | | | | |
|---|---|---|---|---|---|---|
| *CDA* | Capecitabine | | No | Yes | No | Yes |
| *CDA* | Gemcitabine | | No | Yes | No | Yes |
| *CDA* | Azacitidine | | No | Yes | No | No |
| *SLC5A1* | Empagliflozin | inhibitor | No | Yes | No | Yes |
| *SLC5A1* | Ertugliflozin | inhibitor | No | No | No | No |
| *SLC5A1* | Streptozocin | | No | No | No | No |
| *SLC5A1* | Dapagliflozin | inhibitor | No | Yes | No | Yes |
| *SLC5A1* | Canagliflozin | inhibitor | No | Yes | No | Yes |
| *EPHA4* | Vandetanib | inhibitor | No | Yes | No | Yes |
| *HMOX1* | Sunitinib | | Yes | Yes | No | No |
| *HMOX1* | Sorafenib | | No | Yes | No | Yes |
| *HMOX1* | Aspirin | | No | No | No | No |
| *GJA1* | Carvedilol | other/unknown | No | Yes | No | Yes |
| *GJA1* | Labetalol | | No | Yes | No | Yes |
| *GJA1* | Bleomycin | | No | Yes | No | Yes |
| *GJA1* | Propylthiouracil | | No | Yes | No | Yes |
| *GJA1* | Carbenoxolone | inhibitor | No | No | No | No |
| *GJA1* | Atenolol | | No | Yes | No | Yes |
| *HPSE* | Metergoline | | No | No | No | No |

| | | | | | | |
|---|---|---|---|---|---|---|
| *HPSE* | Thrombin | | No | No | No | No |
| *HPSE* | Labetalol | | No | Yes | No | Yes |
| *TUBB2A* | Vinflunine | inhibitor | No | No | No | No |
| *TUBB2A* | Vinorelbine | | No | Yes | No | Yes |
| *TUBB2A* | Vincristine Sulfate | inhibitor | No | No | No | No |
| *TUBB2A* | Brentuximab Vedotin | inhibitor | No | Yes | No | Yes |
| *TUBB2A* | Vorinostat | | No | Yes | No | Yes |
| *TUBB2A* | Colchicine | inhibitor | No | Yes | No | Yes |
| *TUBB2A* | Podofilox | | No | Yes | No | Yes |
| *TUBB2A* | Vinblastine | | No | Yes | No | Yes |
| *TUBB2A* | Docetaxel | inhibitor | No | Yes | No | Yes |
| *TUBB2A* | Vinorelbine Tartrate | inhibitor | No | No | No | No |
| *TUBB2A* | Ixabepilone | inhibitor | No | Yes | No | Yes |
| *TUBB2A* | Vincristine | | No | Yes | No | Yes |
| *TUBB2A* | Vinblastine Sulfate | inhibitor | No | No | No | No |
| *TUBB2A* | Eribulin Mesylate | inhibitor | No | No | No | No |
| *TUBB2A* | Paclitaxel | inhibitor | No | Yes | No | Yes |
| *TUBB2A* | Trastuzumab Emtansine | inhibitor | No | Yes | No | Yes |
| *TUBB2A* | Cabazitaxel | inhibitor | No | Yes | No | Yes |

| | | | | | | |
|---|---|---|---|---|---|---|
| *CXCR2* | Clotrimazole | | No | Yes | No | Yes |
| *CXCR2* | Acetylcysteine | | No | Yes | No | Yes |
| *CXCR2* | Ibuprofen | | No | Yes | No | Yes |
| *GJB2* | Carbenoxolone | inhibitor | No | No | No | No |
| *IDE* | Bacitracin | inhibitor | No | Yes | No | Yes |
| *IDE* | Biotin | | No | Yes | No | Yes |
| *ACE2* | Captopril | | No | Yes | No | Yes |
| *ACE2* | Lisinopril | inhibitor | No | Yes | No | Yes |
| *RAB9A* | Hydralazine | | No | Yes | No | Yes |
| *RAB9A* | Phenelzine | | No | Yes | No | Yes |
| *RAB9A* | Leflunomide | | No | Yes | No | Yes |
| *RAB9A* | Rabeprazole Sodium | | No | No | No | No |
| *RAB9A* | Lansoprazole | | No | Yes | No | Yes |
| *RAB9A* | Riluzole | | No | Yes | No | Yes |
| *RAB9A* | Niclosamide | | No | Yes | No | Yes |
| *RAB9A* | Amlexanox | | No | Yes | No | Yes |
| *SRPRB* | Alcohol | | No | No | No | No |
| *OPLAH* | Melphalan | | No | Yes | No | Yes |
| *TSG101* | Daunorubicin Hydrochloride | | No | No | No | No |

| | | | | | | |
|---|---|---|---|---|---|---|
| *TSG101* | Dipyridamole | | No | Yes | No | Yes |
| *TSG101* | Topotecan Hydrochloride | | No | No | No | No |
| *CPS1* | Carglumic Acid | positive modulator | No | No | No | No |
| *CPS1* | Methionine | | No | No | No | No |
| *GADD45A* | Cisplatin | | Yes | Yes | No | No |
| *GADD45A* | Doxorubicin | | No | Yes | No | Yes |
| ***DSP*** | Enalapril | | No | Yes | No | Yes |
| *CDK7* | Pazopanib | | No | Yes | No | Yes |
| *RAPGEF4* | Acriflavine | | No | No | No | No |
| *RAPGEF4* | Nabumetone | | No | Yes | No | Yes |
| *RAPGEF4* | Idarubicin Hydrochloride | | No | No | No | No |
| *RAPGEF4* | Clotrimazole | | No | Yes | No | Yes |
| *RAPGEF4* | Acitretin | | No | Yes | No | Yes |
| *RAPGEF4* | Lanatoside C | | No | No | No | No |
| *RAPGEF4* | Prochlorperazine | | No | Yes | No | Yes |
| *RAPGEF4* | Hexachlorophene | | No | No | No | No |
| *RAPGEF4* | Dipyridamole | | No | Yes | No | Yes |
| *RAPGEF4* | Benzbromarone | | No | No | No | No |
| *IDS* | Idursulfase | | No | Yes | No | Yes |

| | | | | | | |
|---|---|---|---|---|---|---|
| *ROCK2* | Vandetanib | | No | Yes | No | Yes |
| *ROCK2* | Palbociclib | | No | Yes | No | Yes |
| *ROCK2* | Progesterone | | No | Yes | No | Yes |
| *ROCK2* | Netarsudil | inhibitor | No | No | No | No |
| *ROCK2* | Thrombin | | No | No | No | No |
| *HSPB1* | Urea | | No | Yes | No | Yes |
| *HSPB1* | Vasopressin | | No | Yes | No | Yes |
| *HSPB1* | Omeprazole | | No | Yes | No | Yes |
| *HSPB1* | Triamterene | | No | Yes | No | Yes |
| *HSPB1* | Mometasone Furoate | | No | No | No | No |
| *HSPB1* | Albendazole | | No | Yes | No | Yes |
| *SLC15A1* | Fluvastatin | | No | Yes | No | Yes |
| *SLC15A1* | Acyclovir | | No | No | No | No |
| *PIP5K1C* | Alcohol | | No | No | No | No |
| *SIRT5* | Cefixime | | No | Yes | No | Yes |

Table S10 - Drug-induced gene perturbations identified for overlapping DEGs among OLP and OSCC using the L1000-CDS2 tool.

| Rank | score | Perturbation | Cell-line | Dose | Time | Class (DrugBank) | Class (Other databases) | Tested in OSCC | Tested in other tumors | Approved in OSCC | Approved in other tumors |
|---|---|---|---|---|---|---|---|---|---|---|---|
| 1 | 0.2593 | BRD-K70161581 | PC3 | 10.0um | 24h | Unknown | | No | No | No | No |
| 2 | 0.2222 | BRD-K81473043 | PC3 | 0.35um | 24h | Unknown | | No | No | No | No |

| | | | | | | | | | | | |
|---|---|---|---|---|---|---|---|---|---|---|---|
| 3 | 0.2222 | YM-201636 | BT20 | 10um | 24h | Unknown | | No | No | No | No |
| 4 | 0.2222 | GSK-1059615 | HCC515 | 3.33um | 24h | Not available | | No | No | No | No |
| 5 | 0.1852 | geldanamycin | PC3 | 10.0um | 24h | Tyrosine kinase inhibitor, protease inhibitor, cysteine proteinase inhibitor, antibiotic, antineoplastic | | No | No | No | No |
| 6 | 0.1852 | Kenpaullone | HA1E | 10.0um | 24h | Unknown | | No | No | No | No |
| 7 | 0.1852 | bongkrekic acid | HA1E | 10.0um | 6h | Unknown | Apoptosis inhibitor (PubChem) | No | No | No | No |
| 8 | 0.1852 | NALTREXONE HYDROCHLORIDE | HCC515 | 10.0um | 6h | Opiate antagonist, alcohol deterrent, central nervous system agent | | No | No | No | No |
| 9 | 0.1852 | Naltrexone hydrochloride | A549 | 10.0um | 24h | Opiate antagonist, alcohol deterrent, central nervous system agent | | No | No | No | No |
| 10 | 0.1852 | KU 0060648 trihydrochloride | HA1E | 10.0um | 24h | Unknown | DNA-dependent protein kinase inhibitor ( https://www.medchemexpress.com/ku-0060648.html ) | No | No | No | No |
| 11 | 0.1852 | BRD-K81473043 | HT29 | 0.35um | 24h | Quinone, protein-serine-threonine-kinase inhibitor, HSP90 antagonist | | No | No | No | No |
| 12 | 0.1852 | vorinostat | NCIH1836 | 11.1um | 6h | Histone deacetylase inhibitor, immunosuppressive agent | | No | Yes | No | Yes |
| 13 | 0.1852 | HY-10181 | HA1E | 10.0um | 6h | Tyrosine kinase inhibitor | | No | No | No | No |
| 14 | 0.1852 | BRD-K19220233 | PC3 | 10.0um | 24h | Unknown | | No | No | No | No |
| 15 | 0.1852 | EI-156 | PC3 | 10.0um | 24h | Unknown | | No | No | No | No |
| 16 | 0.1852 | 10012682 | PC3 | 10.0um | 24h | Unknown | | No | No | No | No |

| | | | | | | | | | | | |
|---|---|---|---|---|---|---|---|---|---|---|---|
| 17 | 0.1852 | BRD-K17953061 | PC3 | 10.0um | 24h | Enzyme inhibitor | | No | No | No | No |
| 18 | 0.1852 | Olmesartan | PC3 | 10.0um | 6h | Angiotensin II receptor blocker | | No | No | No | Yes |
| 19 | 0.1852 | CGP-60474 | HEPG2 | 0.37um | 24h | Unknown | | No | No | No | No |
| 20 | 0.1852 | GDC-0980 | MCF10A | 1.11um | 3h | PI3K inhibitor | | No | No | No | No |
| 21 | 0.1852 | torin-2 | MCF10A | 1.11um | 3h | | mTOR inhibitor (PubChem) https://pubchem.ncbi.nlm.nih.gov/compound/Torin-2 | No | No | No | No |
| 22 | 0.1852 | geldanamycin | MCF7 | 1.11um | 24h | Cysteine proteinase inhibitor, tyrosine kinase inhibitor | | No | No | No | No |
| 23 | 0.1852 | pelitinib | SKBR3 | 0.37um | 24h | ErbB receptor, antagonist and inhibitor | | No | No | No | No |
| 24 | 0.1852 | GDC-0980 | SKBR3 | 1.11um | 24h | PI3K inhibitor | | No | No | No | No |
| 25 | 0.1852 | mitoxantrone | SKBR3 | 1.11um | 3h | Enzyme inhibitor, immunosuppressive agent, antineoplastic | | No | Yes | No | Yes |
| 26 | 0.1852 | fostamatinib | BT20 | 10um | 24h | Tyrosine kinase inhibitor, cytochrome P-450 enzyme inhibitor | | No | No | No | No |
| 27 | 0.1852 | KIN001-043 | BT20 | 10um | 24h | Unknown | | No | No | No | No |
| 28 | 0.1852 | dasatinib | HME1 | 3.33um | 3h | Tyrosine kinase inhibitor | | No | Yes | No | Yes |
| 29 | 0.1852 | GSK-1059615 | HME1 | 10um | 3h | Not available | | No | No | No | No |
| 30 | 0.1852 | afatinib | SKBR3 | 0.12um | 24h | ErbB family blocker, protein kinase inhibitor | | No | Yes | No | Yes |
| 31 | 0.1852 | CGP-60474 | A375 | 0.04um | 24h | Unknown | | No | No | No | No |
| 32 | 0.1852 | GSK-1059615 | HCC515 | 3.33um | 24h | Not available | | No | No | No | No |
| 33 | 0.1852 | INK-128 | HEPG2 | 0.04um | 24h | Unknown | | No | No | No | No |

| | | | | | | | | | | | |
|---|---|---|---|---|---|---|---|---|---|---|---|
| 34 | 0.1481 | COSMOSIIN | HCC515 | 10.0um | 24h | Benzopyran, flavonoid | | No | No | No | No |
| 35 | 0.1481 | PPT | HA1E | 10.0um | 6h | | Estrogen receptor agonist (PubChem) https://pubchem.ncbi.nlm.nih.gov/compound/Propyl-pyrazole-triol | No | No | No | No |
| 36 | 0.1481 | ETHYL-beta-CARBOLINE-3-CARBOXYLATE | HA1E | 10.0um | 6h | Unknown | | No | No | No | No |
| 37 | 0.1481 | Phenformin hydrochloride | VCAP | 10.0um | 24h | Hypoglycemic, OCT1 and OCT2 inhibitor | | No | No | No | No |
| 38 | 0.1481 | TC 2559 difumarate | VCAP | 10.0um | 24h | Unknown | | No | No | No | No |
| 39 | 0.1481 | CGP 57380 | HCC515 | 10.0um | 24h | Unknown | MAPK inhibitor | No | No | No | No |
| 40 | 0.1481 | DAUNORUBICIN | PC3 | 10.0um | 24h | Anthracycline topoisomerase inhibitor | | No | Yes | No | Yes |
| 41 | 0.1481 | METHOXSALEN | A375 | 10.0um | 24h | Cytochrome P-450 CYP2A6 inhibitor | | No | Yes | No | Yes |
| 42 | 0.1481 | geldanamycin | HA1E | 10.0um | 24h | Tyrosine kinase inhibitor, protease inhibitor, cysteine proteinase inhibitor, antibiotic, antineoplastic | | No | No | No | No |
| 43 | 0.1481 | N-FORMYLMETHIONYLALANINE | HA1E | 10.0um | 6h | Unknown | | No | No | No | No |
| 44 | 0.1481 | vorinostat | HA1E | 10.0um | 6h | Histone deacetylase inhibitor, immunosuppressive agent | | No | Yes | No | Yes |
| 45 | 0.1481 | COT-10b | A673 | 44.4um | 6h | Unknown | | No | No | No | No |
| 46 | 0.1481 | QUINACRINE HYDROCHLORIDE | EFO27 | 10.0um | 6h | Enzyme inhibitor, antimalarial agent | | No | No | No | No |
| 47 | 0.1481 | BMS-536924 | HA1E | 11.1um | 24h | Unknown | | No | No | No | No |

| | | | | | | | | | | | |
|---|---|---|---|---|---|---|---|---|---|---|---|
| 48 | 0.1481 | BRD-K41859756 | HA1E | 10.0um | 24h | | HSP90 inhibitor (PubChem) https://pubchem.ncbi.nlm.nih.gov/compound/135539077 | No | No | No | No |
| 49 | 0.1481 | PI 103 hydrochloride | HA1E | 11.1um | 24h | Unknown | | No | No | No | No |
| 50 | 0.1481 | BRD-A62809825 | HA1E | 10.0um | 6h | Unknown | | No | No | No | No |
